# Development of ketobenzothiazole-based peptidomimetic TMPRSS13 inhibitors with low nanomolar potency

**DOI:** 10.1101/2024.08.28.609965

**Authors:** Alexandre Joushomme, Antoine Désilets, William Champagne, Malihe Hassanzadeh, Gabriel Lemieux, Alice Gravel-Trudeau, Matthieu Lepage, Sabrina Lafrenière, Ulrike Froehlich, Karin List, Pierre-Luc Boudreault, Richard Leduc

**Affiliations:** Department of Pharmacology-Physiology, Faculty of Medicine and Health Sciences, Université de Sherbrooke, Sherbrooke, Québec, Canada; Department of Pharmacology, Wayne State University, Detroit, MI 48202, USA

**Keywords:** Peptidomimetic, TMPRSS13, protease inhibitor, compound screening, SARS-CoV-2

## Abstract

TMPRSS13, a member of the Type II Transmembrane Serine Proteases (TTSP) family, is involved in cancer progression and in cell entry of respiratory viruses. To date, no inhibitors have been specifically developed toward this protease. In this study, a chemical library of 65 ketobenzothiazole-based peptidomimetic molecules was screened against a proteolytically active form of recombinant TMPRSS13 to identify novel inhibitors. Following an initial round of screening, subsequent synthesis of additional derivatives supported by molecular modelling, uncovered important molecular determinants involved in TMPRSS13 inhibition. One inhibitor, N-0430, achieved low nanomolar affinity towards TMPRSS13 activity in a cellular context. Using a SARS-CoV-2 pseudovirus cell entry model, we further show the ability of N-0430 to block TMPRSS13-dependent entry of the pseudovirus. The identified peptidomimetic inhibitors and the molecular insights of their potency gained from this study will aid in the development of specific TMPRSS13 inhibitors.

## Introduction

Proteolysis is a finely regulated physiological process and protease malfunction can lead to a wide range of pathologies^1^. Such deregulation of one particular family of proteolytic enzymes, the Type II Transmembrane Serine Proteases (TTSP) family, is associated with a variety of conditions such as digestive malfunctions, iron overload, cancer, skin disorders and others^2,3^. TTSPs are constituted of an N-terminal cytoplasmic tail followed by a transmembrane domain, an intermediate region containing different ancillary domains, and a C-terminal extracellular serine protease domain exposed at the cell surface^4–6^. They are synthesized as single-chain inactive zymogen, requiring proteolytic activation at specific arginine or lysine residues preceding the catalytic domain^5^. Following activation, the catalytic domain remains bound to ancillary domains via a disulfide bridge^6^. Moreover, cell-surface shedding of TTSP ectodomains allows the release of an active form of the enzyme into the pericellular space^7,8^.

TMPRSS13, a TTSP highly expressed in the skin and to a lesser extent in the proximal digestive tract, contributes to the formation of the epidermal barrier^9^. It is also expressed in lungs, pancreas, prostate, colon, breast and various tissues of the digestive system^6,10,11^. Recent work has revealed its involvement in the progression of both breast and colorectal cancer, where TMPRSS13 silencing augments apoptosis, reduces cell invasion and increases sensitivity to chemotherapeutic agents^11,12^. Importantly, TMPRSS13 deficiency was associated to decreasing tumor growth *in vivo*^11^. These results position TMPRSS13 as an interesting therapeutic target in cancer treatment, where its inhibition could enhance the efficacy of commonly used chemotherapies. Additionally, TMPRSS13 is also known as a viral-priming protease by its ability to activate type I surface glycoproteins of respiratory viruses through proteolytic cleavage, an essential step for their entry into host cells. Hence, it was demonstrated that TMPRSS13 cleaves and primes SARS-CoV^13^, MERS-CoV^13^, SARS-CoV-2^14–16^, influenza A virus^17^ as well as highly pathogenic avian influenza virus (HPAIV) Spike or hemagglutinin glycoproteins known to harbor multi-basic cleavage sites^18^. Recently, TMPRSS13 was also identified as playing a key role in the onset of Swine Acute Diarrhea Syndrome (SADS-CoV)^19^. This virus originates in bats^20^, thus presenting a high risk of transmission between species. The implications of TMPRSS13 in various respiratory viruses posing pandemic threats makes this protease an important target to consider in the context of antiviral preparedness.

Currently, TMPRSS13 inhibition has been reported with broad-acting serine protease inhibitors such as aprotinin^6,18,21^, benzamidine^6^, Bowman-Birk trypsin inhibitors^6,18^, camostat^19^, nafamostat^16^ and decanoyl-RVKR-CMK, a proprotein convertase inhibitor containing an irreversible chloromethyl ketone warhead^18,22^. However, no studies aimed at developing inhibitors directed specifically against TMPRSS13 have been reported. We have previously achieved potent inhibition of different TTSPs (matriptase^23,24^, TMPRSS6^25–27^and TMPRSS2^28^) using peptidomimetic compounds harboring a ketobenzothiazole (kbt) warhead, able to form a covalent but reversible bond between its ketone and the serine of the protease’s catalytic triad, thereby acting as serine trap. Inhibition of viral serine proteases has previously been approved by the FDA in the treatment of hepatitis C virus (HCV) ^29,30^ with Telaprevir and Boceprevir, two protease inhibitors having a mechanism of inhibition similar to the ketobenzothiazole warhead. As therapies targeting host factors were also proven to be successful through FDA approbation for HCV and HIV^31–33^, this places the development of TTSP inhibitors in an attractive position, as evidenced by the recent surge in their interest as potential therapeutic targets^25,28,34–37^. Here, the production of active recombinant TMPRSS13 has allowed the screening of a library of peptidomimetic compounds to identify high-affinity inhibitors. The most promising compound was then optimized through the synthesis of derivatives and important pharmacophores for TMPRSS13 inhibition were identified. *In cellulo*, a proof-of-concept of the most potent compound as a potential antiviral is also presented. Overall, this study reports the development of first generation potent TMPRSS13 inhibitors with nanomolar activity both *in vitro* and *in cellulo*, which, following further optimization, could be potentially exploited in different therapeutic contexts.

## Materials and methods

### Cell culture

Simian kidney Vero C1008 (Vero E6, ATCC, CRL-1587) cells were cultured in Eagle’s Minimum Essential Medium (EMEM) (Wisent, 320-005-CL) containing 10% fetal bovine serum (FBS; 080-150 Wisent), and 1 % antibiotics mix (Penicillin Streptomycin, Wisent, 450-202-EL). Human 293T (ATCC, CRL-3216,) and HEK293SL (gift from Stephane Laporte, McGill, Canada), a 293-cell line subclone selected for enhanced adherence^38^, were maintained in Dulbecco’s modified Eagle’s medium high Glucose (DMEM) (Wisent, 319-005-CL) complemented with 10 % FBS and 1 % antibiotics mix. Expi293F cells (Thermo Fisher Scientific, A14527) were cultured in Expi293^TM^ expression medium (Thermo Fisher Scientific, A1435101). All cell lines were cultured in a humidified incubator at 37°C with 5% CO_2_ in air atmosphere, except Expi293F cells with 8% CO_2_.

### DNA constructs

In this study, a variant of TMPRSS13 isoform 1 (GenBank: NM_001077263.3, 567 amino acids), in which one of the five-amino-acid cytoplasmic tandem repeat is missing, was used as the wild type (WT) (GenBank: BC114928.1, 562 amino acids). The TMPRSS13 coding sequence was subcloned in pcDNA 3.4 TOPO vector using the pcDNA 3.4 TOPO TA cloning kit (Thermo Fisher Scientific, A14697) to obtain the TMPRSS13-WT construct^39^. TMPRSS13-AFAAS-6His was obtained by substitution of amino acids R558, R560 and K561 and addition of a 6xHis tag at the c-terminal using the QuikChange Lightning site-directed mutagenesis kit (Agilent, 210518).

### Expression of TMPRSS13 in HEK293SL cells, activity measurement and immunodetection

HEK239SL cells were transfected with 1 µg of plasmids (empty vector, TMPRSS13-WT or TMPRSS13-AFAAS-6His) using Lipofectamine 3000 (Thermo Fisher Scientific, L3000001) and Opti-MEM (Thermo Fisher Scientific, 31985062), in 6-well plates at a density of 500 000 cells per well. 24h after transfection, cells were washed with PBS (Wisent, 311-425-CL) and HCell-100 media was added (Wisent, 001-035-CL) for 24h before samples were collected. 1,5 mL of cell media was collected before cells were lysed at 4 °C using a solution containing 1 % Triton, 50 mM Tris pH 7.4, 150 mM NaCl and 5 mM EDTA, supplemented with protease inhibitor cocktail (Millipore Sigma, 11697498001). Lysate was cleared by centrifugation at 15 000 g, 4 °C, 15 minutes, and protein concentration was calculated using a protein assay dye (Bio-Rad Laboratories, 5000006). Cell media was cleared by centrifugation at 15 000 g, 4 °C, 15 minutes, and proteolytic activity was measured by monitoring Boc-RVRR-AMC (Bachem, 4018735.0025) cleavage using BioTek Synergy HTX Multimode Reader (Agilent Technologies). Cell media was concentrated for western blot using Amicon Ultra centrifugal filters 10 K (Millipore, UFC501096). Equal quantities of proteins from lysates and 30 µl of concentrated media were diluted in Laemmli sample buffer (Bio-Rad Laboratories, 1610747) with 2.5 % β-mercaptoethanol and separated on 4-15 % SDS-page precast gels (Bio-Rad Laboratories, 4561084). Proteins were transferred on nitrocellulose membranes (iBlot 2 transfer stacks, Thermo Fisher Scientific, IB23001). Primary antibodies used were rabbit anti-TMPRSS13 (1:2500 ; Thermo Fisher scientific, PA5-30935), mouse HRP linked anti-β-actin (1:1000 ; Cell Signaling technology, 12262S), and rabbit anti-Histidine (1:1000 ; Cell signaling technology, 2365S). When necessary, HRP-linked goat anti-rabbit as a secondary antibody (1:1000; Cell signaling technology, 7074) was used. Revelation was done using Clarity Western ECL substrate (Bio-Rad Laboratories, 1705061) on an imaging system (Vilber). Membranes were stripped with Restore PLUS Western Blot Stripping Buffer (Thermo Fisher Scientific, 46430) for 15 minutes at room temperature before wash and addition of new primary antibodies.

### Production of recombinant TMPRSS13 for compound screening

TMPRSS13-AFAAS-6His was expressed using the Expi293 Expression System Kit (Thermo Fisher Scientific, A14635) according to the manufacturer’s instructions and purified by IMAC chromatography. Briefly, Expi293F cells were transfected for 48h with TMPRSS13-AFAAS-6His construct and media containing the shed recombinant protease was injected on a HisTrap Excel column (Cytiva, 17371206) using an ÄKTA Start protein purification system (Cytiva). Bound proteins were then eluted (20 mM sodium phosphate pH 7,4, 500 mM NaCl, 500 mM Imidazole) and TMPRSS13 containing fractions were dialyse 16h at 4°C in storage buffer (20 mM sodium phosphate pH 7,4, 150 mM NaCl, 10% Glycerol). Proteins were then concentrated using Amicon Ultra centrifugal filter 10 kDa (Millipore Sigma, UFC8010) and kept at −80°C.

### Mass Spectrometry

Protein preparation content was analysed by Université de Sherbrooke Proteomic Platform to identify TMPRSS13 cleavage fragments. Briefly, and as previously described^40^, mass spectrometry analysis of chymotrypsin peptides was performed using an Orbitrap QExactive mass spectrometer (Thermo Fisher Scientific). Data analysis was performed with the MaxQuant software.

### Synthesis of compounds

All compounds were synthetized as described in the general procedure method (Scheme 1). The elongation of the peptide-based inhibitors was carried out as follows: (a) The first Fmoc-protected amino acid was loaded on the resin using DIPEA in DCM. (b) The Fmoc protecting group was removed with 20% piperidine in DMF. (c) The next Fmoc-protected amino acid was coupled using HATU and DIPEA in DMF. Steps (b) and (c) were repeated sequentially until the desired peptide sequence was completed. (d) The peptide was cleaved from the resin using 20% HFIP in DCM at room temperature while keeping the protecting group. The warhead part was coupled to the tripeptides. (a) Amidation of the peptide was carried out with NH_2_-Arg(OH)bt in the presence of HATU and DIPEA in DMF. Boc-Arg(OH)bt commercially available (Piramal Group, Mumbai, India). The Boc protecting group of Boc-Arg(OH)bt was removed using a solution of 20% TFA in DCM). (b) Oxidation of the secondary alcohol to the ketone was performed using DMP in DCM. (c) Final deprotection and cleavage were achieved for 2-3 hours using a TFA/TIPS/H2O (95:2.5:2.5) mixture. Peptidomimetics compounds were finally purified by preparative HPLC-MS, where the major diastereoisomer was isolated (S) when possible and a purity of >95% was obtained. Complete list of compounds used in this study is shown in Table S1. Detailed procedures for the synthesis and characterization of compounds 1 to 53 and 67 were described in our previous studies: synthesis of compounds 1 to 3 was described in Colombo et al. 2012 ^23^, compounds 4 to 15 was described in Duchêne et al. 2014 ^41^, compounds 16 to 44 was described in St-Georges et al. 2017 ^26^. The synthesis of compound 45 and 67 was described in Shapira et al. 2022 ^28^. The synthesis of compounds 46 to 53 was described in Colombo et al. 2024^24^. UPLC chromatograms and MS characterizations are available in supplementary data for the compounds 54-68 (Supplementary figures from S5 to S40 and tables S5 to S9).

**Scheme 1.**
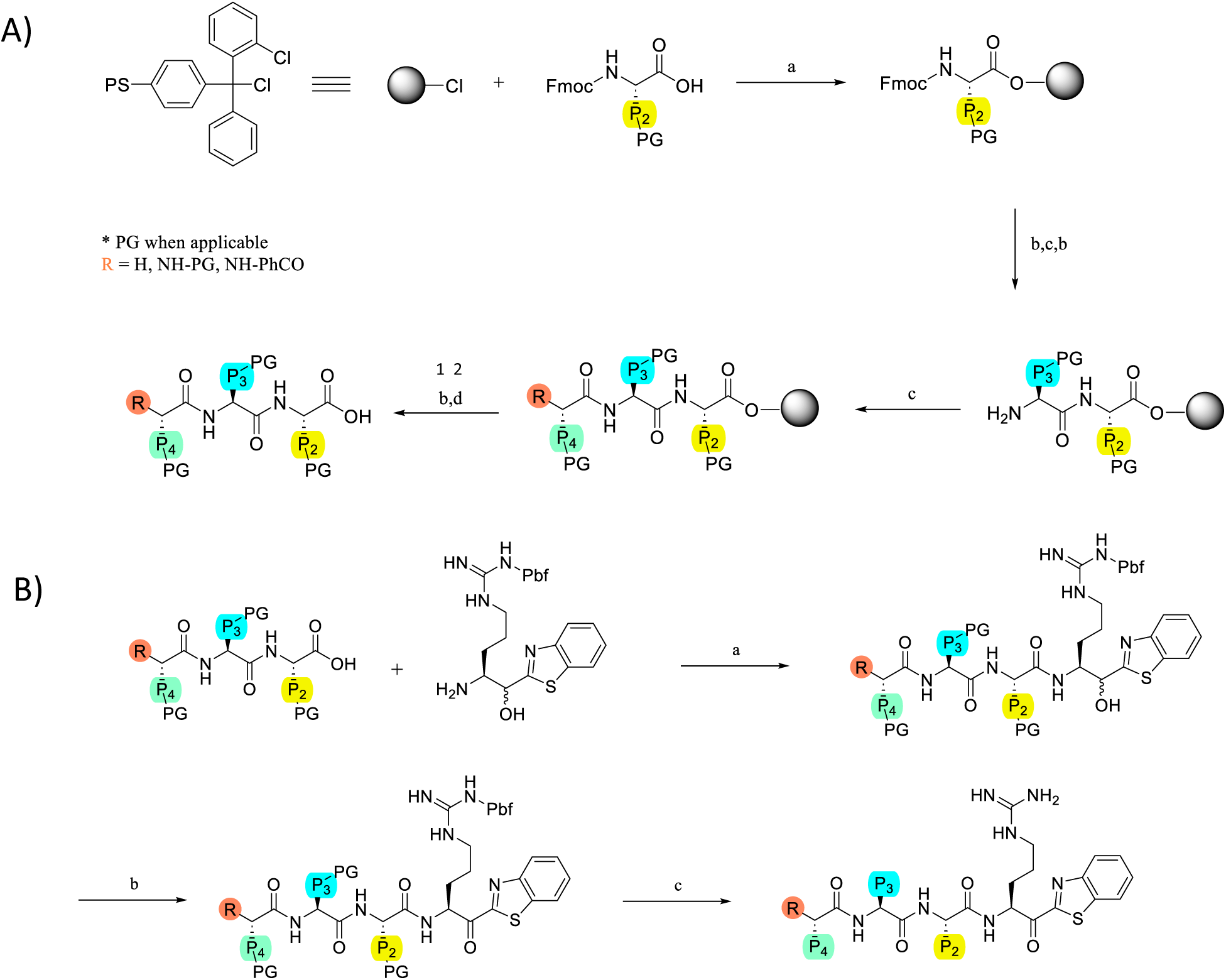
Synthesis. A) General synthesis of the peptidomimetic inhibitors: (a) Fmoc-AA, DIPEA, DCM, (b) Piperidine 20%/DMF, (c) Fmoc-AA, HATU, DIPEA, DMF, (d) 20% HFIP/DCM, r.t. (1: When applicable; 2: Capping group PhCO was added on the N-terminus part using standard peptide coupling conditions c). B) Warhead coupling to the protected peptides: (a) HATU, DIPEA, DMF, (b) DMP, DCM, (c) TFA/TIPS/H2O (95:2.5:2.5).

### *In vitro* inhibition assay

Recombinant TMPRSS13-AFAAS-6His (this study) and matriptase^42^ were expressed and purified as described. Recombinant human furin, Factor Xa (R&D Systems) and thrombin (MilliporeSigma) were from commercial sources. Assays were performed in a buffer composed of 50 mM HEPES pH 7.4, 1 mM β-mercaptoethanol, 1 mM CaCl2 and 500μg/ml BSA for furin and 50 mM HEPES pH 7.4, 150 mM NaCl and 0.1% BSA for other proteases. Assays were performed at room temperature using fluorogenic substrates (Boc-RVRR-AMC for furin and TMPRSS13, Boc-QAR-AMC for other proteases). Enzymatic activity was measured using an HTX Synergy microplate reader (Agilent) by fluorescence (excitation 360 nm and emission 460 nm) at room temperature. *K*_i_s for TMPRSS13 and Factor Xa (FXa) were derived from IC_50_ dose-response curves using the Cheng-Prusoff equation for reversible competitive inhibitors. For matripase, a plot of enzyme velocity as a function of inhibitor concentration were fitted with non-linear regression analysis to a Morrison *K*_i_ equation as previously described^27^.

### *In cellulo* inhibitory activity and IC50 determination

Vero C1008 were grown in 12-well plate (200K cells per well) and transfected for 24-hours using Lipofectamine 3000 (Life Technologies) with one of the following plasmids: pcDNA3.1 (Empty vector, EV), pcDNA3.1-TMPRSS13 WT or pcDNA3.1-TMPRSS13-AFAAS-6His. Cells were then washed 2 times with PBS before adding of HCell-100 medium (Wisent) containing indicated concentrations of each compound or control condition (DMSO 0.01%). After 24 h of incubation with the compounds, the medium was harvested and 90 μL were placed in 96-well plate wells with 10 µL of 200 μM Boc-RVRR-AMC. Fluorescence was acquired at room temperature in an FLx800 TBE microplate reader, with excitation at 360 nm and emission at 460 nm. The background was then subtracted by removing the signal recorded with the cells transfected with the empty vector (pcDNA3.1). Proteolytic activity results are represented as a percentage of the DMSO-treated condition. The IC_50_ was calculated by non-linear regression of compound concentration relative to velocity.

### Compound stability

For plasma stability, a 96-well plate with 2.5 μL of compound (1 mM aqueous solution) was incubated with 27 μL of plasma (from male Sprague−Dawley rat) at 37 °C for 0, 1, 2, 4, 7, and 24 h (or 0, 5, 15, 30, 60, 90, 120 min for less stable analogues). Degradation was quenched by adding 140 μL of acetonitrile-ethanol (1:1) solution containing N,N-dimethylbenzamide 0.25 mM (internal standard). This mixture was transferred to a filter plate Impact Protein Precipitation (Phenomenex, California), and another 96-well UPLC plate was placed at the bottom to collect filtrates. Both plates were centrifuged at 500 g for 10 min at 4 °C. The filtrates were diluted with 30 μL of distilled water and analyzed using an Acquity UPLC-MS system class H (column Acquity UPLC protein BEH C18 (2.1 mm × 50 mm), 1.7 μm particles with pore 130 Å). Peptide half-life was calculated from the degradation curve using the exponential one-phase decay function in GraphPad Prism 9. Mass spectra at the time point near half-life were compared with those at 0 min (plasma-inactivated before adding the compound) to identify cleavage fragments.

For proteolysis assay, human trypsin (#T6424, Sigma-Aldrich, Saint-Louis, Missouri, USA) was resuspended in 1mM HCl, 20 mM CaCl2, pH 3 and diluted to a final concentration of 0.1 mg/ml in PBS buffer (pH 7.4, 10 mM PO43-, 150 mM NaCl). The control peptide R8 and the compounds of interest were added to a 96 wells plate (37°C) to a final concentration of 10 µM. Trypsin (0.1 ug/ml) was added to the wells for 0, 0.5, 1, 2, 4 and 6h. The plate was quenched with AcN/EtOH(1:1) 1% formic acid, 0.25 mM N,N-dimethylbenzamide (IS) and centrifuged at 4°C. Samples were then analyzed by Acquity UPLC-MS system class H (column Acquity UPLC protein BEH C18 (2.1 mm × 50 mm), 1.7 μm particles with pore 130 Å). Sample half-life was calculated from the degradation curve using the exponential one-phase decay function in GraphPad Prism 9.

### Cytotoxicity

Vero C1008 cells (6 000 cells for sample and 5 000 to 90 000 cells for standard curve) were seeded in 96 wells plates. After 24h cells were washed with D-PBS and EMEM containing 10 % FBS and 1 % PSG were added. 10 µM of compounds were added for 24 h. Cell viability were determined with CellTiter-Glo 2.0 viability assay kit (Promega, G9242). Cellular viability is expressed relative to DMSO treated cells (vehicle). The luminescence readout was measured using the TriStar LB 942 multimode reader (Berthold Technologies). A standard curve of luminescence readout as a function of the number of cells was obtained by seeding known amounts of cells in the drug-exposed cell plate immediately before adding the reagent. Assays were performed independently at least three times in triplicate. The results were background corrected using negative controls and presented as the mean cell viability (%) compared to vehicle-treated cells ± standard deviation (SD).

### Molecular modelling

Molecular modeling studies were carried out using the Molecular Operating Environment (MOE) software, version MOE2022.02. The 3D coordinates of TMPRSS13 and matriptase were obtained from the Protein Data Bank, PDB ID: 6KD5 and 6N4T respectively. The models were loaded into MOE, and polar hydrogens and partial charges were added using the ‘Protonate 3D’ function of MOE. During the protonation process, a temperature of 300 K, a concentration of 0.1 mol/L salt in the solvent, and a pH of seven were used. Any missing atoms, alternate geometries, or other crystallographic artifacts were fixed using the ‘QuickPrep’ function. The structures were then energy-minimized in the Amber99 force field to achieve a root mean square (RMS) gradient of 0.1 kcal/mol. 3D models were created for all the compounds studied and their energies were minimized to an RMS gradient of 0.1 kcal/mol using the Amber99 force field. The structures were then protonated, and partial charges were calculated to assign ionization states and position hydrogens in the macromolecular structure based on its 3D coordinates. Before carrying out the conformational search, a covalent bond was formed between the serine of the catalytic triad and the ketone of the ketobenzothiazole. Compounds N-0388, N-0130, and N-0430 were chosen for molecular modeling studies based on the experimental assay results and were built using the ‘Builder’ tool.

A short molecular dynamics (MD) simulation was conducted on the inhibitor-enzyme complex using MOE and Berendsen equations (BER). The Berendsen velocity/position scaling methodology was used for system equilibration, as it is insensitive to configuration conditions and suitable for peptide conformations. The guanidine group of the arginine in P1 was constrained by key interactions with Asp at the bottom of the S1 pocket, and a covalent bond was created between the serine of the catalytic triad and the ketone of the warhead. The MD simulations were performed using the AMBER99 force field. The system was neutralized with Na^+^ and Cl^-^ ions at a concentration of 0.1 mol/l, and a water model was used for the solvation box. The enzyme-inhibitor solvated complex was energetically minimized for 10 ps. The system was then equilibrated to constant temperature (NVT) and pressure (NPT) conditions at 310 K and 1 atm for 100 ps each. An equilibrated system was used for the production run, which lasted for 1 ns. The trajectory with the lowest potential energy values was chosen for further analysis. The interactions, geometry, and orientation of all molecules in the binding cavity were analyzed and visualized using MOE. The ‘Align/Superpose’ function in MOE was used to align and superpose the complexes and sequences.

### Virus-Like Particles (VLP) production

VLP have been done as previously reported^43^. Briefly, HEK293T/17 were seeded at 6.10^6^ cells per T75 flask during 24 h and were transfected using Lipofectamine 3000 following manufacturer’s instructions. For transfection, 8 μg of reporter gene (Luciferase-ZsGreen, NR-52516), 6 μg helper plasmid (psPAX2, Addgene #12260), and 6 μg of SARS-CoV-2 Spike protein of the delta variant. VLP deprived of surface glycoprotein were produced by transfecting 6 μg of empty vector (pcDNA3.1) and were used as negative control. VLP quantification and normalization was as previously described^43^.

### VLP infection and inhibition assay

2 x 10^4^ Vero E6 cells were seeded in 96-well white culture plates and incubated 24h. The cells were transfected, as mentioned above, with 40 ng of TMPRSS13 plasmid or an empty vector (pcDNA3.1) and were incubated for 24h. A 3h pre-treatment was performed with 10 µM of the compounds. A normalized preparation of VLPs (δ variant or mock) containing 10 µM of the compounds was added to the wells and the cells were incubated 72h. The luminescence was detected with the ONE-Glo-EX luciferase assay (Promega, E8130) following the manufacturer’s instructions. Total luminescence was measured with a Berthold TriStar 2 LB 942 microplate reader.

### Statistical analyses

Data was obtained from at least three different experiments (mean +/- SD). Curves, diagrams, and statistical analysis was done using GraphPad Prism version 10.0.0 for Windows, GraphPad Software (Boston, Massachusetts USA ; www.graphpad.com). The size of the error bars indicates the S.D. within the data set. All our results are the average of at least 3 independents experiments.

## Results

### Production of a recombinant proteolytically active TMPRSS13 form

To identify TMPRSS13 inhibitors, we first expressed and purified a soluble, active recombinant form of this protein.. An initial TMPRSS13 construct containing a fused C-terminal His-Tag (schematic representation depicted in Fig.1A) was designed for expression and purification in mammalian Expi293 cells, a commonly used cell line for production of recombinant proteins^44^, with the intent to harvest and purify the spontaneously occurring shed form of TMPRSS13^8^. However, since TMPRSS13 is known to cleave at arginine or lysine^5,45,46^, C-terminal cleavages and potential loss of the His-tag were expected to arise during overexpression due to the presence of an Arg-Phe-Arg-Lys-Ser motif located at amino acid 558-562 (Fig.1A). To monitor recombinant protease integrity following overexpression in mammalian cells, a mass spectrometry analysis after benzamidine purification was performed on the media of Expi293 cells transfected with TMPRSS13-WT construct. This led to the identification of two spontaneous cleavage sites (R558 and R560) in the last five amino acids of the C-terminal region (Fig.1A, residues in red, mass spectra available in Fig.S1). Near these residues, another basic amino acid (K561) could also potentially be susceptible to cleavage by TMPRSS13. Thus, to prevent the loss of His-tag, we designed a construct containing protease resistant residues at these positions (R_558_-F-R-K-S_562_ → A_558_-F-A-A-S_562_).

**Figure 1.**
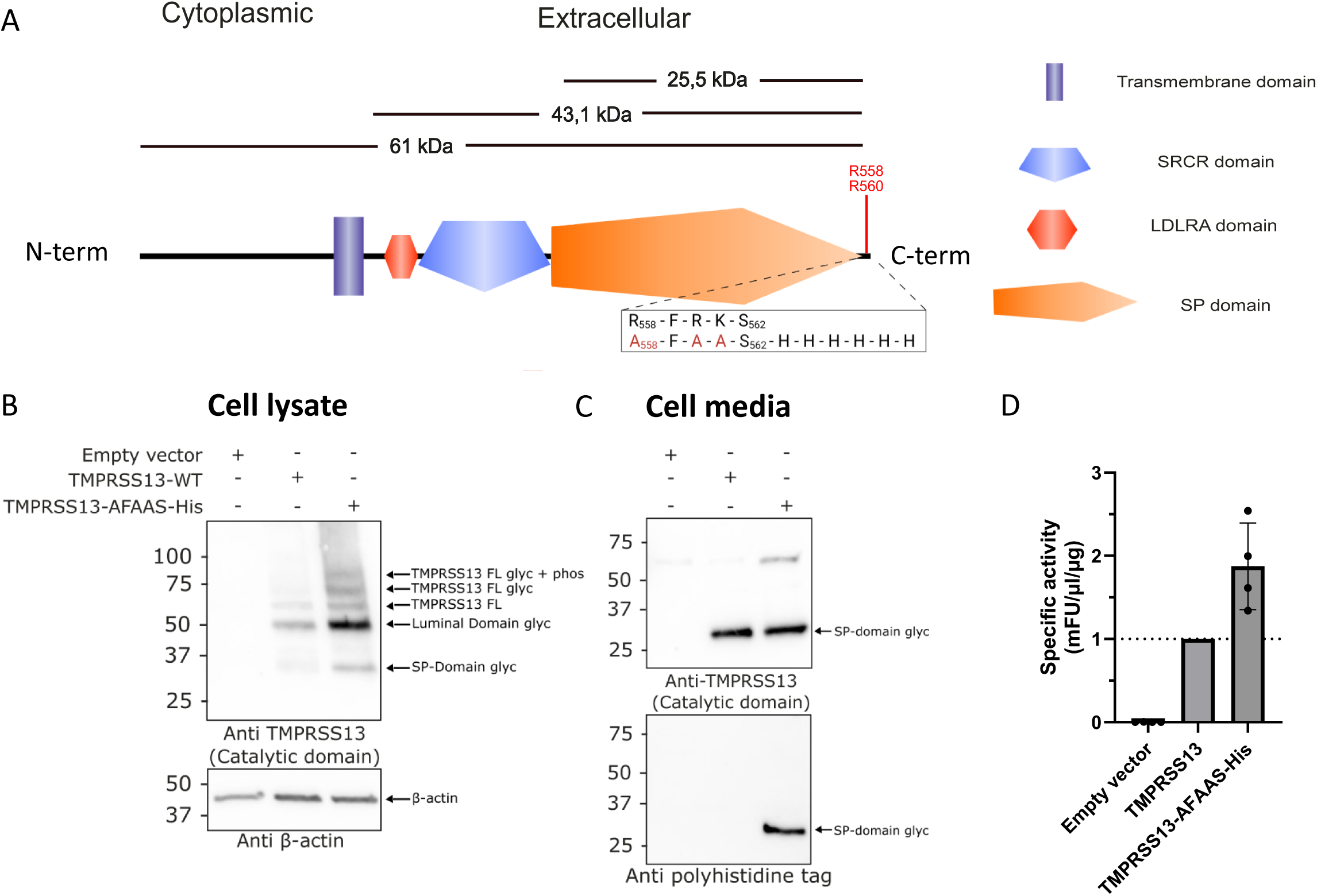
Schematic representation of TMPRSS13 and expression of active recombinant TMPRSS13 forms in HEK293SL cells. **A)** Representation of TMPRSS13. Depicted are the different domains of TMPRSS13, the theoretical molecular weight of specific segments, and a close-up view of the his-tagged mutated sequence. In red are noted the amino acids which are subject to spontaneous cleavage following expression and enrichment in the extracellular media (spectra in supplementary data 1). The theoretical weights of 25.5 kDa for the catalytic domain, 43.1 kDa for the extracellular part and 61 kDa for the total protein are given as an indication of what the weights would be without glycosylation. **B)** HEK293SL cells were transfected with pcDNA3.1 (empty vector), the full length TMPRSS13 (TMPRSS13-WT) and the his-tagged TMPRSS13 (TMPRSS13 -AFAAS-His). Proteins extracted from cell lysates were separated using 15% SDS-PAGE gels under reducing conditions. The separated proteins were then analyzed by Western blotting. Proteins were detected with the rabbit C-terminal TMPRSS13 antibody (PA5-30935), targeting the catalytic region of TMPRSS13, and the mouse anti-β-actin antibody (12262S). **C)** HEK293SL cells were transfected with an empty vector, the full length TMPRSS13 (TMPRSS13 WT) and the his-tagged TMPRSS13 (TMPRSS13 -AFAAS-His). Media concentrates were separated by SDS-PAGE under reducing conditions using 15% gels and then analyzed through Western blotting. Proteins were detected with the rabbit C-terminal TMPRSS13 antibody (PA5-30935), targeting the catalytic region of TMPRSS13, or the rabbit anti-His antibody (2365S). In these western blots, FL means Full length, Gly means Glycosylated, Phos means phosphorylated, and SP is the serine protease domain. **D)** Media samples from transfected HEK193SL cells were assessed for TMPRSS13 enzymatic activity using the Boc-RVRR-AMC substrate (100 µM). TMPRSS13 cleaves after an R-R site, releasing the AMC fluorophore. Fluorescence was measured every minute for 60 minutes. Results were normalized to WT TMPRSS13 specific activity.

To assess the expression pattern of TMPRSS13-AFAAS-6His constructs, HEK293SL cells were transfected, and both cell lysates and media were analysed by Western blot (Fig. 1B-C). Consistent with previous reports^39^, cell lysate analysis using an antibody targeting the protease catalytic domain revealed different TMPRSS13 entities corresponding to: a full-length glycosylated and phosphorylated form (∼80 kDa), a glycosylated form (∼70 kDa), a full-length unglycosylated form (∼61 kDa), a cleaved luminal form (∼50 kDa), and finally the active glycosylated protease domain (∼30 kDa)(Fig. 1B). Notably, although the expression pattern was identical for both constructs, the expression levels were higher with the AFAAS-6His construct. Shedding of TMPRSS13 catalytic domain was detected for both constructs in the cell media using an antibody directed against TMPRSS13 (Fig. 1C, upper panel), and was also observed for the AFAAS-6His construct using an anti-His-Tag antibody (Fig. 1C, lower panel).

Next, to determine whether the AFAAS-6His modifications had an impact on TMPRSS13 catalytic activity, cleavage of the Boc-RVRR-AMC fluorogenic substrate by WT or TMPRSS13-AFAAS-6HisTMPRSS13 was monitored in the cell culture media of transfected HEK293 cells (Fig. 1D). The media of cells transfected with AFAAS-6His construct did not show a loss of activity but exhibited an increase in specific activity when compared to the WT enzyme. These results confirm that the TMPRSS13-AFAAS-6His construct is suitable to use as a tool for recombinant protein production of soluble, active protease, with a similar expression pattern to the WT and robust catalytic activity associated with the shed form.

### Identification of potent TMPRSS13 inhibitors

To identify novel TMPRSS13 inhibitors and important molecular determinants for the inhibition of this protease, we screened a library of 65 peptidomimetic compounds containing various natural and non-natural amino acids as well as a ketobenzothiazole warhead (Fig. 2A for general scaffold and Table S1 for the detailed list of all compounds used in this study) using the TMPRSS13-AFAAS-6His recombinant protein purified from the media of Expi293-transfected cells. Of those 65 molecules, 54 reduced TMPRSS13 activity by more than 50 % at a concentration of 10 µM, 30 at 1 µM, and 11 at 100 nM (Fig. 2B). The ten peptidomimetic compounds that most significantly reduced the relative activity of TMPRSS13 were selected as hit molecules to be further characterized, and for each molecule, a new N-XXXX nomenclature was attributed (Table 1). Interestingly, all hits contain an arginine at the P4 position and nine of them a glutamine in P3 position, even though only 38% and 40 % of the library presented those residues at this position respectively. This suggests that a tetrapeptidic scaffold constituted of a P4 Arg and a P3 Gln is preferred to achieve potent inhibition of TMPRSS13.

**Figure 2.**
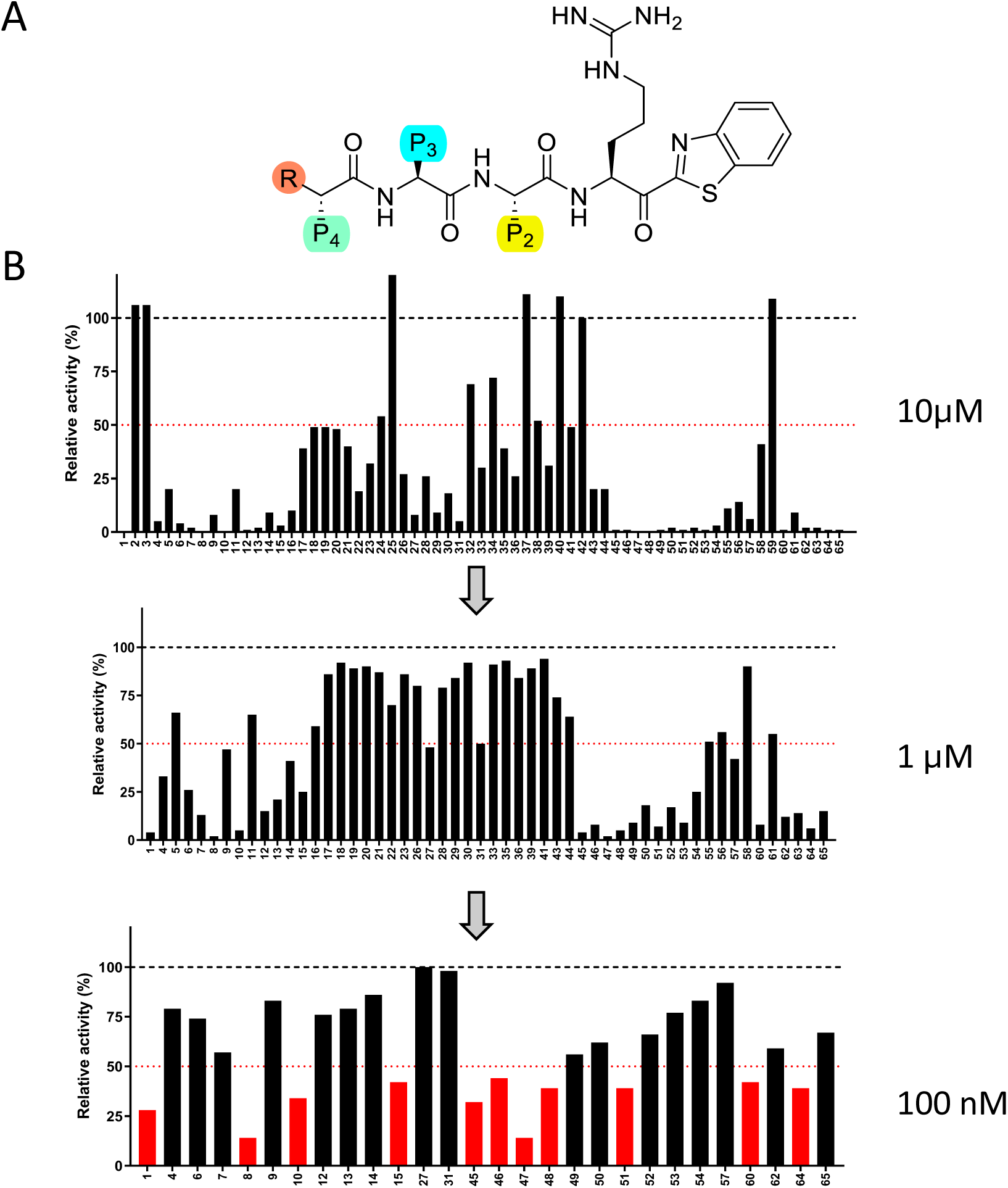
Screening of a library of 65-compound ketobenzothiazole-based against TMPRSS13. A) Scaffold of ketobenzothiazole-based inhibitors. P1 is an Arginine, P2, P3, and P4 in red were modified with different amino acids. In C-terminal position, a kentobenzothiazole, in orange, permit to trap serine in catalytic pocket. B) Compounds that were active at 10 µM were screened at 1µM then at 100 nM against recombinant TMPRSS13. Compounds affecting > 50% activity were selected for further characterization.

**Table 1.** Ki parameters for hits compounds of the original screen.

| Screening number | Compounds | Sequence |  |  |  |  |  | K<sub>i</sub> TMRSS13 (nM) |  |
| --- | --- | --- | --- | --- | --- | --- | --- | --- | --- |
|  |  | R | P4 | P3 | P2 | P1 | Warhead |  |  |
| 1 | N-0100 | - | R | Q | A | R | Kbt | 31.4 | ± 9.1 |
| 8 | N-0107 | - | R | Q | P | R | Kbt | 14.7 | ± 2.6 |
| 10 | N-0108 | - | R | Q | Y | R | Kbt | 22.3 | ± 6.9 |
| 15 | N-0117 | - | R | Q | L | R | Kbt | 119.9 | ± 41.9 |
| 45 | N-0678 | H | R | Q | Cha | R | Kbt | 30,0 | ± 10.4 |
| 47 | N-0159 | - | R | Q | hF | R | Kbt | 10.3 | ± 1.7 |
| 48 | N-0136 | - | R | Q | F(4F) | R | Kbt | 17.7 | ± 8.3 |
| 51 | N-0179 | - | hR | Q | A | R | Kbt | 24.8 | ± 5.1 |
| 60 | N-0439 | H | R | A | F | R | Kbt | 35.9 | ± 4.5 |
| 64 | N-0182 | PhCO | R | Q | A | R | Kbt | 28.5 | ± 4.3 |

K<sub>i</sub> were determined using dose-response curve and the Cheng-Prusoff equation for competitive inhibitors.

Next, the *in vitro* inhibition constant (*K*_i_) of these compounds towards TMPRSS13 was determined (Table 1). All selected compounds were confirmed as potent TMPRSS13 inhibitors in the nanomolar range and notably, the best compound from this selection, N-0159 (RQhFR-kbt), exhibited a *K*_i_ of 10.3 nM. Derivatives of N-0159 were next synthesized to investigate the contribution of each pharmacophore on TMPRSS13 inhibition (Table 2). We previously demonstrated that modification of the N-terminal amine, such as its deamination, was essential to achieve compound stability in biological contexts^24^. Thus, the N-0159 secondary N-terminal amine was removed yielding compound N-0430 ((H)RQhFR-kbt). This led to an inhibitor with a two-fold increase in potency at 5.3 nM. At the P2 position, the replacement of the unnatural homo-Phe (N-0430) with the proteogenic amino acid Phe (N-0130) led to a slight decrease in activity (*K*_i_ = 25 nM). Finally, the removal of Arg at P4 position in N-0388 resulted in a drastic reduction in TMPRSS13 inhibition yielding a compound with a Ki in the µM range (1.4 µM). Having identified N-0430 as our lead compound, we synthesized N-0430(OH), harboring a hydroxyl instead of the reactive ketone, which prevents the serine trap from being active. This compound was used as a negative control and, as expected, did not inhibit TMPRSS13 at 10 µM^43^. Hence, characterization of N-0159 derivatives led to the identification of a potent, low-nanomolar TMPRSS13 inhibitor and confirms that the presence of an Arg at P4 position in the peptidomimetic is a crucial molecular determinant for its inhibition.

**Table 2.** Ki parameters for N-0159 derivatives.

| Compounds | Sequence |  |  |  |  |  | K<sub>i</sub> TMPRSS13 (nM) |  |
| --- | --- | --- | --- | --- | --- | --- | --- | --- |
|  | R | P4 | P3 | P2 | P1 | Warhead |  |  |
| N-0430 | H | R | Q | hF | R | Kbt | 5.3 | ± 1.4 |
| N-0130 | H | R | Q | F | R | Kbt | 24.6 | ± 7,3 |
| N-0388 | H | - | Q | F | R | Kbt | 1443 | ± 1071 |
| N-0430(OH) | H | - | Q | hF | R | (OH)bt | > 10 000 |  |

K<sub>i</sub> were determined using dose-response curve and the Cheng-Prusoff equation for competitive inhibitors.

### Insights on pharmacodynamic proprieties of TMPRSS13 inhibitors

To investigate inhibitor selectivity, compounds N-0130, N-0430 and N-0388 were tested against a small panel of proteases including Matriptase, another TTSP implicated in cancer and viral entry^36,47^, as well as potential off-target proteases (FXa, thrombin and furin) (Fig. 3, Table S2). N-0430 and N-0130 were found to be sub-nanomolar inhibitors of matriptase, thus demonstrating better affinity for this protease than TMPRSS13. Removal of the P4 Arg in N-0388 also led to a decrease of affinity as in TMPRSS13, but the compound remained a potent matriptase inhibitor with a Ki of 24.2 nM. Compounds N-0430 and N-0130 also achieved nanomolar inhibition of FXa, although to a lesser degree than against TMPRSS13 with respective *K*_i’_s of 57 and 48 nM. Similarly to TMPRSS13 and matriptase, a loss of potency was detected after removing the P4 Arg in N-0388, with a *K*i of 1.2 µM against FXa. Finally, the three compounds do not inhibit or are weak inhibitors of both thrombin and furin. These results suggests that this first generation of TMPRSS13 inhibitors exhibits partial selectivity, with reduced potency against other families of serine proteases but high affinities against a member of the same family, matriptase. Interestingly, the introduction of a P4 Arg was essential to achieve nanomolar inhibition of TMPRSS13. Similarly, the P4 Arg positively impacted matriptase inhibition, although it was not necessary to achieve nanomolar inhibition of matriptase.

**Figure 3.**
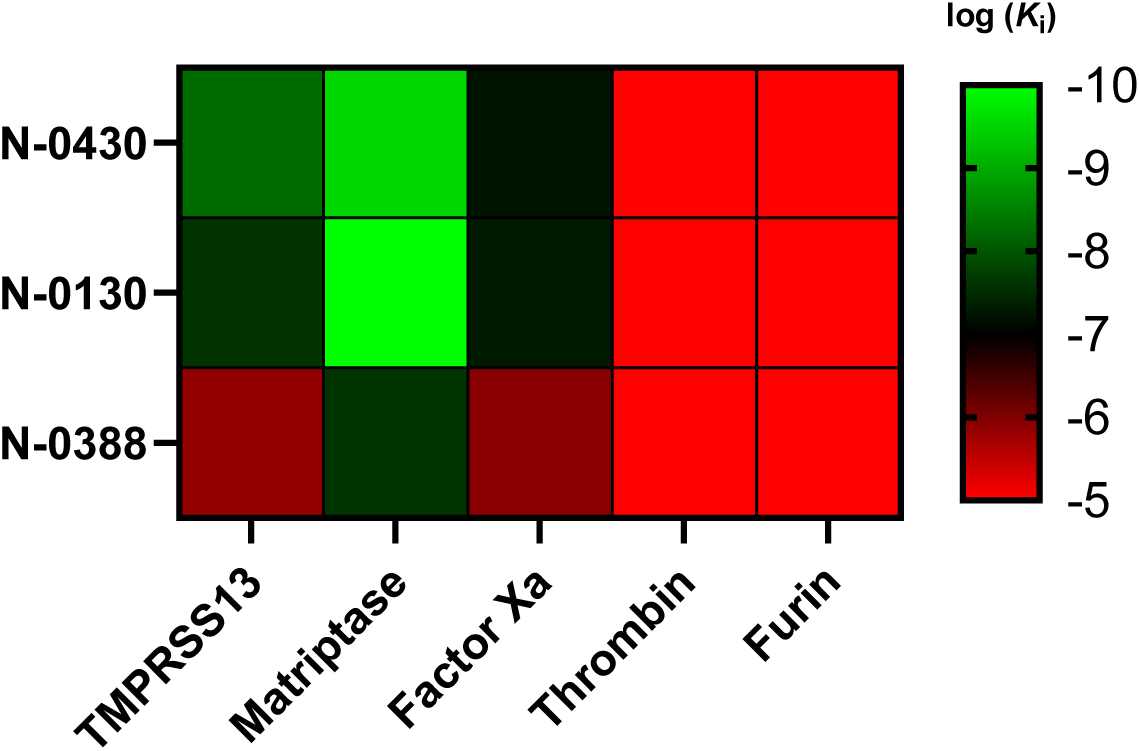
Selectivity heatmap of peptide inhibitors against fives targets. The heatmap displays the selectivity profiles of three compounds (N-0430, N-0130, and N-0388) tested against TMPRSS13, matriptase, Factor Xa, thrombin, and furin. The color scale on the right ranges from red (log(ki) = 5) to green (log(ki) = −10), indicating the logarithmic inhibitory constant (ki) values for each compound-enzyme pair. Higher log(ki) values (red) represent weaker inhibition, while lower log(ki) values (green) indicate stronger inhibition. Data are available in Table S2.

To better understand the impact of the addition of an Arg at the P4 position for TMPRSS13 inhibition, we performed molecular modeling. Compounds N-0130 ((H)RQFR-Kbt) and N-0388 ((H)QFR-Kbt), were first superimposed inside of TMPRSS13’s catalytic pocket (Fig. 4A, 2D interaction diagrams available for both compounds in Fig. S2). Constant interactions across compounds were observed through the formation of a similar hydrogen bond network between P1 Arg and residues Asp505, Ser506 and Gly534 of the S1 pocket. However, the inclusion of a supplementary Arg in P4 of N-0130 led to several important interactions, each around −9 to −8 kcal/mol (Table S3) with residues Tyr486, Asp487 and Ser488 present in TMPRSS13’s S4 pocket. Optimal binding at the P4 position required a noticeable conformational shift in the N-0130 peptide backbone, which led to a weakened interaction between its P3 Gln from Gln508 when comparing with N-0388 (−2.4 vs −4.5 kcal/mol respectively). A similar, but more subtle phenomenon was observed in matriptase (Fig. 4B) where the addition of a P4 Arg in N-0130 led to a gain of interactions with Gln783. However, the P3 Gln in N-0388 can interact with both residues Asp828 and Tyr755 of matriptase, compensating slightly for the lack of P4, which is consistent with the *in vitro* results. The preference of TMPRSS13 for tetrapeptides is also reflected in the overall shape of its S4 substrate pocket (Fig. 4C) compared to matriptase (Fig. 4D, detailed interaction properties available in Table S4). Both pockets are similar in length but differ strikingly in their layout. TMPRSS13 demonstrates a narrower hydrophobic pocket, resulting from Tyr489 and Asp411 hydrogen bond formation, while the corresponding amino acids in matriptase (Phe708 and Gln783) do not interact, forming an open groove. Thus, the TMPRSS13 S4 pocket is more restrictive, limiting inhibitor size and flexibility, whereas matriptase is more accommodating.

**Figure 4.**
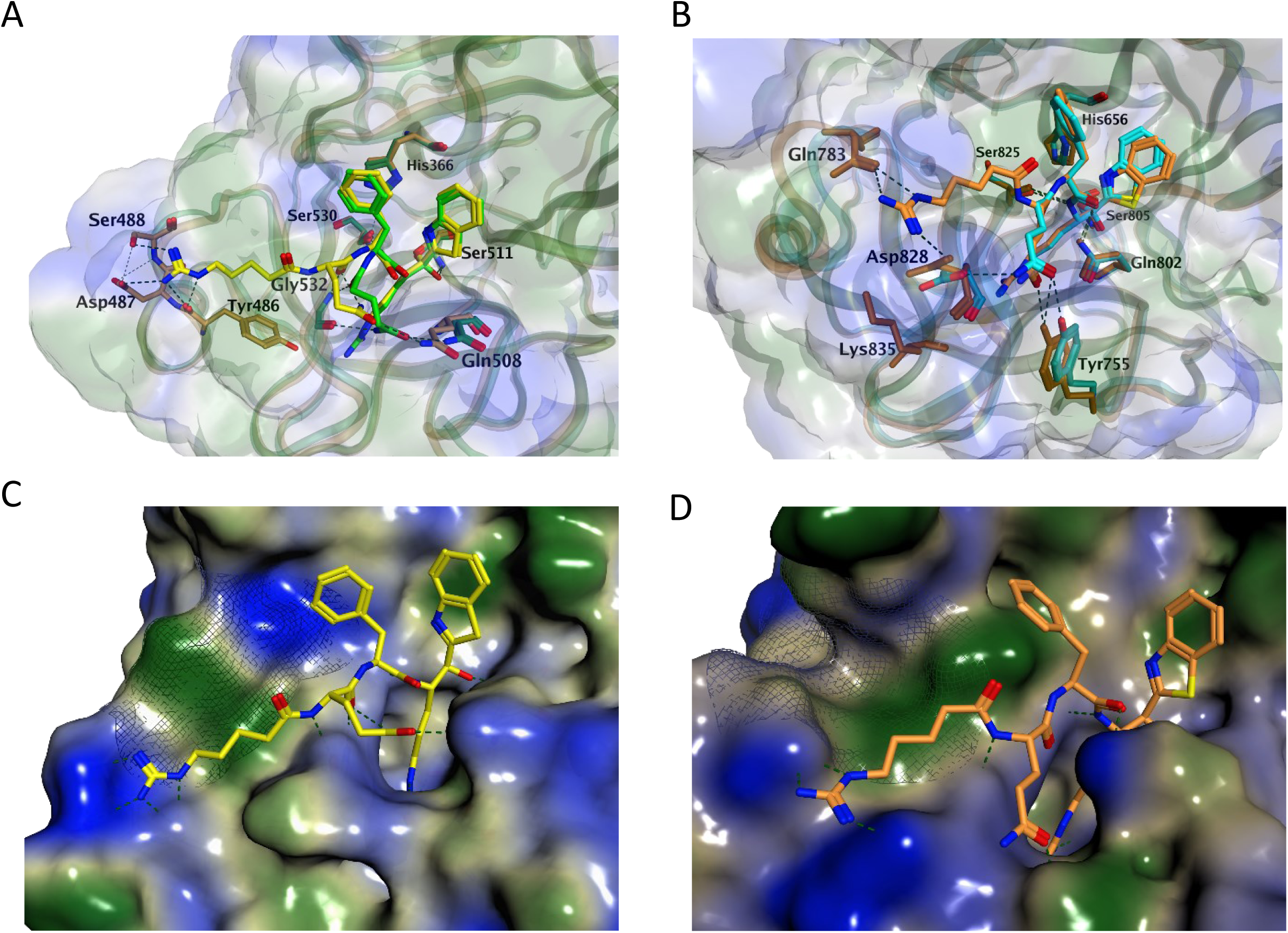
Molecular modeling. **A)** Docking of N-0130 ((H)RQFR-Kbt, yellow for compound and dark yellow for ligands) and N-0388 ((H)QFR-Kbt, green for compound and dark green for ligands) to TMPRSS13. **B)** Docking of N-0130 ((H)RQFR-Kbt, orange) and N-0388 ((H)QFR-Kbt, cyan) to matriptase. **C)** Docking of N-0130 ((H)RQFR-Kbt, yellow) in TMPRSS13. **D)** Docking of N-0130 ((H)RQFR-Kbt, orange) in Matriptase. For all panels, hydrophilic part of inhibitor is depicted in blue, hydrophobic in green, and residues involved in interactions are illustrated with dashed lines with each inhibitor are represented in stick form and labeled.

We observed that the presence of a P2 hPhe in N-0430 led to an increase in affinity of ̴5-fold when compared to N-0130 containing the natural Phe at this position. To explain this gain in potency, both compounds were docked inside TMPRSS13’s catalytic pocket (Fig. 5, Table S3). The side chain elongation in the hPhe not only allows better filling of the binding pocket but also enforces an intramolecular H-arene interaction between the aromatic cycle at P2 position and the α-carbon of the P3 position (position A), which stabilizes its conformation. This allows N-0430 to achieve an interaction between the carbonyl group in the backbone of its P2 position with Gln508, whereas N-0130 uses its P3 Gln to interact with this residue (position B). Although those two different interactions are of similar potencies in both compounds (−2.3 kcal/mol for N-0430 and −2.4 kcal/mol for N-0130), the P2 backbone interaction in N-0430 frees its P3 Gln to achieve an additional H-bond with Gln534 (position C), thus increasing the interaction with this residue (−8.1 kcal/mol for N-0430 and −4.6 kcal/mol for N-0130). Furthermore, N-0430 optimized conformation in TMPRSS13 also allows for an additional interaction with Asp487, lowering the interaction energy of −5.24 kcal/mol when compared to N-0130. These results support the detected increase in affinity of N-0430 towards TMPRSS13, when compared to N-0130. Overall, these results gave us insights on compound selectivity and enabled us to determine molecular interactions that favor TMPRSS13 inhibition.

**Figure 5.**
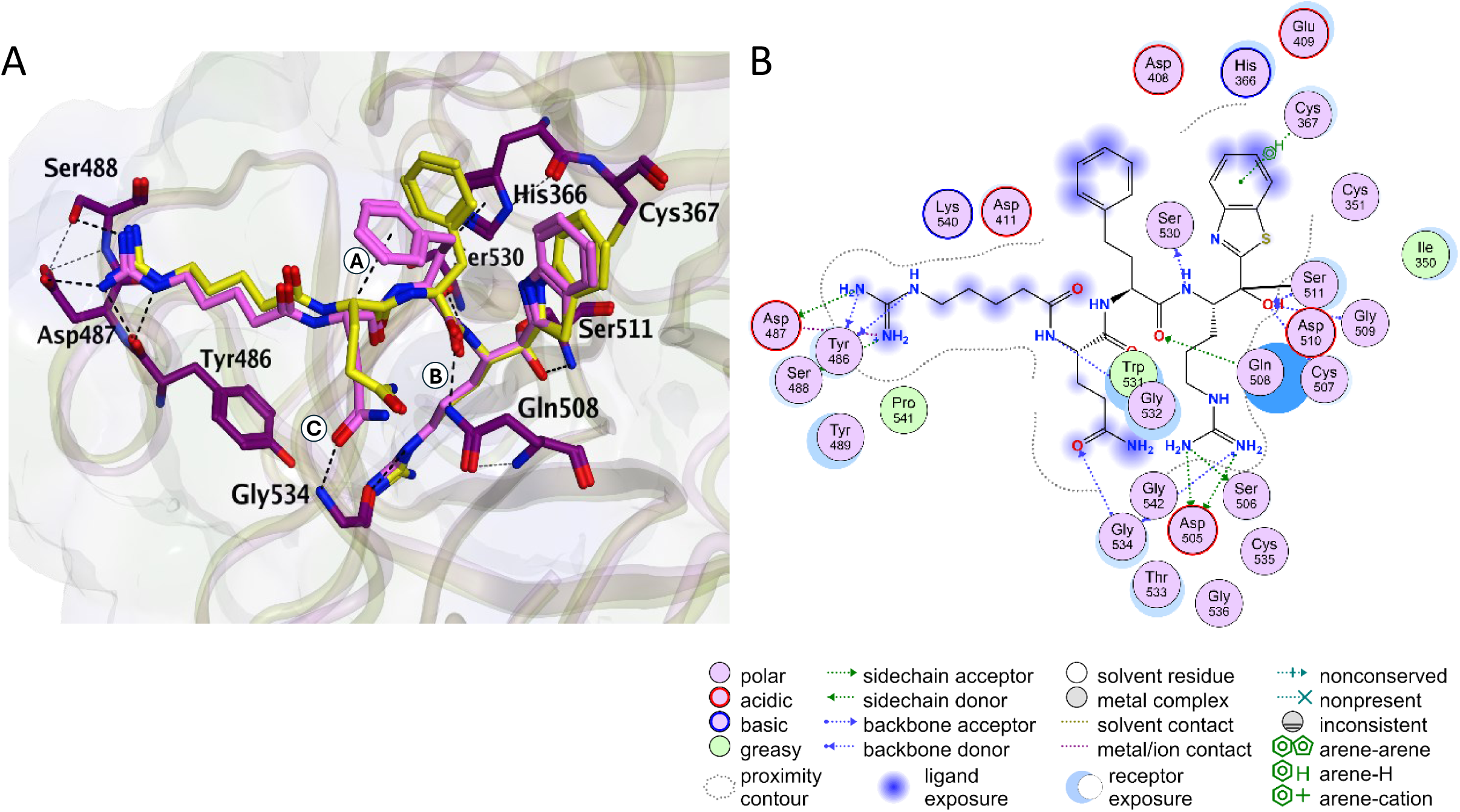
Representation of the interactions between N-0430 and TMPRSS13. Representations are in 3D (A) and 2D (B). N-0430 is shown in light pink stick mode and TMPRSS13 shown in purple stick mode. For comparison, N-0130 is superimposed and represented in yellow stick.

### *In cellulo* potency and stability of TMPRSS13 inhibitors

To assess whether inhibitors were potent in a cellular environment, compounds were next screened at 10 µM on WT TMPRSS13-tranfected Vero cells and media was harvested to measure activity (Fig. 6A). Potent inhibition of TMPRSS13 was observed for both deaminated tetrapeptides N-0130 (18.4 % residual activity) and N-0430 (9.0 % residual activity), which were more effective than N-0159 (37.7 % residual activity) containing the unmodified N-terminal amine. In line with the *in vitro* results, N-0388 presented limited inhibition of TMPRSS13 with residual activities of 72% and 92% respectively. Next, *in cellulo* TMPRSS13 inhibition by N-0130 and N-0430 was determined through the realization of dose-response curves (IC_50_) using the same assay (Fig. 6B-C). With an IC_50_ of 53 nM, compound N-0430 was found to be ̴4-fold more potent than N-0130. These results demonstrate that N-0430 is a potent *in cellulo* inhibitor of TMPRSS13, which was thus identified as the lead compound of this study.

**Figure 6.**
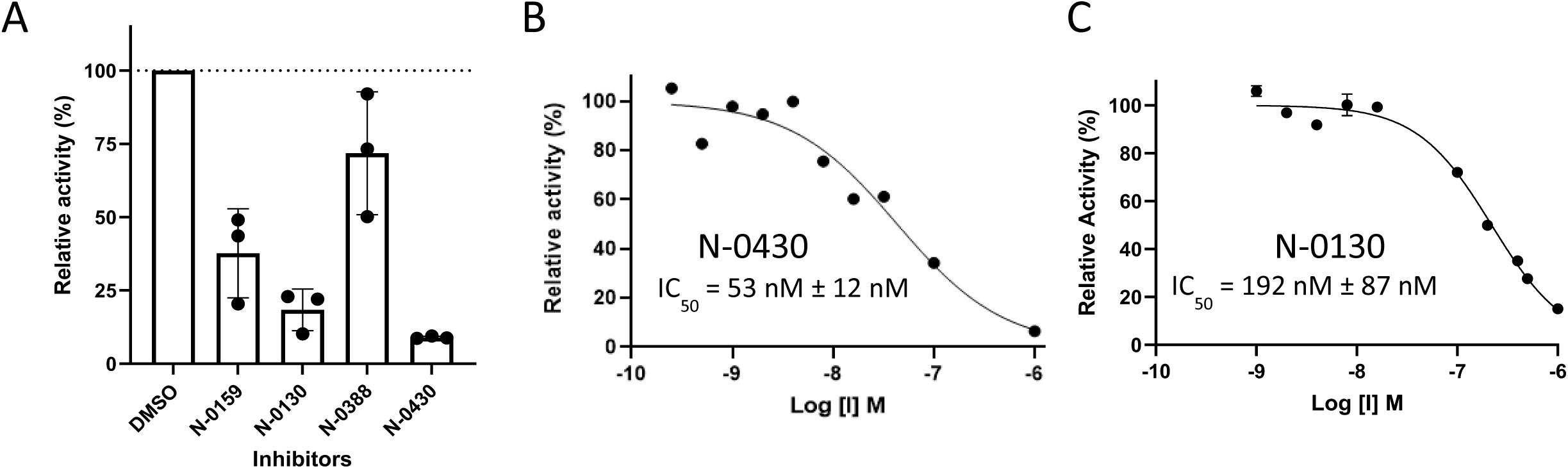
Evaluation of the potency in a cellular context. Vero cells were transfected with WT TMPRSS13, and activity was monitored using Boc-RVRR-AMC as a fluorogenic. (A) Relative activity of TMPRSS13 in the presence of 10 µM of compounds (N-0159, N-0430, N-0130, N-0388 and N-0430-OH) or vehicles (DMSO), displaying relative activity as a percentage of DMSO. (B) Relative activity of TMPRSS13 as a function of N-0430 concentration. (C) Relative activity of TMPRSS13 as a function of N-0130 concentration.

We next sought to investigate preliminary pharmacokinetic properties of selected compounds. To do so, the degradation rate of peptidomimetics N-0130, N-0159 and N-0430 was determined in presence of human trypsin and in rat plasma (Table 3). N-0130 presented a slightly better stability than the other compounds when incubated with trypsin, but all molecules achieved a half-life > 1 day. In rat plasma however, compound N-0159 presented a drastic decrease with a calculated half-life of 9.0 min, whereas both deaminated compounds were quite stable with measured half-life of 3.3 and 1.7 days for N-0130 and N-0430, respectively. As expected, these results demonstrate that the deamination of compounds N-0130 and N-0430 led to an improvement in their biological stability. To ensure that these results were not due to cytotoxicity, cellular toxicity tests were carried out with these different compounds, and none were identified as toxic at the highest concentration used (10 µM ; Fig. S3).

**Table 3.**
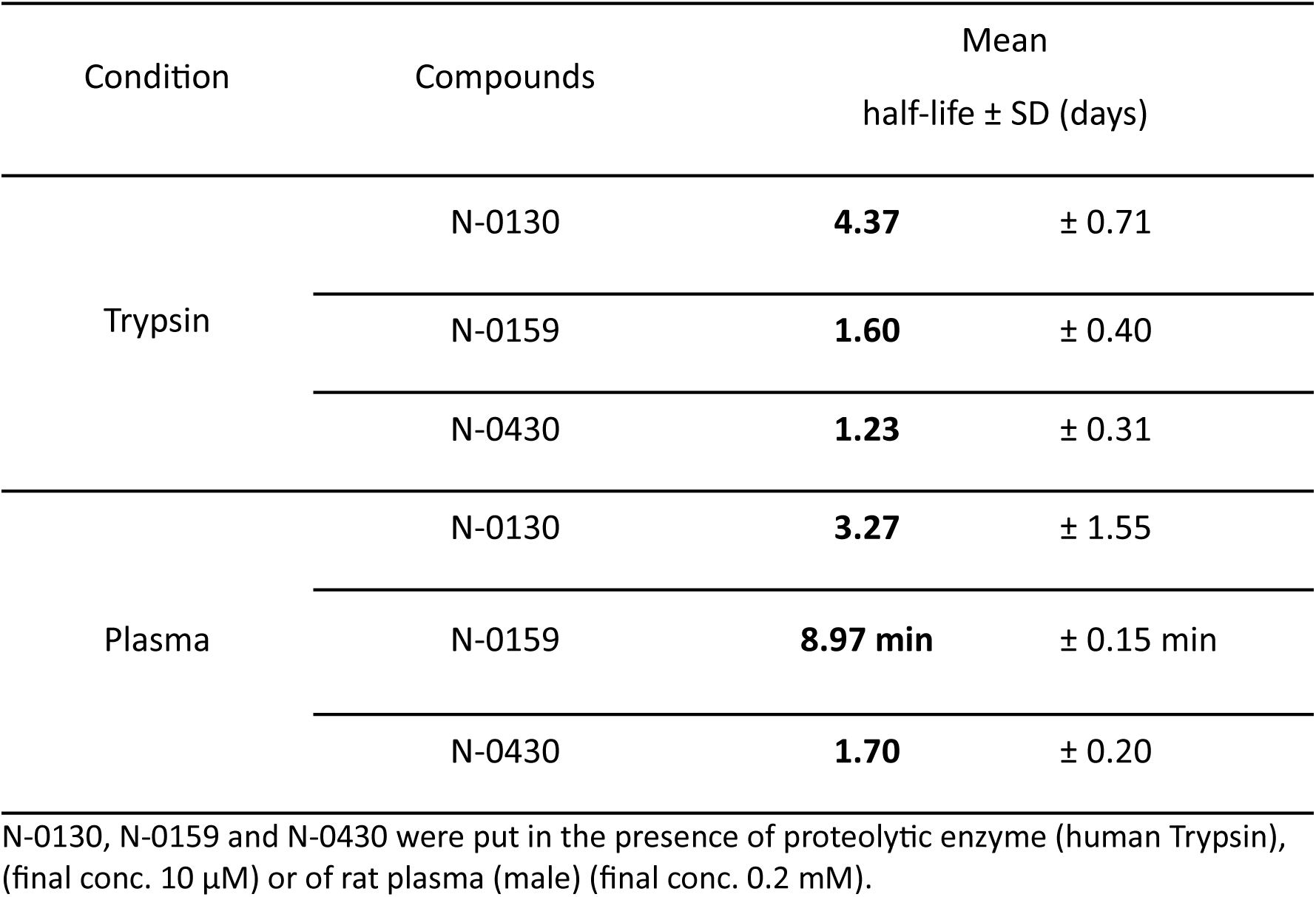
Stability of peptidomimetics N-0130, N-0159 and N-0430.

| Condition | Compounds | Mean<br>half-life ± SD (days) | |
| --- | --- | --- | --- |
| Trypsin | N-0130 | **4.37** | ± 0.71 |
| | N-0159 | **1.60** | ± 0.40 |
| | N-0430 | **1.23** | ± 0.31 |
| Plasma | N-0130 | **3.27** | ± 1.55 |
| | N-0159 | **8.97 min** | ± 0.15 min |
| | N-0430 | **1.70** | ± 0.20 |

N-0130, N-0159 and N-0430 were put in the presence of proteolytic enzyme (human Trypsin), (final conc. 10 μM) or of rat plasma (male) (final conc. 0.2 mM).

### Effect of N-0430 on the cellular entry of SARS-CoV-2 delta pseudovirus

TMPRSS13 has been associated with viral priming, including SARS-CoV-2^13,14,18,19^. To evaluate the potential use of these TMPRSS13 inhibitors, we developed a proof-of-concept assay using virus-like particles (VLPs) pseudotyped with SARS-CoV-2 delta variant. Vero E6 cells, transfected with either an empty plasmid (pcDNA3.1) or TMPRSS13-WT, were infected VLPs in presence of vehicle (DMSO), N-0430, or N-0430(OH) (Fig. 7). TMPRSS13 expression increased VLP entry in the vehicle condition while treatment with 10 µM N-0430 completely abolished the TMPRSS13-dependent VLP entry. Conversely, its inactive homolog, N-0430(OH) did not inhibit VLP entry. These results confirm that N-0430 achieved *in cellulo* inhibition of TMPRSS13 and demonstrated the potential use of this class of compounds for TMPRSS13-dependent pathological processes.

**Figure 7.**
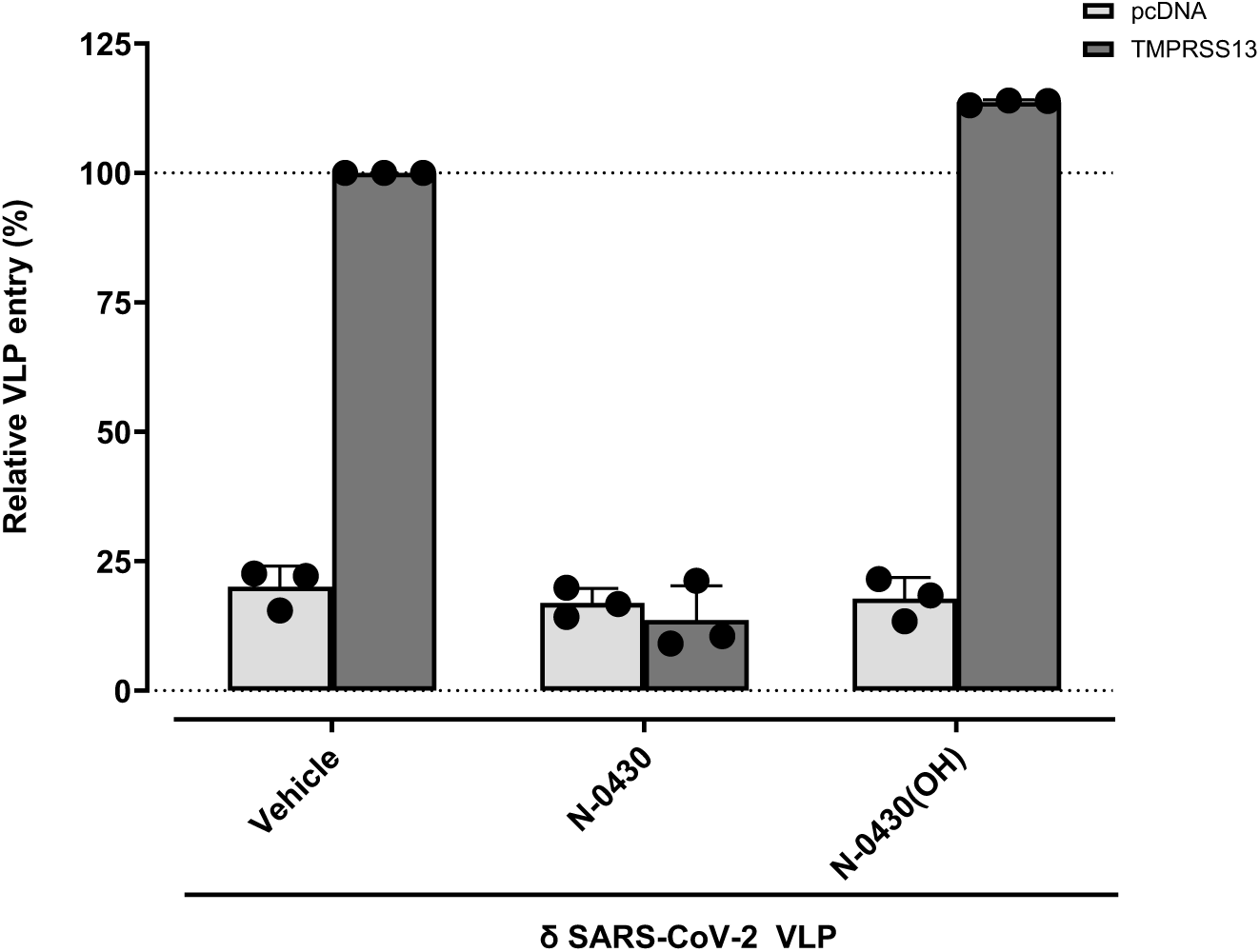
δ SARS-CoV-2 VLPs entry in human VERO E6 cells TMPRSS13 transfected in presence of N-0430, N-0430-OH and vehicles. VLPs infection in the presence of N-430 or N-430(OH) relative to the vehicle-treated condition (DMSO condition). Cells were transfected with TMPRSS13 WT 24 h before incubation with VLP. Compounds were added 3 h before and during the 72 h VLP incubation on cells. The background was subtracted from all particles using a VLP without surface glycoprotein for both experiments. Each experiment was repeated at least three times with vehicle condition, N-0430 and N-0430(OH), with five technical replicates per biological replicate.

## Discussion

Given the different roles of TMPRSS13 in pathologies such as cancer^11,12^ and viral entry^14,18^, selective enzymatic inhibitors could be valuable tools either for gaining new insights into the functions of TMPRSS13 or as a potential therapeutic agent. However, the availability of such inhibitors has been limited, with only broad-spectrum or irreversible inhibitors currently available.

In this study, a shed, soluble and active form of TMPRSS13 enabled the screening of a compound library against this protease. This led to the characterization of a first generation set of TMPRSS13 inhibitors and the identification of crucial molecular interactions contributing to their potency, thus representing a significant advancement in the development of selective inhibition of TMPRSS13. From an initial screening of 65 compounds, N-0159 was identified as the most potent. The synthesis of analogs of this compound led to the discovery of N-0430, which exhibited an *in vitro K*_i_ of 5.3 nM towards TMPRSS13. Compounds were selective for TMPRSS13 compared to thrombin and furin, for which they showed weak or no inhibition. However they were potent inhibitors of FXa, a serine protease implicated in the blood coagulation cascade, and matriptase, another TTSP of the same subfamily implicated in cancer as well as in viral entry^5,47^. This was expected since this class of compounds had previously achieved potent inhibition of these proteases^23,24,48^. Thus, depending on the mode of administration, the desired targets and pathological condition to treat, further optimization of these compounds will be necessary to achieve selectivity towards these enzymes.

Developed tetrapeptides N-0130 and N-0430 harboring a deaminated amino acid at P4 position demonstrated increased stability and were proven to be nanomolar *in cellulo* inhibitors of TMPRSS13. The inhibition of TMPRSS13 by N-0430 in a cellular context was further evidenced by its ability to block the entry of pseudotyped SARS-CoV-2 VLPs. These results are consistent with inhibition of TMPRSS13-dependant SARS-CoV-2 entry or cell fusion using camostat in other cell lines^16,49,50^. Beyond betacoronaviruses^14,51^, TMPRSS13 was also reported to activate monobasic cleavage sites of influenza A viruses^13,21^ and, more recently, SADS-CoV-2^19^. However, its role was found to be more pronounced in the activation of highly pathogenic influenza viruses (HPAIV) of the H5 subtype, characterized by a multibasic cleavage site containing a lysine at the P4 position (KKKR↓G), which are poorly activated by either furin or PC5/6^18,52^. Although cleavage sites of recent H5N1 HPAIV outbreaks of the clade 2.3.4.4b harbors an Arg at P4 position (RRKR↓G)^53^, suggesting cleavage by furin, sequences prone to TMPRSS13 proteolytis can naturally occur as they were previously detected in poultry^54^. Thus, selective TMPRSS13 inhibition or its inclusion in a pan-viral priming protease inhibitor could be of significant interest.

Inhibition of TMPRSS13 proteolytic activity could also be valuable for investigating its role in cancer and develop therapeutics. TMPRSS13 silencing is currently employed to decipher its implication in colorectal and breast cancer^11,12^. Although the use of siRNA effectively reduces expression, it does not discriminate between the loss of enzymatic activity and disruption of protein-protein interactions. Therefore, the development of TMPRSS13 inhibitors could provide a supplementary tool to distinguish these functions, thereby advancing our understanding of TMPRSS13 in oncogenesis^11^.

SAR analysis using the *in vitro* results and molecular modeling provided insights into key structural elements required to achieve nanomolar potency towards TMPRSS13. This analysis helps us better understand why our lead compound, N-0430, features a hPhe at the P2 position and an Arg at the P4 position. The gain in potency, through the elongation of the Phe residue, is proposed to occur via an intramolecular interaction that stabilizes the optimal compound conformation and improves pocket occupancy, leading to more potent interactions with TMPRSS13 residues. Furthermore, the presence of an Arg at P4 position was confirmed to be critical for TMPRSS13 inhibition, as its removal led to a substantial decrease in compound potency. At this position, the TMPRSS13 catalytic site presented a closed shape, and the presence of amino acids complementary to the positively charged guanidium group of Arg led to the formation of several crucial potent interactions. This correlates with the TMPRSS13 crystal structure, which showed that the TMPRSS13 substrate pocket recognizes both Arg and Lys residues at the P4 position^22^.

Docking poses of tri- or tetrapeptidomimetic compounds were also compared to matriptase binding sites. Similar to TMPRSS13, the removal of Arg at the P4 position led to a decrease in compound potency towards matriptase. This was expected, as this moiety was reported to benefit substrate cleavage or matriptase inhibition^41,55,56^. However, it was not required for achieving nanomolar inhibition of matriptase as it was the case for TMPRSS13. Modelling studies highlight a more open S4 pocket in matriptase, which could accommodate alternative inhibitor conformations. Identified differences in the overall shape of the S4 pocket between TMPRSS13 and matriptase could be exploited, through the use of arginine or lysine analogs and isosteres, to achieve selectivity. Recently, the use of a hybrid combinatorial substrate library (HyCoSul) led to the identification of several unnatural P4 residues that could modulate potency and specificity of matriptase, hepsin and hepatocyte growth factor activator (HGFA)^56,57^. This suggests an opportunity to further increase affinity of compounds for TMPRSS13 compared to other TTSPs such as matriptase.

Overall, this study presents the first generation of ketobenzothiazole-based inhibitors designed to target TMPRSS13. Among these, N-0430 was demonstrated to be a potent *in cellulo* inhibitor of this protease. Further optimization of these compounds will be necessary to achieve selectivity against closely related TTSP. Such compounds could serve as molecular tools to confirm the involvement of TMPRSS13 proteolytic activity in various pathologies, including colorectal and breast cancers. Moreover, TMPRSS13 inhibition could have a significant potential in antiviral preparedness, particularly in the development of broad-spectrum antiviral therapies.

## Acknowledgments

We thank Dominique Lévesque (Proteomics Platform by Mass Spectrometry at the Université de Sherbrooke) for his valuable assistance in mass spectrometry analysis.

## Author contributions

A.J., A.D., G.L., K.L., P.-L.B., R.L. participated in conception and design ; A.J., A.D., G.L., W.C., A.G-T, M.H., M.L., S.L., U.F. analysis and interpretation of the data ; A.J., A.D., G.L., W.C., A.G-T, M.H. drafting of the paper ; A.J., A.D., G.L., K.L., P.-L.B., R.L. revising it critically for intellectual content and the final approval of the version to be published. All authors have read and agreed to the published version of the manuscript.

