## Supplementary Data for "Development of ketobenzothiazole-based peptidomimetic TMPRSS13 inhibitors with low nanomolar potency"

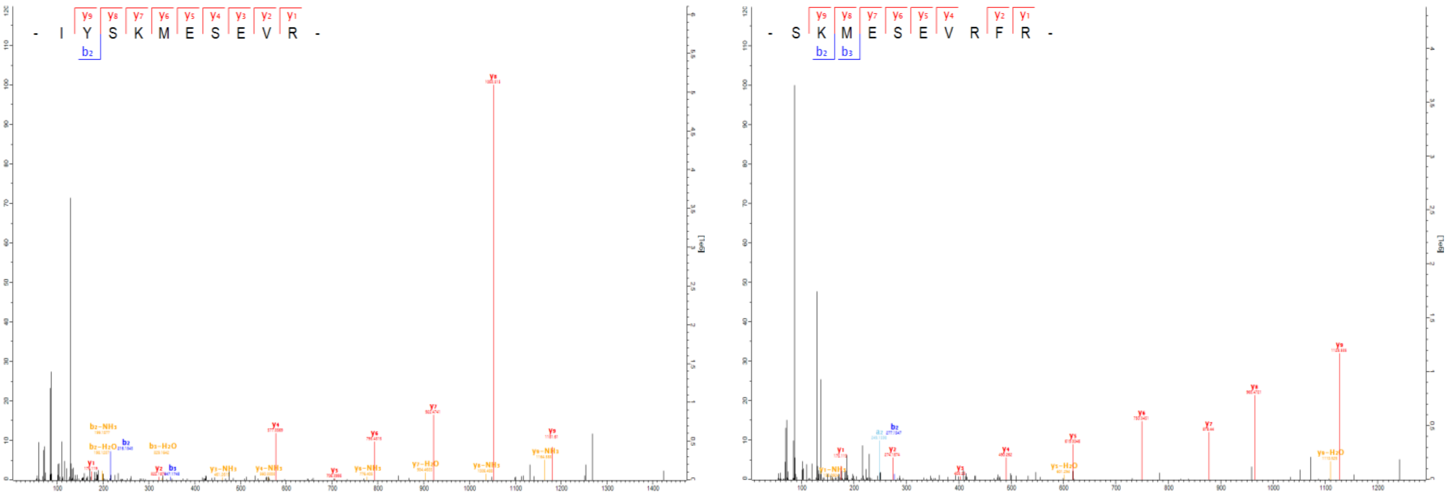

**Figure S1. Tandem mass spectrometry spectra showing cleavage of TMPRSS13 at R558 and R560.** Tandem mass spectrometry spectra from peptide ion trap collision-induced dissociation fragmentation for peptides IYSKMESEVR and SKMESEVRFR identified after in vitro digestion with chymotrypsin and LC-MS/MS analysis of fragments generated by TMPRSS13 cleavage. Here is a representative annotated MS/MS fragmentation spectrum with the identified matched N terminus-containing ions (b ions) in blue, peptides formed by internal cleavage within the protein sequence, often arising from nonspecific cleavages or internal fragmentation (a ion) in cyan, and the C terminus-containing ions (y ions) in red. Peptides that have undergone loss of ammonia (NH3) during fragmentation are identified as $Y\text{-}NH_3$ and Peptides that have undergone loss of water (H2O) during fragmentation are identified as $Y\text{-}H_2O$. Peptide intensities were summarized per amino acid residue and plotted in relation to each other. The detected peptide sequences indicate that R558 and R560 are mains cleavage sites.

**Table S1 - Overview of Peptide Inhibitors Screened Against TMPRSS13**

| # Compound | Position | | | | | Warhead | Molecular formula |
|---|---|---|---|---|---|---|---|
| | R | $P_4$ | $P_3$ | $P_2$ | $P_1$ | | |
| 1 | $NH_2$ | Arg | Gln | Ala | Arg | Kbt | $C_{27}H_{42}N_{12}O_5S$ |
| 2 | $NH_2$ | Arg | Gln | Ala | Arg | (OH)-Kbt | $C_{27}H_{44}N_{12}O_5S$ |
| 3 | $NH_2$ | - | - | - | Arg | Kbt | $C_{13}H_{17}N_5OS$ |
| 4 | $NH_2$ | Arg | Leu | Ser | Arg | Kbt | $C_{28}H_{45}N_{11}O_5S$ |
| 5 | $NH_2$ | Trp | Arg | Glu | Arg | Kbt | $C_{35}H_{46}N_{12}O_6S$ |
| 6 | $NH_2$ | Lys | Asn | Ala | Arg | Kbt | $C_{26}H_{40}N_{10}O_5S$ |
| 7 | $NH_2$ | Arg | Asn | Pro | Arg | Kbt | $C_{28}H_{42}N_{12}O_5S$ |
| 8 | $NH_2$ | Arg | Gln | Pro | Arg | Kbt | $C_{29}H_{44}N_{12}O_5S$ |
| 9 | $NH_2$ | Leu | Gln | Ala | Arg | Kbt | $C_{27}H_{41}N_9O_5S$ |
| 10 | $NH_2$ | Arg | Gln | Tyr | Arg | Kbt | $C_{33}H_{46}N_{12}O_6S$ |
| 11 | $NH_2$ | Ser | Gln | Ala | Arg | Kbt | $C_{24}H_{35}N_9O_6S$ |
| 12 | $NH_2$ | Trp | Cys | Tyr | Arg | Kbt | $C_{36}H_{41}N_9O_5S_2$ |
| 13 | $NH_2$ | Leu | Trp | Trp | Arg | Kbt | $C_{41}H_{48}N_{10}O_4S$ |
| 14 | $NH_2$ | Tyr | Lys | Ala | Arg | Kbt | $C_{31}H_{43}N_9O_5S$ |
| 15 | $NH_2$ | Arg | Leu | Gln | Arg | Kbt | $C_{30}H_{48}N_{12}O_5S$ |
| 16 | H | Tyr | Tyr | Tyr | Arg | Kbt | $C_{40}H_{43}N_7O_7S$ |
| 17 | H | Ala(Ada) | Tyr | Val | Arg | Kbt | $C_{40}H_{53}N_7O_5S$ |
| 18 | H | Leu | Tyr | Val | Arg | Kbt | $C_{33}H_{45}N_7O_5S$ |
| 19 | H | Phe | Tyr | Val | Arg | Kbt | $C_{36}H_{43}N_7O_5S$ |
| 20 | $NH_2$ | Tyr | Phe | Val | Arg | Kbt | $C_{36}H_{44}N_8O_5S$ |
| 21 | $NH_2$ | Tyr | (3-Cl)Phe | Val | Arg | Kbt | $C_{36}H_{43}ClN_8O_5S$ |
| 22 | H | Phe | Ala(Thiazol-4-yl) | Val | Arg | Kbt | $C_{33}H_{40}N_8O_4S_2$ |
| 23 | H | Phe | (Obn)Phe | Val | Arg | Kbt | $C_{43}H_{49}N_7O_5S$ |

| | | | | | | | |
|---|---|---|---|---|---|---|---|
| 24 | H | Phe | *h*Phe | Val | Arg | Kbt | $C_{37}H_{45}N_7O_4S$ |
| 25 | H | Phe | Tyr | Dip | Arg | Kbt | $C_{46}H_{47}N_7O_5S$ |
| 26 | $NH_2$ | Tyr | Tyr | His | Arg | Kbt | $C_{37}H_{42}N_{10}O_6S$ |
| 27 | $NH_2$ | Tyr | Tyr | (2, 3-OH)Phe | Arg | Kbt | $C_{40}H_{44}N_8O_8S$ |
| 28 | $NH_2$ | Tyr | Tyr | Ala(Thiazol-4-yl) | Arg | Kbt | $C_{37}H_{41}N_9O_6S_2$ |
| 29 | $NH_2$ | Tyr | Tyr | 2-Nal | Arg | Kbt | $C_{44}H_{46}N_8O_6S$ |
| 30 | $NH_2$ | Tyr | Tyr | Ala[3-(2-Thienyl)] | Arg | Kbt | $C_{37}H_{41}N_9O_6S_2$ |
| 31 | $NH_2$ | Tyr | Tyr | Trp | Arg | Kbt | $C_{42}H_{45}N_9O_6S$ |
| 32 | H | Phe | *h*Phe | (D)Val | Arg | Kbt | $C_{37}H_{45}N_7O_4S$ |
| 33 | H | Phe | *h*Phe | Leu | Arg | Kbt | $C_{38}H_{47}N_7O_4S$ |
| 34 | H | Phe | *h*Phe | Ile | Arg | Kbt | $C_{38}H_{47}N_7O_4S$ |
| 35 | H | Phe | *h*Phe | (D)Ile | Arg | Kbt | $C_{38}H_{47}N_7O_4S$ |
| 36 | H | Phe | *h*Phe | *Allo*Ile | Arg | Kbt | $C_{38}H_{47}N_7O_4S$ |
| 37 | H | Phe | *h*Phe | (D)*Allo*Ile | Arg | Kbt | $C_{38}H_{47}N_7O_4S$ |
| 38 | H | Phe | *h*Phe | Abu | Arg | Kbt | $C_{36}H_{43}N_7O_4S$ |
| 39 | H | Phe | *h*Phe | Nva | Arg | Kbt | $C_{37}H_{45}N_7O_4S$ |
| 40 | H | Phe | *h*Phe | (*t*-butyl)Gly | Arg | Kbt | $C_{38}H_{47}N_7O_4S$ |
| 41 | H | Phe | *h*Phe | Thr | Arg | Kbt | $C_{36}H_{43}N_7O_5S$ |
| 42 | H | Phe | *h*Phe | *AlloThr* | Arg | Kbt | $C_{36}H_{43}N_7O_5S$ |
| 43 | $NH_2$ | Tyr | Tyr | Val | Arg | Kbt | $C_{36}H_{44}N_8O_6S$ |
| 44 | H | Tyr | Tyr | Val | Arg | Kbt | $C_{36}H_{43}N_7O_6S$ |
| 45 | H | Arg | Gln | Cha | Arg | Kbt | $C_{33}H_{51}N_{11}O_5S$ |
| 46 | $NH_2$ | Arg | Gln | (4-$NO_2$)Phe | Arg | Kbt | $C_{33}H_{45}N_{13}O_7S$ |
| 47 | $NH_2$ | Arg | Gln | *h*Phe | Arg | Kbt | $C_{34}H_{48}N_{12}O_5S$ |
| 48 | $NH_2$ | Arg | Gln | (4-F)Phe | Arg | Kbt | $C_{33}H_{45}FN_{12}O_5S$ |
| 49 | $NH_2$ | Arg | Gln | (4-Cl)Phe | Arg | Kbt | $C_{33}H_{45}ClN_{12}O_5S$ |

| | | | | | | | |
|---|---|---|---|---|---|---|---|
| 50 | $NH_2$ | Arg | (NMe)Gln | Ala | Arg | Kbt | $C_{28}H_{44}N_{12}O_5S$ |
| 51 | $NH_2$ | *h*Arg | Gln | Ala | Arg | Kbt | $C_{28}H_{44}N_{12}O_5S$ |
| 52 | $NH_2$ | (D)Arg | Gln | Ala | Arg | Kbt | $C_{27}H_{42}N_{12}O_5S$ |
| 53 | $NH_2$ | (D)Arg | (D)Gln | Ala | Arg | Kbt | $C_{27}H_{42}N_{12}O_5S$ |
| 54 | $NH_2$ | Ile | Arg | Ala | Arg | Kbt | $C_{28}H_{45}N_{11}O_4S$ |
| 55 | $NH_2$ | Ile | Gln | Ala | Arg | Kbt | $C_{27}H_{41}N_9O_5S$ |
| 56 | $NH_2$ | Arg | Gln | Asp | Arg | Kbt | $C_{28}H_{42}N_{12}O_7S$ |
| 57 | $NH_2$ | Arg | Leu | Ala | Arg | Kbt | $C_{28}H_{45}N_{11}O_4S$ |
| 58 | $NH_2$ | Arg | Gln | (4-$CF_3$)Phe | Arg | Kbt | $C_{34}H_{45}N_{12}O_5S$ |
| 59 | $NH_2$ | Arg | Gln | Ala | Arg | Kbt | $C_{27}H_{42}N_{12}O_5S$ |
| 60 | H | Arg | Ala | Phe | Arg | Kbt | $C_{31}H_{42}N_{10}O_4S$ |
| 61 | H | Ser | Gln | Phe | Arg | Kbt | $C_{30}H_{38}N_8O_6S$ |
| 62 | H | Arg | Gln | Ser | Arg | Kbt | $C_{27}H_{41}N_{11}O_6S$ |
| 63 | H | Arg | Gln | Bpa | Arg | Kbt | $C_{40}H_{49}N_{11}O_6S$ |
| 64 | PhCO | Arg | Gln | Ala | Arg | Kbt | $C_{34}H_{46}N_{12}O_6S$ |
| 65 | - | - | [GABA] | Phe | Arg | Kbt | $C_{31}H_{38}N_8O_5S$ |
| 66 | H | Arg | Gln | *h*Phe | Arg | Kbt | $C_{34}H_{47}N_{11}O_5S$ |
| 67 | - | - | (H)Gln | Phe | Arg | Kbt | $C_{27}H_{33}N_7O_4S$ |
| 68 | H | Arg | Gln | *h*Phe | Arg | (OH)-bt | $C_{34}H_{49}N_{11}O_5S$ |

Table listing the peptide inhibitors. The first column indicates the compound number. The second column lists R, the third column lists *P4*, the fourth column lists *P3*, the fifth column lists *P2*, the sixth column lists *P1,* the seventh column *list* the warheads, and the eighth column provides the molecular formula of the peptide.

**Table S2. Inhibitory constants of peptide inhibitors against five targets.**

| | TMPRSS13 | Matriptase | Factor Xa | Thrombin | Furin |
|---|---|---|---|---|---|
| N-0430 | 5.3 ± 1.4 | 0.28 ± 0.05 | 56.9 ± 9.0 | >10000 | >10000 |
| N-0130 | 24.6 ± 7.3 | 0.13 ± 0.03[a] | 48.2 ± 3.7 | 8231 ± 1546[a] | >10000[a] |
| N-0388 | 1443 ± 1071 | 24.2 ± 4.5 | 1204.0 ± 166.6 | >10000 | >10000 |

This table presents the inhibitory constants ki (nM) for three compounds (N-0430, N-0130, and N-0388) tested against TMPRSS13, matriptase, Factor Xa, thrombin, and furin. The Values provided were used to generate the selectivity heatmap in figure 3. [a] As published in Shapira et al. 2022 Nature[1].

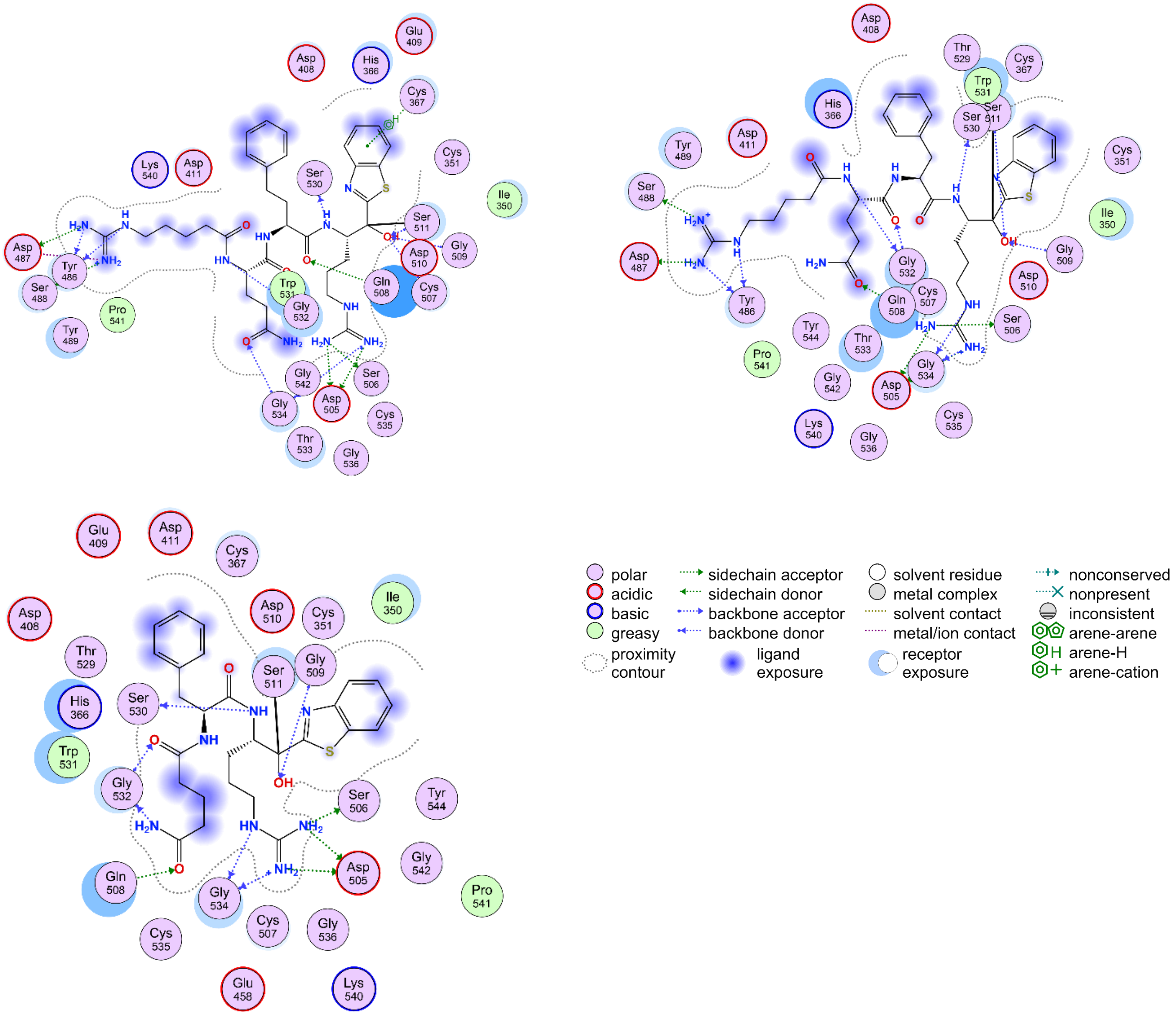

**Figure S2**. 2D interaction diagram, generated using MOE, illustrates the binding interactions between a) N-0430, b) N-0130 and c) N-0388 with TMPRSS13. The legend was shown for the type of interactions. Key interactions are as they have been listed in the tables.

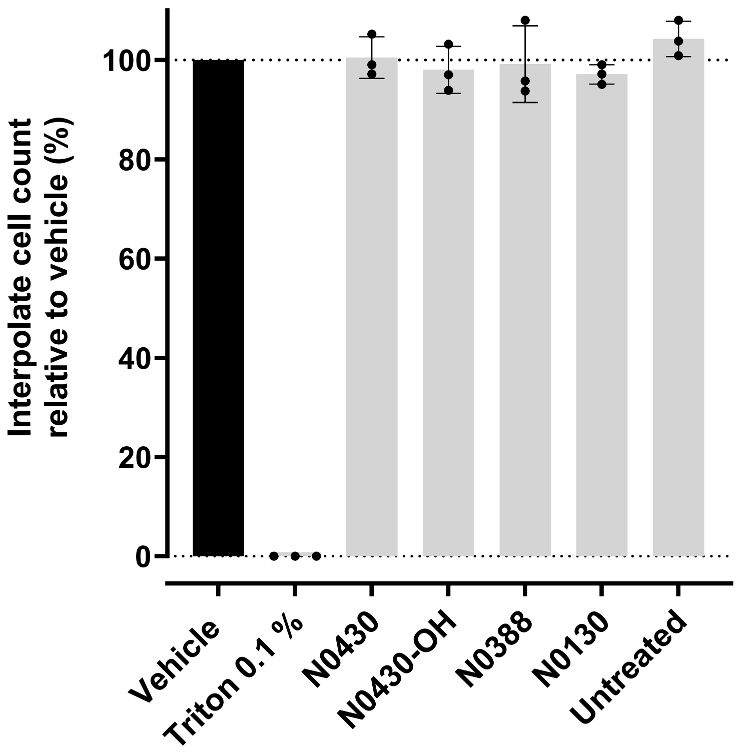

**Figure S3.** Viability test of selected compounds on Vero E6 cells. Cells were incubated for 24h with the indicated compounds at 10 µM and viability was assessed using Cell titer-glo. Compounds are non-toxic at this concentration. Each assays were performed in triplicates, three times. Results are background corrected and presented as the mean viability (%) compared to vehicle treated (DMSO 0.01 %) condition ± standard deviation (SD). Triton X-100 0.1% was used as a toxicity control.

**Table S3. Tables of interactions between TMPRSS13 and compounds.**

| Amino acids | Type | N-430 | | N-0130 | | N-0388 | |
|---|---|---|---|---|---|---|---|
| | | Energy (kcal/mol) | Distance (Å) | Energy (kcal/mol) | Distance (Å) | Energy (kcal/mol) | Distance (Å) |
| Asp487 | IH | -13.72 | 3.18 | -8.48 | 3.04 | - | - |
| Asp505 | IH | -34.69 | 2.95 | -36.02 | 2.96 | -37.91 | 2.92 |
| Cys367 | A | -0.6 | 4.25 | - | - | - | - |
| Cys507 | H | -1.7 | 2.82 | - | - | - | - |
| Gln508 | H | -2.3 | 3.01 | -2.4 | 3.15 | -4.5 | 2.78 |
| Gly509 | H | -1.4 | 3.22 | -1.7 | 2.89 | -2.1 | 3.02 |
| Gly532 | H | -8.4 | 2.89 | -9.1 | 2.98 | -7.0 | 2.99 |
| Gly534 | H | -8.1 | 2.93 | -4.6 | 3.12 | -9.3 | 2.77 |
| Ser488 | H | -2.5 | 3 | -3.8 | 2.92 | - | - |
| Ser506 | H | -0.9 | 3.07 | -3.4 | 2.83 | -2.9 | 2.9 |
| Ser511 | CH | -0.4 | 2.13 | 0.2 | 2.14 | - | - |
| | C | - | - | - | - | 1 | 1.41 |
| Ser530 | H | -4.1 | 2.82 | -3.2 | 2.94 | -2.3 | 3.1 |
| Tyr486 | H | -8.3 | 2.9 | -9.8 | 2.78 | - | - |

Table illustrates the diverse array of *N-0430*/TMPRSS13 interactions, showcasing the essential amino acids in maintaining affinity and inhibition. It also highlights aberrant *N-0130*/TMPRSS13 interactions and delves into the intricate landscape of *N-0388*/TMPRSS13 interactions. Binding energy and distant values were shown for each pair interaction and were sorted form the lowest one. Type H stands for Hydrogen bond and C for covalent bond and IH for ionic hydrogen bond. MOE is used through 'contact' tool to provide the tables.

**Table S4. Tables of interactions between matriptase and compounds.**

| Amino acids | Type | N-0130 | | N-0388 | |
|---|---|---|---|---|---|
| | | Energy (kcal/mol) | Distance (Å) | Energy (kcal/mol) | Distance (Å) |
| Asp799 | IH | -31,41 | 3,04 | -30,64 | 2,99 |
| Asp828 | IH | -18,03 | 3,08 | - | - |
| | H | - | - | -5,2 | 2,9 |
| Gln783 | H | -10,5 | 2,82 | - | - |
| Gln802 | H | -1,3 | 2,78 | -1,3 | 2,83 |
| Gly803 | H | -1,8 | 2,93 | -2,1 | 2,95 |
| Gly827 | H | -4,8 | 2,92 | -0,7 | 3,46 |
| Gly829 | H | -6,1 | 2,78 | -6,5 | 2,76 |
| Ser800 | H | -2,3 | 2,9 | -2,4 | 2,88 |
| Ser805 | C | 1 | 1,4 | - | - |
| | CH | - | - | -0,1 | 2,18 |
| Ser825 | H | -2,4 | 3,23 | -1,6 | 3,38 |
| Tyr755 | H | -2,9 | 2,72 | -3,5 | 2,72 |

Table illustrates the diverse array of N-0130/matriptase interactions in, showcasing the essential amino acids in maintaining affinity and inhibition. Aberrant N-0388/matriptase interactions are also described. Binding energy and distant values were shown for each pair interaction and were sorted form the lowest one. Type H stands for Hydrogen bond and C for covalent bond and IH for ionic hydrogen bond. MOE is used through 'contact' tool to provide the tables.

**Table S5. Accurate mass measurement (HRMS) and purity of the leads and hits.**

| COMPOUND | ANALYSIS | CHARGE | MOLECULAR FORMULA | M/Z THEORICAL | M/Z MEASURED | PURITY UPLC-MS (%) |
|---|---|---|---|---|---|---|
| N-0430 | ESI+ Q-TOF (maXis) | $[M+2H]^{2+}$ | $C_{34}H_{47}N_{11}O_5S$ | 361.6814 | 361.6812 | 98.21 |
| N-0430-OH 1 | | $[M+2H]^{2+}$ | $C_{34}H_{49}N_{11}O_5S$ | 362.6892 | 362.6891 | 99.51 |
| N-0430-OH 2 | | $[M+2H]^{2+}$ | $C_{34}H_{49}N_{11}O_5S$ | 362.6892 | 362.6880 | 99.46 |
| N-0388 | | $[M+H]^{+}$ | $C_{27}H_{33}N_7O_4S$ | 552.2388 | 552.2392 | 97.84 |
| N-0439 | | $[M+2H]^{2+}$ | $C_{31}H_{42}N_{10}O_4S$ | 326.1628 | 326.1624 | 99.77 |
| N-0182 | | $[M+2H]^{2+}$ | $C_{34}H_{46}N_{12}O_6S$ | 376.1765 | 376.1758 | 96.55 |

### UPLC characterization data of TMPRSS13 inhibitors

S3: Compound **#54**. IRAR-Kbt

Molecular formula: $C_{28}H_{45}N_{11}O_4S$

MW calculated: 631.80, m/z found: 632.40 $[M+H]^+$.

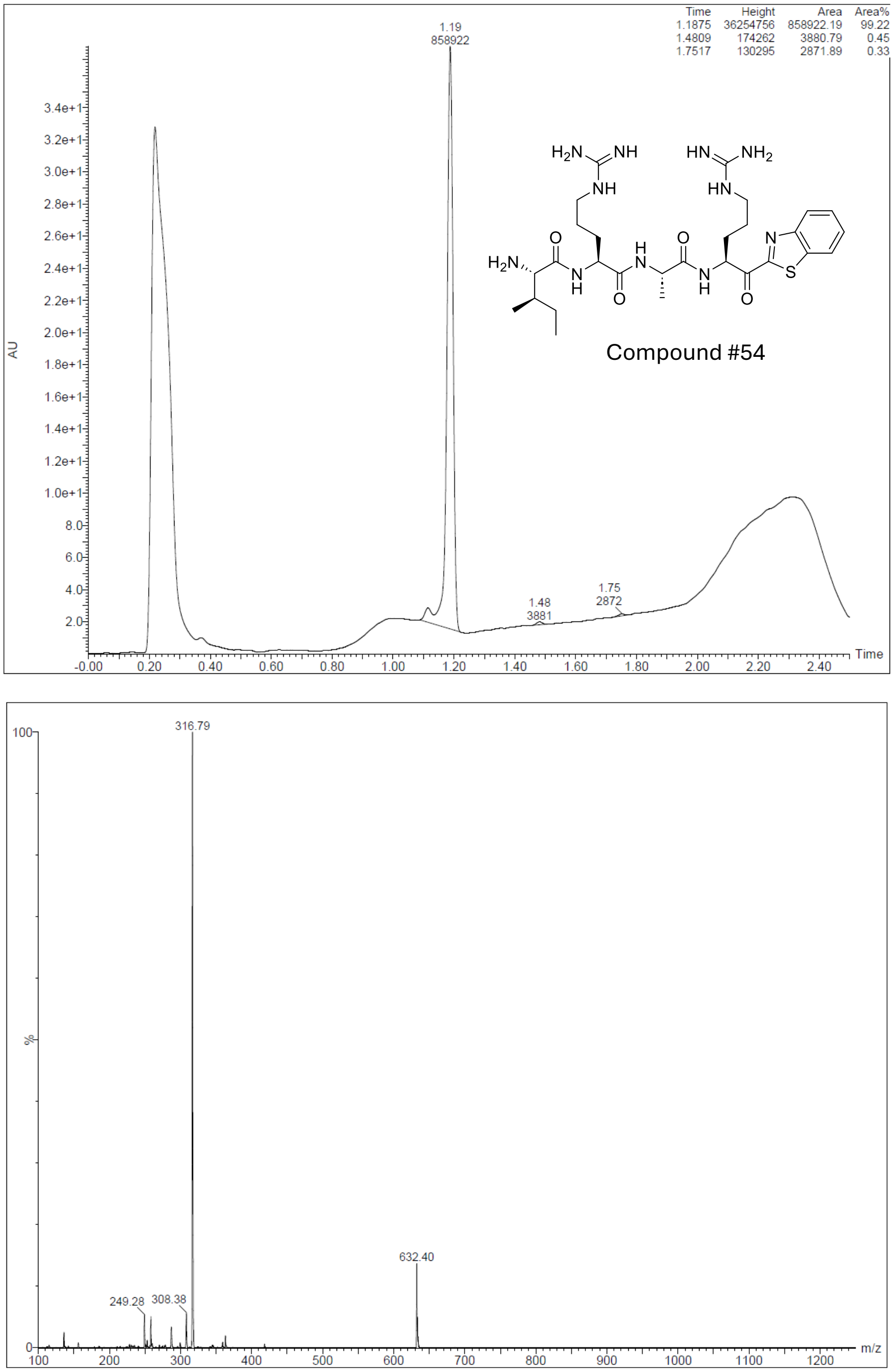

**Figure S4**. UPLC chromatogram and MS of compound #54

S4: Compound **#55**. IQAR-Kbt

Molecular formula: $C_{27}H_{41}N_9O_5S$

MW calculated: 603.74, m/z found: 604.44 $[M+H]^+$.

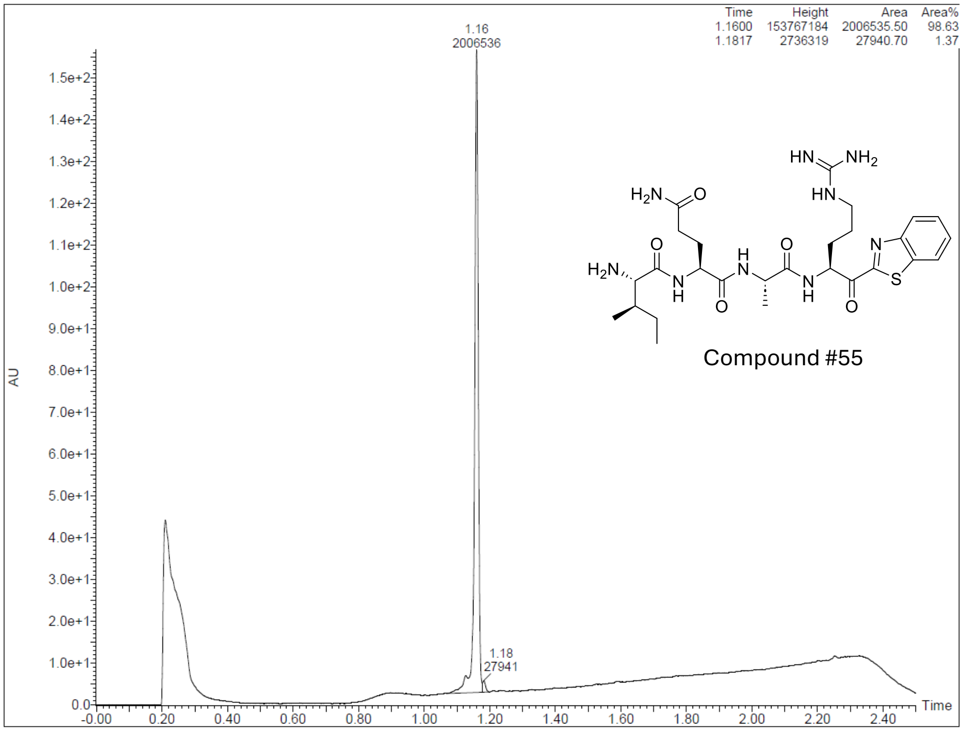

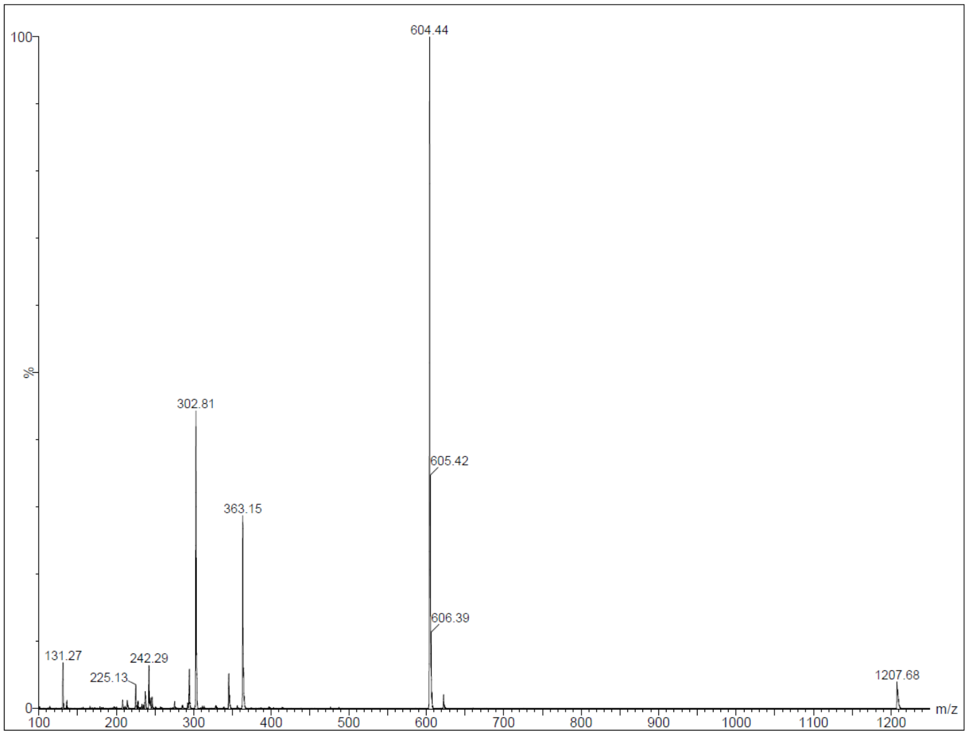

**Figure S5.** UPLC chromatogram and MS compound #55

S5: Compound **#56**. RQDR-Kbt

Molecular formula: $C_{28}H_{42}N_{12}O_7S$

MW calculated: 690.78, m/z found: 691.33 $[M+H]^+$.

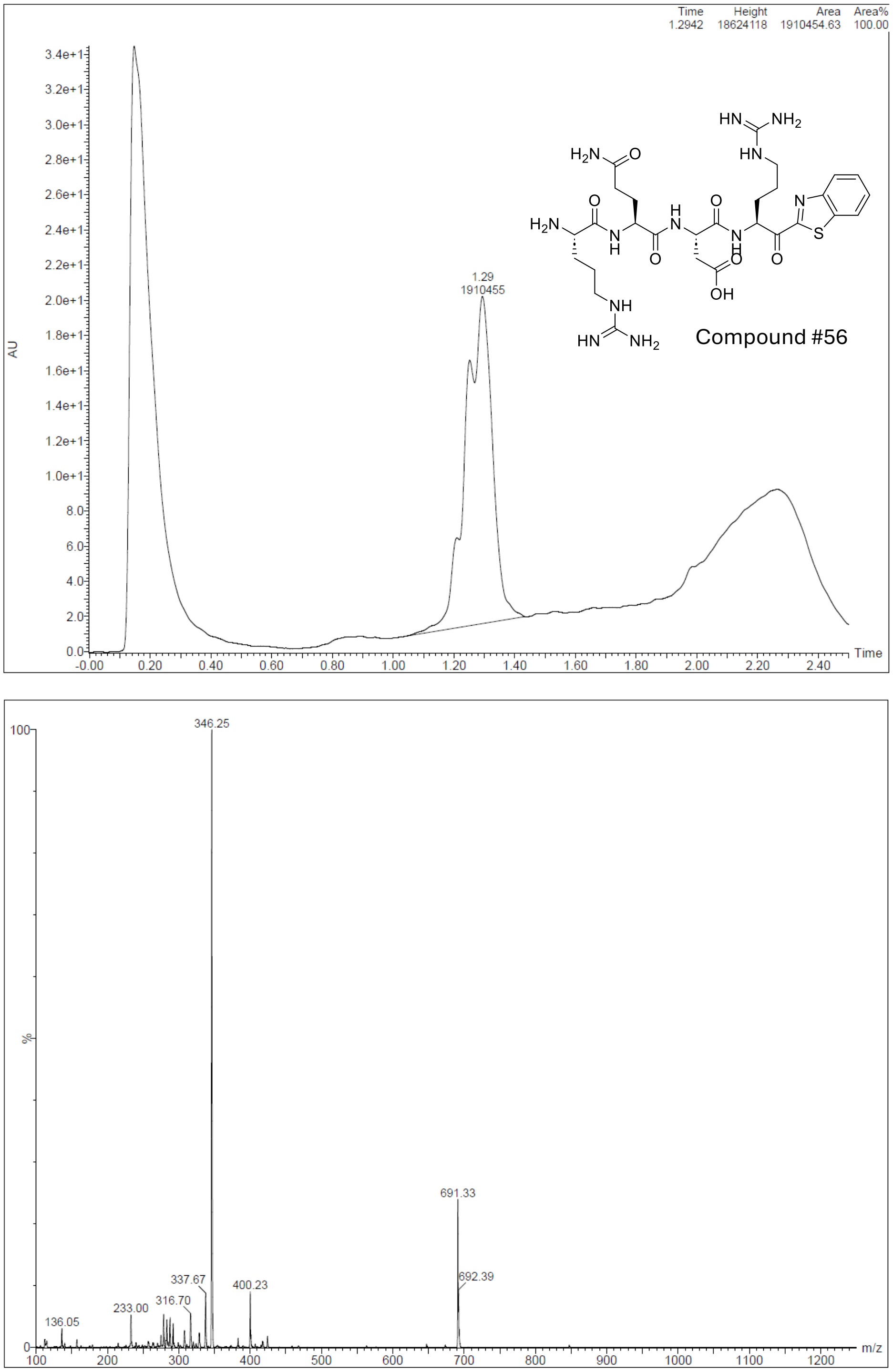

**Figure S6.** UPLC chromatogram and MS of compound #56

S6: Compound **#57**. RLAR-Kbt

Molecular formula: $C_{28}H_{45}N_{11}O_4S$

MW calculated: 631.80, m/z found: 632.60 $[M+H]^+$.

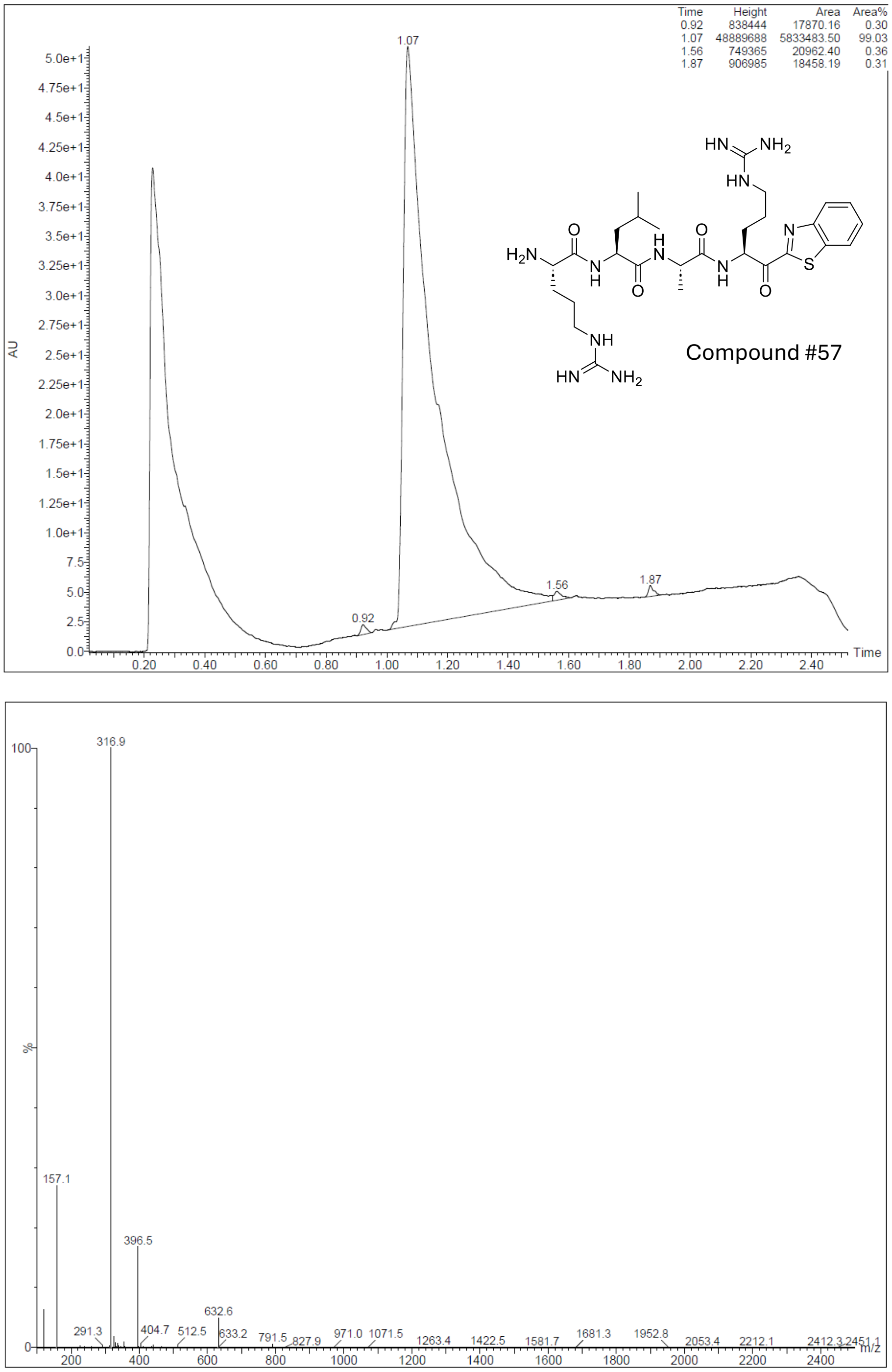

**Figure S7.** UPLC chromatogram and MS of compound #57

S7: Compound **#58**. RQF($CF_3$)R-Kbt

Molecular formula: $C_{34}H_{45}N_{12}O_5S$

MW calculated: 790.87, m/z found: 791.50 $[M+H]^+$.

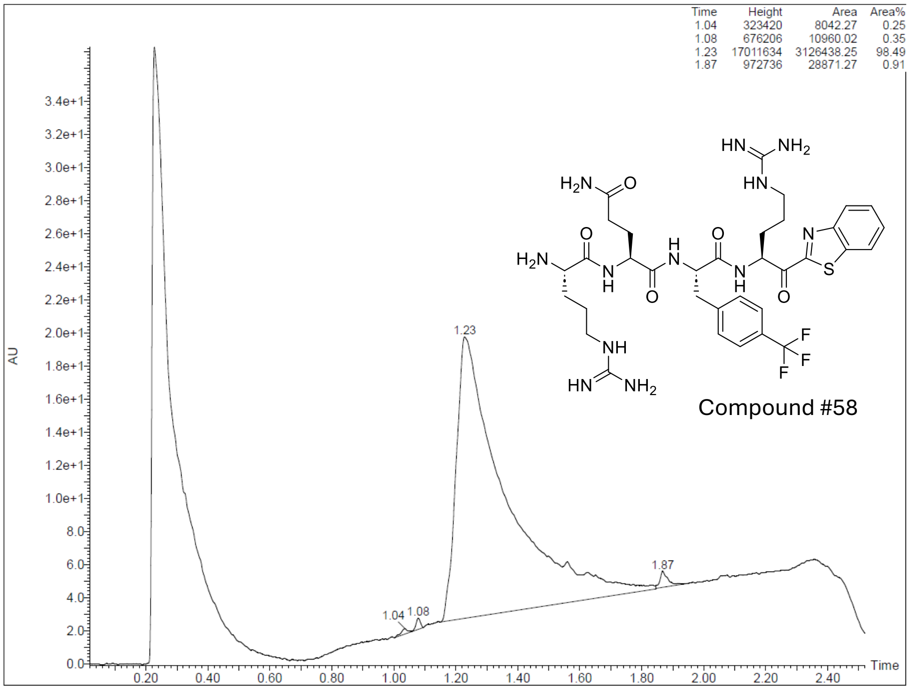

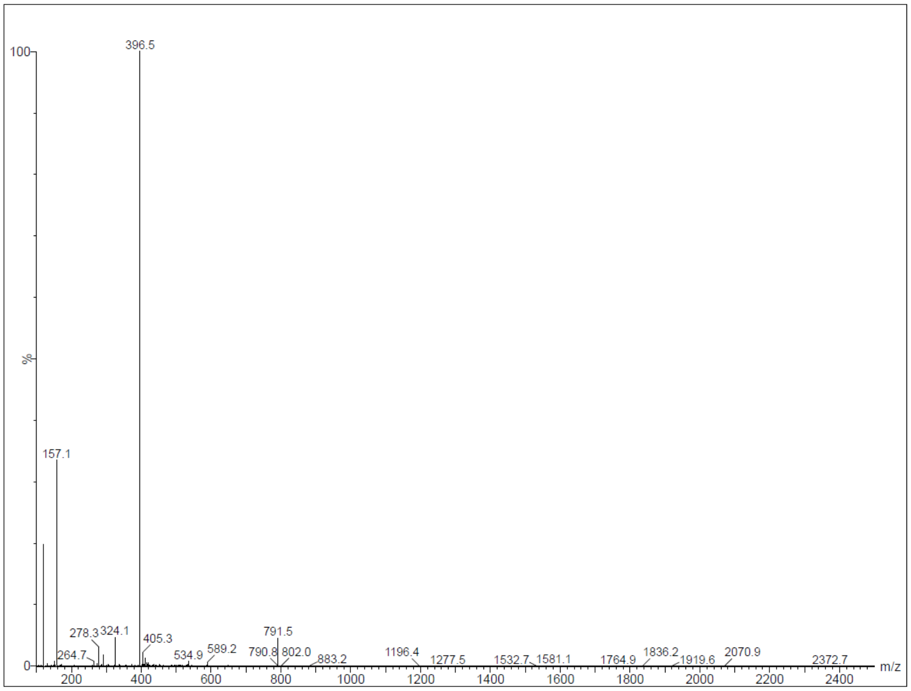

**Figure S8.** UPLC chromatogram and MS of compound #58

S8: Compound **#59**. RQAN(Arg)-Kbt

Molecular formula: $C_{27}H_{42}N_{12}O_5S$

MW calculated: 646.77, m/z found: 647.60 $[M+H]^+$.

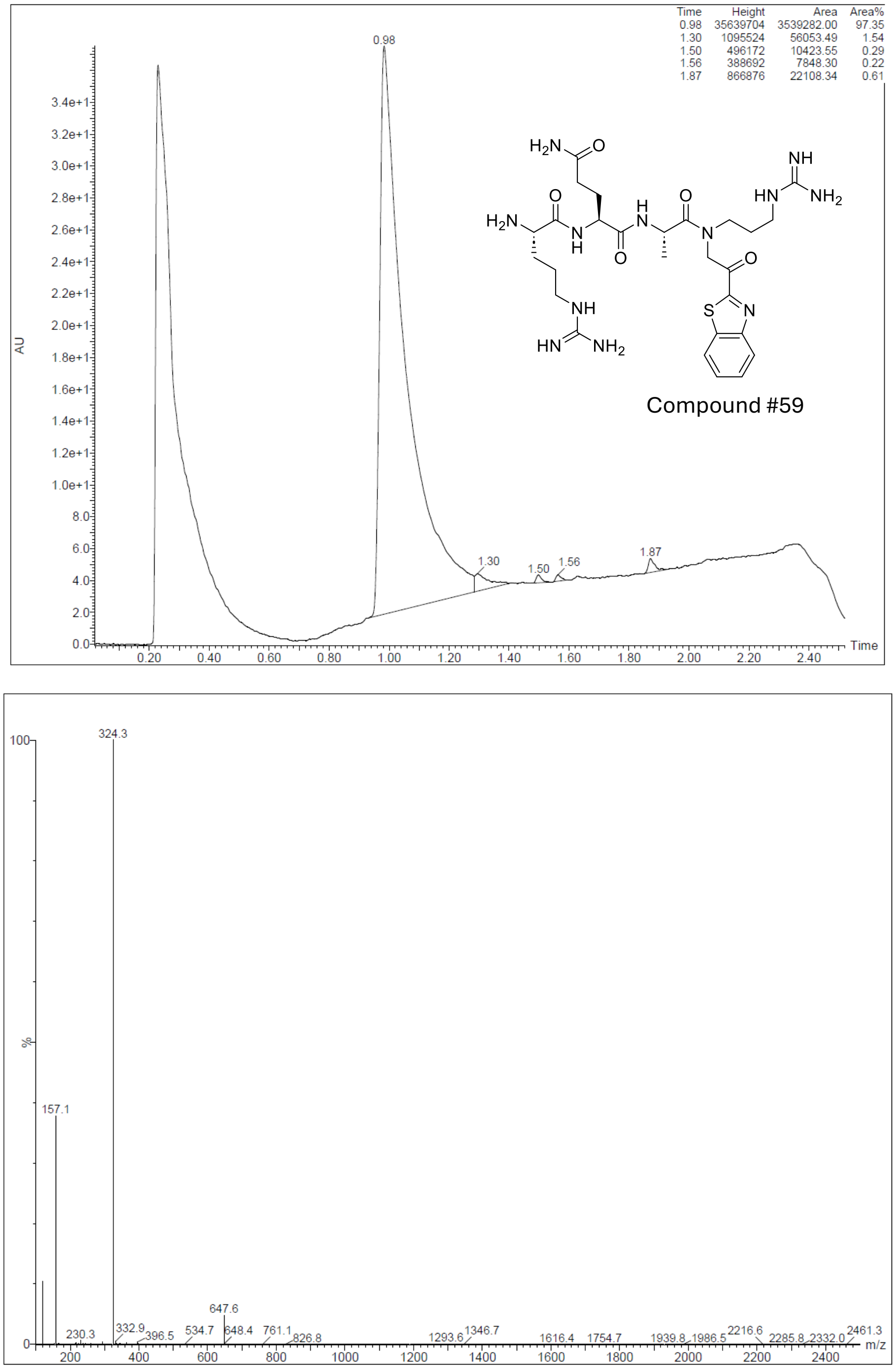

**Figure S9.** UPLC chromatogram and MS of compound #59

S9: Compound **N-0439 (#60)**. (H)-RAFR-Kbt

Molecular formula: $C_{31}H_{42}N_{10}O_4S$

MW calculated: 650.80, m/z found: 651.34 $[M+H]^+$.

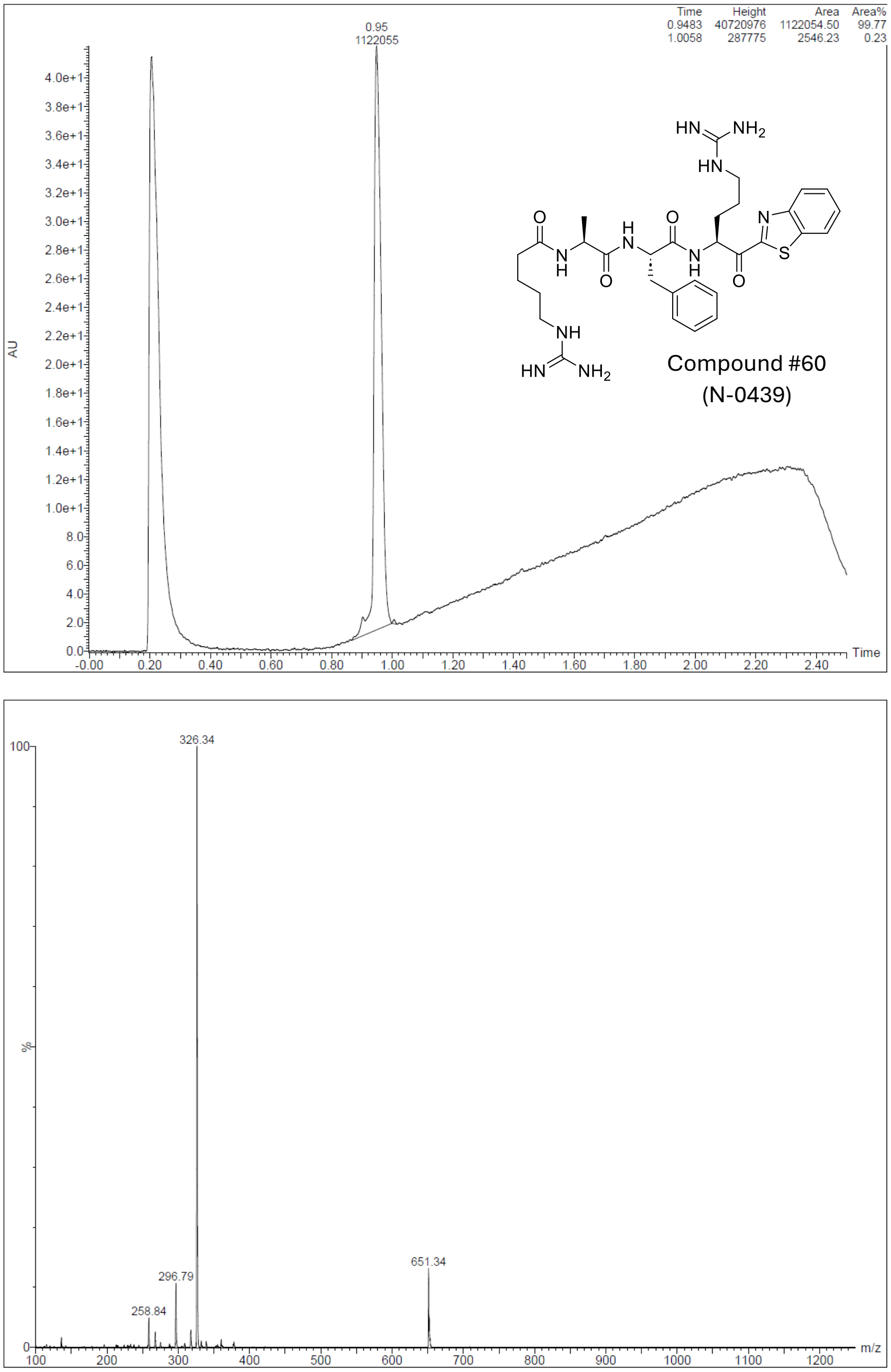

**Figure S10.** UPLC chromatogram and MS of N-0439 (compound #60)

S10: Compound **#61**. (H)-SQFR-Kbt

Molecular formula: $C_{30}H_{38}N_8O_6S$

MW calculated: 638.74, m/z found: 639.39 $[M+H]^+$.

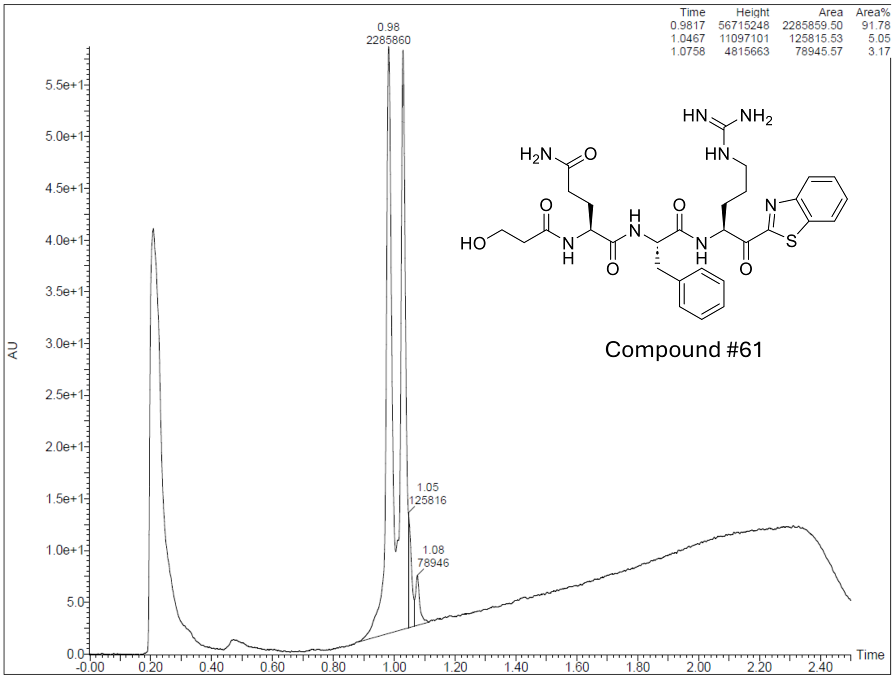

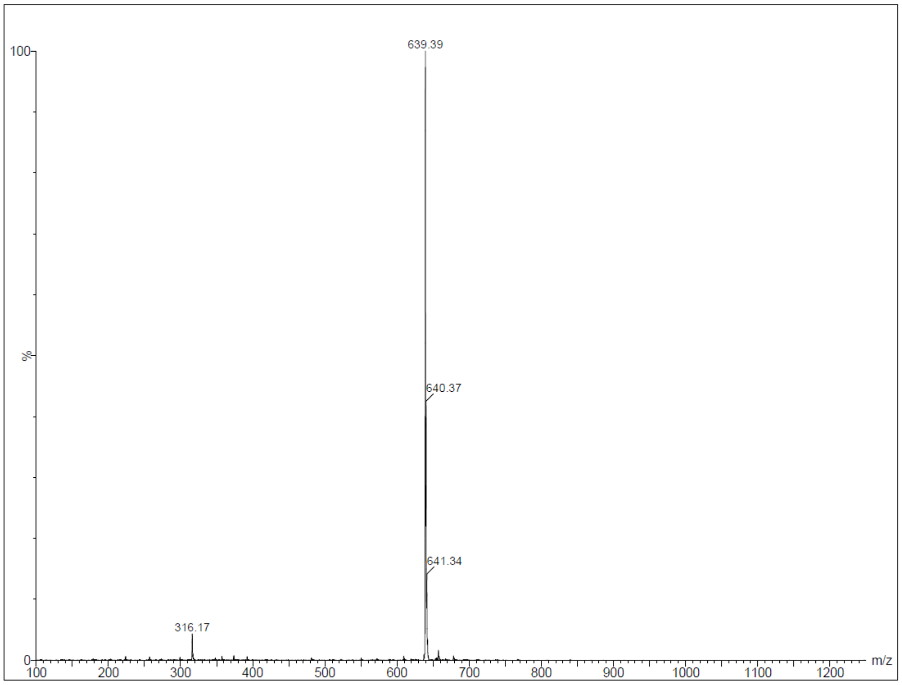

**Figure S11.** UPLC chromatogram and MS of compound #61

S11: Compound **#62**. (H)-RQSR-Kbt

Molecular formula: $C_{27}H_{41}N_{11}O_6S$

MW calculated: 647.76, m/z found: 648.42 $[M+H]^+$.

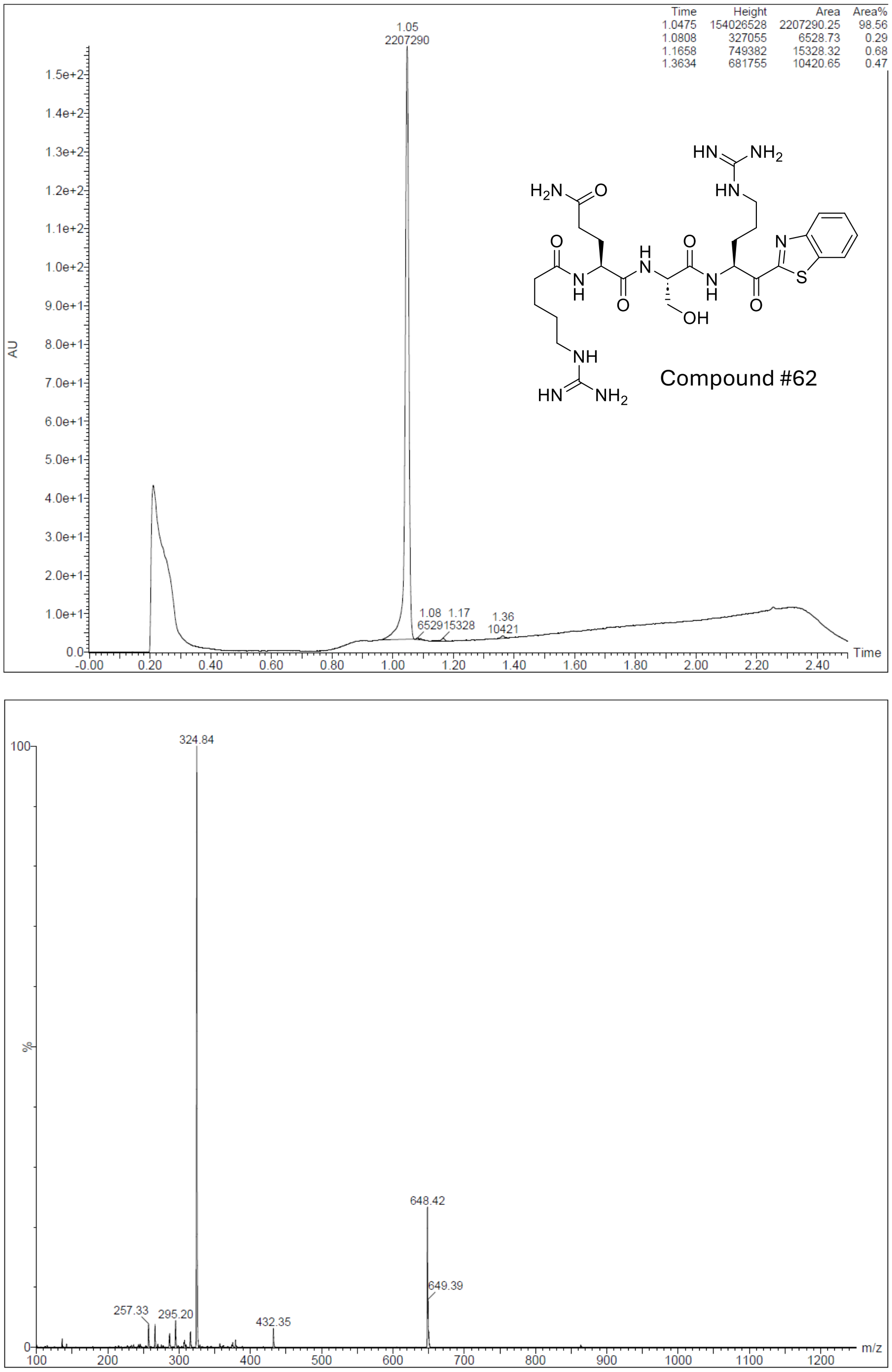

**Figure S12.** UPLC chromatogram and MS of compound #62

S12: Compound **#63**. (H)-RQBpaR-Kbt

Molecular formula: $C_{40}H_{49}N_{11}O_6S$

MW calculated: 811.96, m/z found: 812.44 $[M+H]^+$.

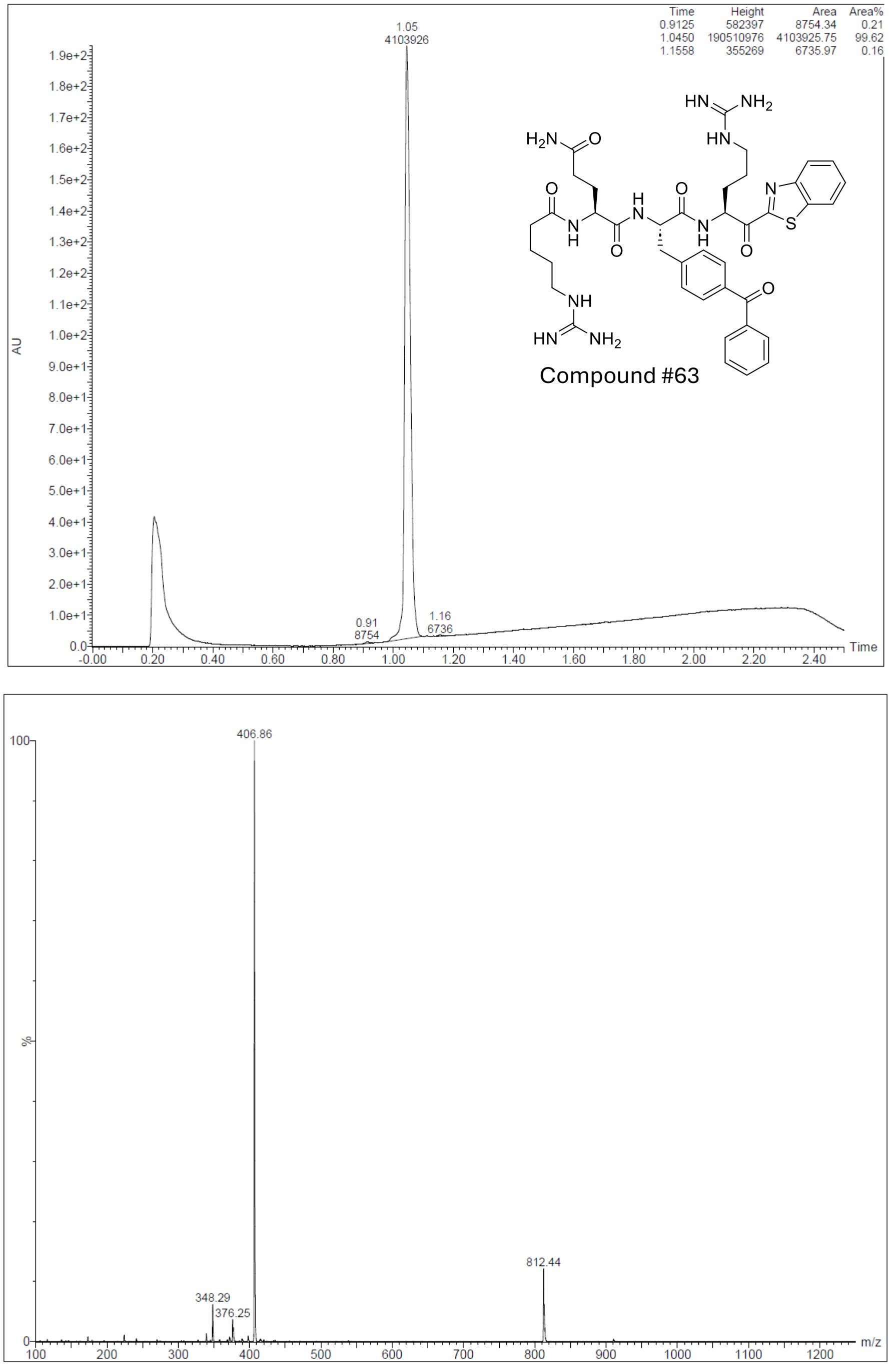

**Figure S13.** UPLC chromatogram and MS of compound #63

S13: Compound **N-0182 (#64)**. PhCO-RQAR-Kbt

Molecular formula: $C_{34}H_{46}N_{12}O_6S$

MW calculated: 750.88, m/z found: 751.47 $[M+H]^+$.

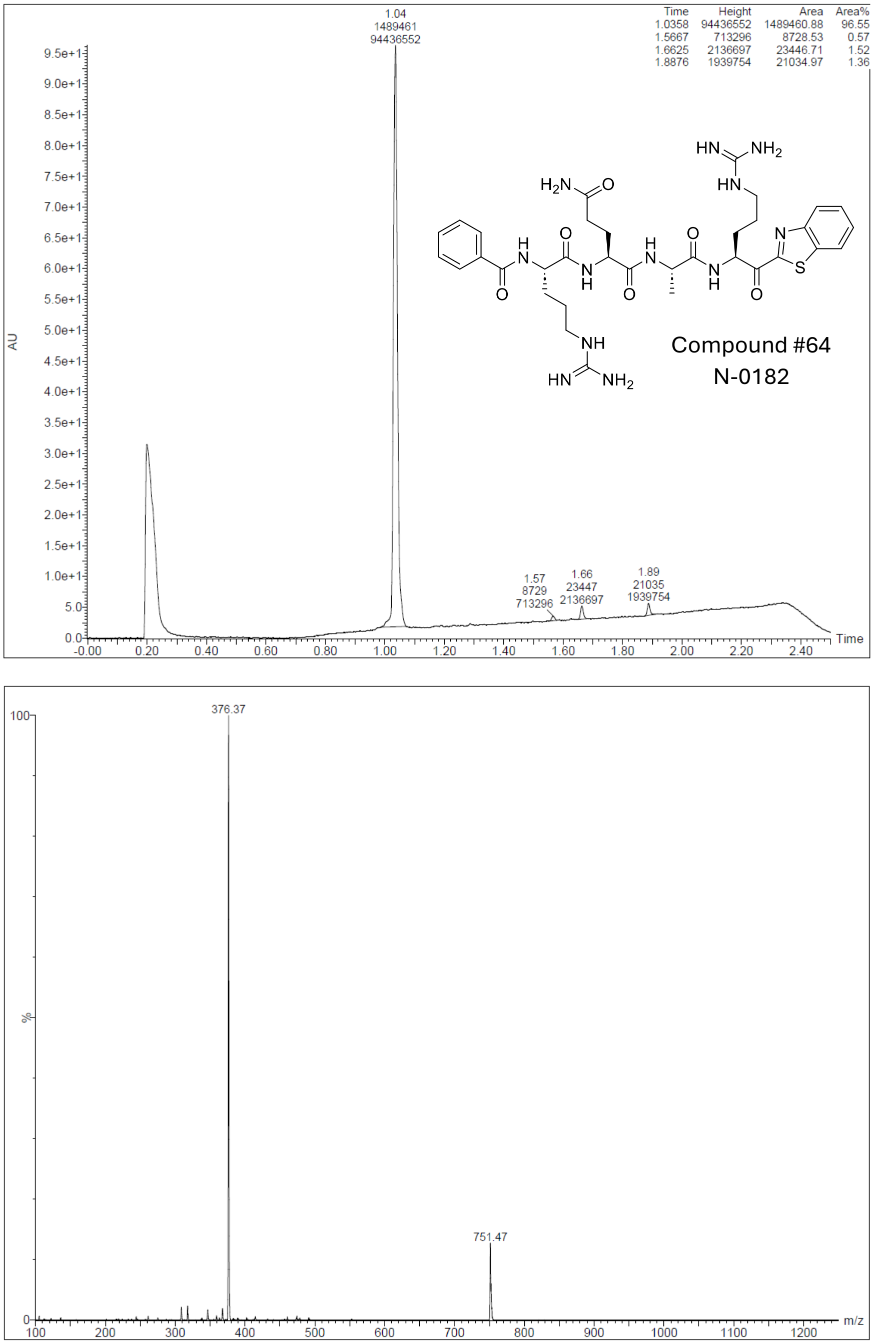

**Figure S14.** UPLC chromatogram and MS of N-0182 (compound #64)

S14: Compound #**65**. [GABA-E]FR-Kbt

Molecular formula: $C_{31}H_{38}N_8O_5S$

MW calculated: 634.76, m/z found: 635.41 $[M+H]^+$.

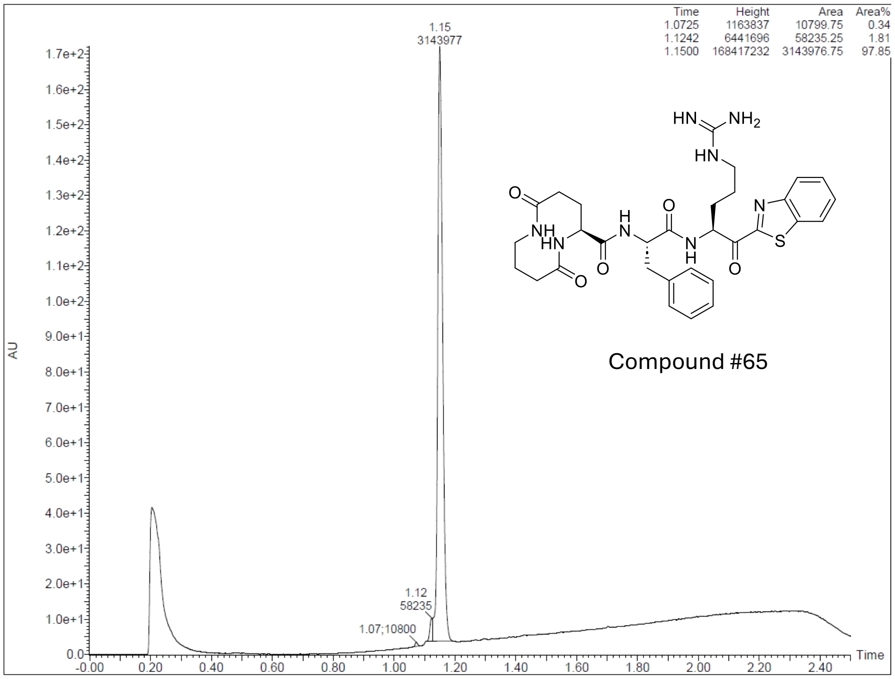

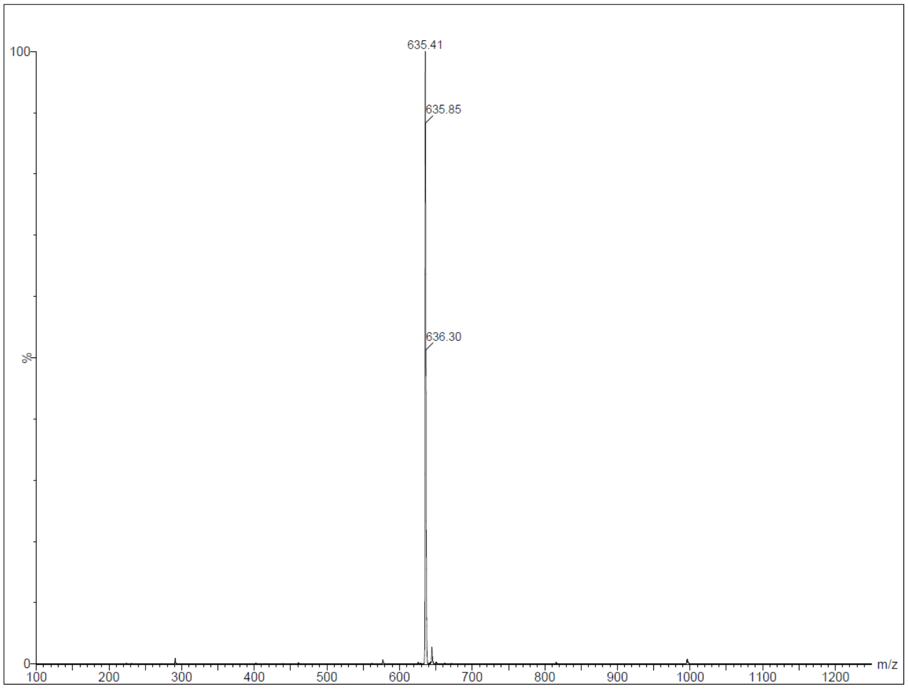

**Figure S15.** UPLC chromatogram and MS of compound #65

S15: Compound **N-0430 (#66)**. (H)-RQhFR-Kbt

Molecular formula: $C_{34}H_{47}N_{11}O_5S$

MW calculated: 721.88, m/z found: 722.44 $[M+H]^+$.

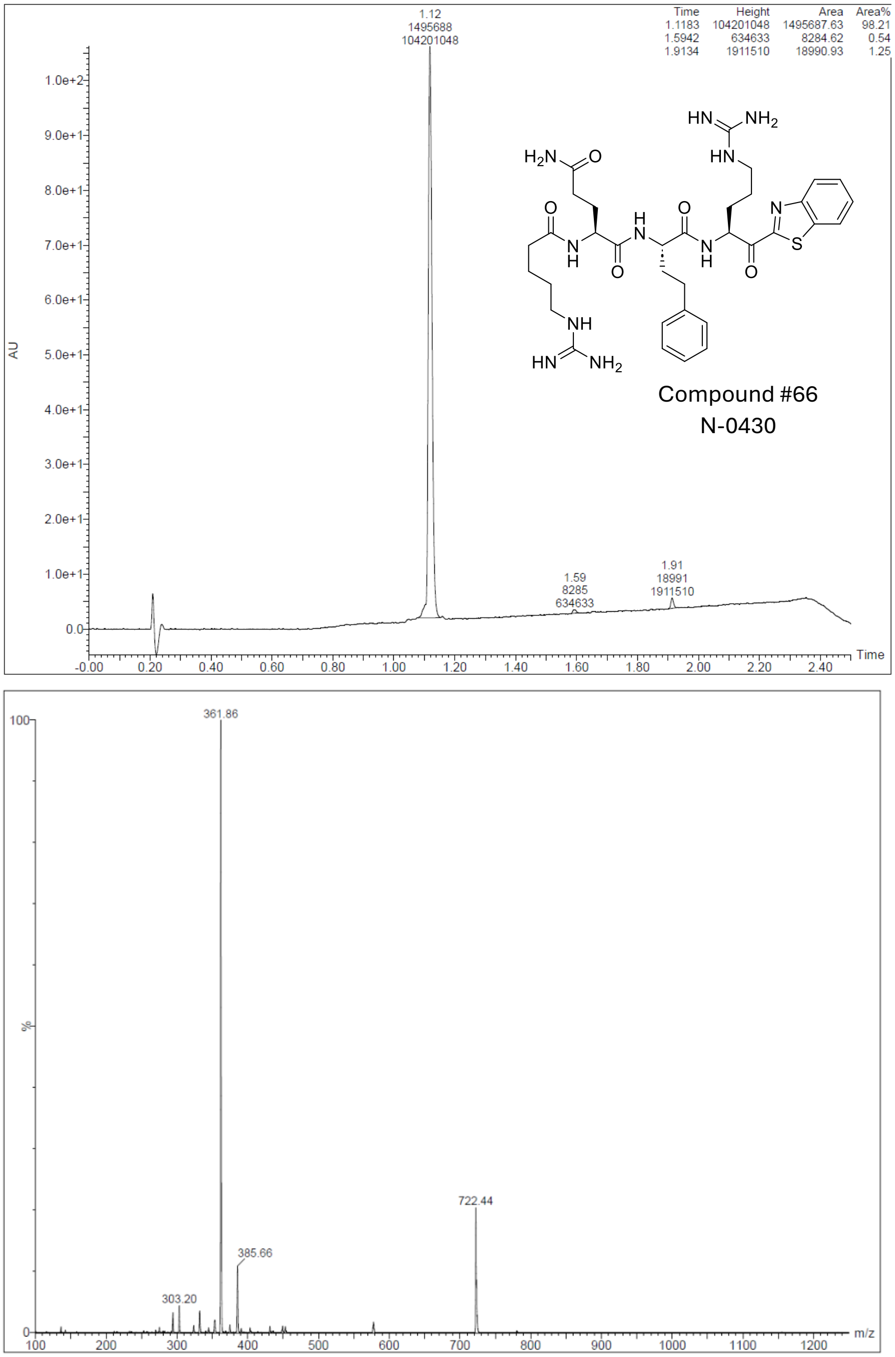

**Figure S16.** UPLC chromatogram and MS of N-0430 (compound #66)

S16: Compound **N-0388 (#67)**. (H)-QFR-Kbt

Molecular formula: $C_{27}H_{33}N_7O_4S$

MW calculated: 551.67, m/z found: 552.50 $[M+H]^+$.

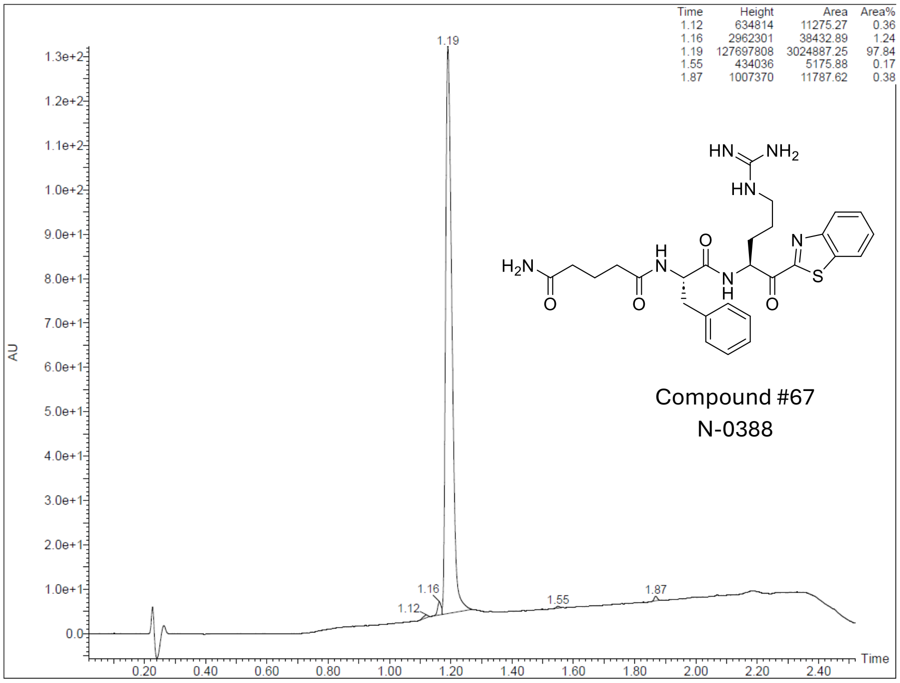

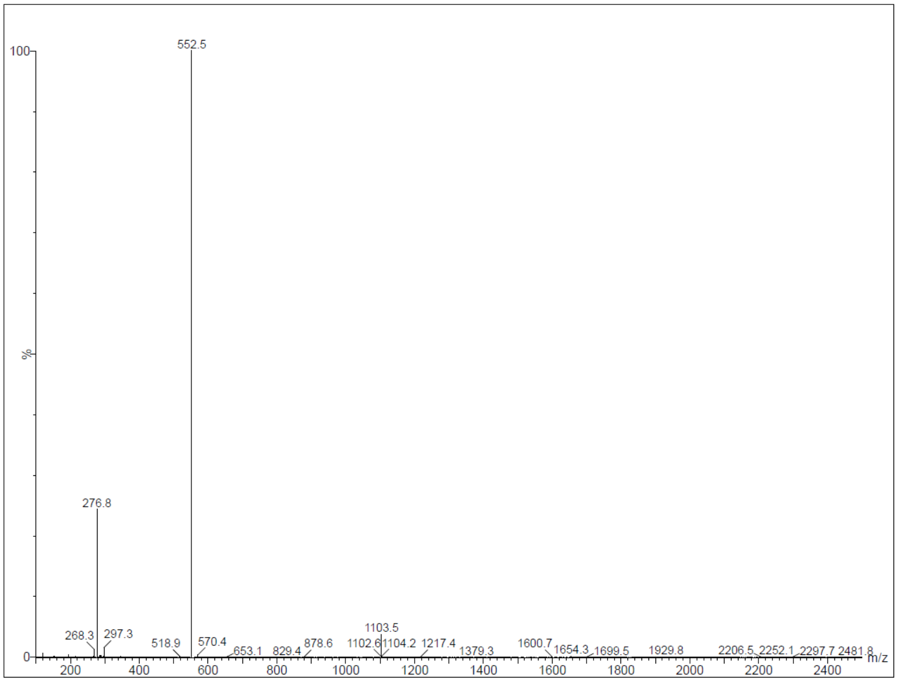

**Figure S17.** UPLC chromatogram and MS of N-0388 (compound #67)

S17: Compound **N-0430-OH-dia 1 (#68)**. (H)-RQhFR-(OH)bt

Molecular formula: $C_{34}H_{49}N_{11}O_5S$

MW calculated: 723.90, m/z found: 724.57 $[M+H]^+$.

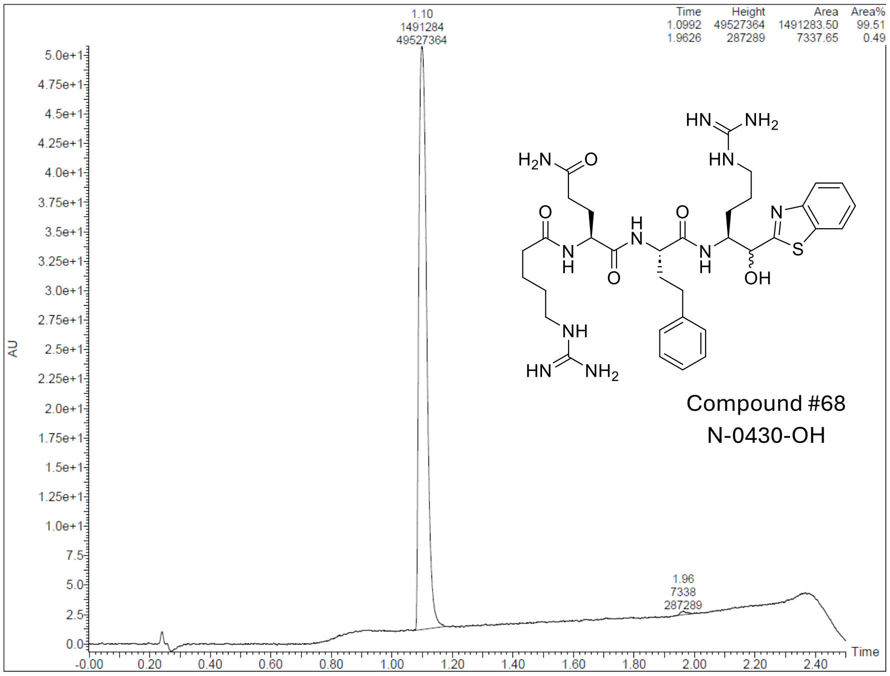

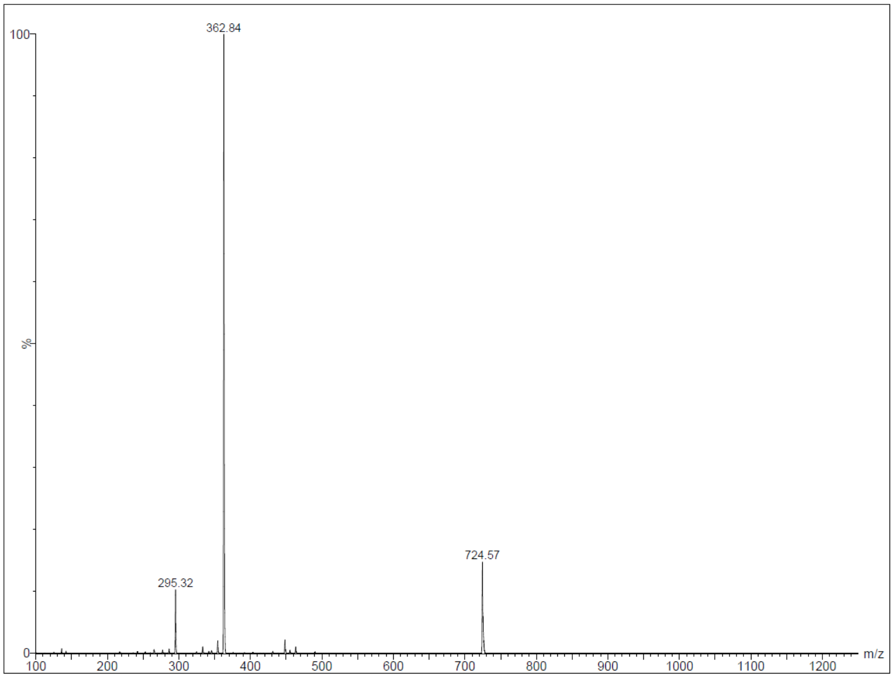

**Figure S18.** UPLC chromatogram and MS of N-0430-OH (dia 1) (compound #68)

S18: Compound **N-0430-OH-dia 2 (#68)**. (H)-RQhFR-(OH)bt

Molecular formula: $C_{34}H_{49}N_{11}O_5S$

MW calculated: 723.90, m/z found: 724.57 $[M+H]^+$.

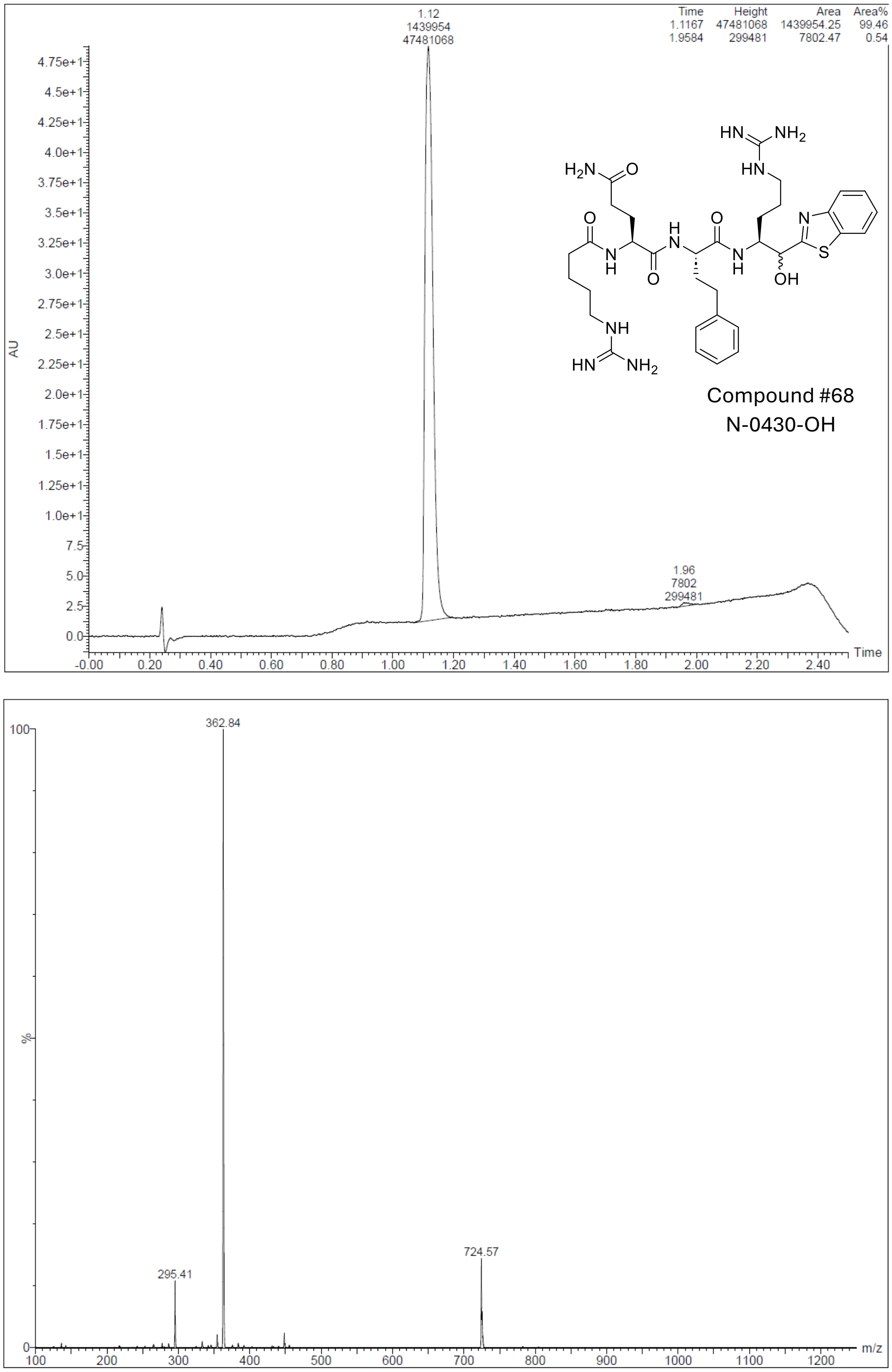

**Figure S19.** UPLC chromatogram and MS of N-0430-OH (dia 2) (compound #68)

### NMR characterisation data of TMPRSS13 inhibitors

#### Compound #66

$^{1}H$

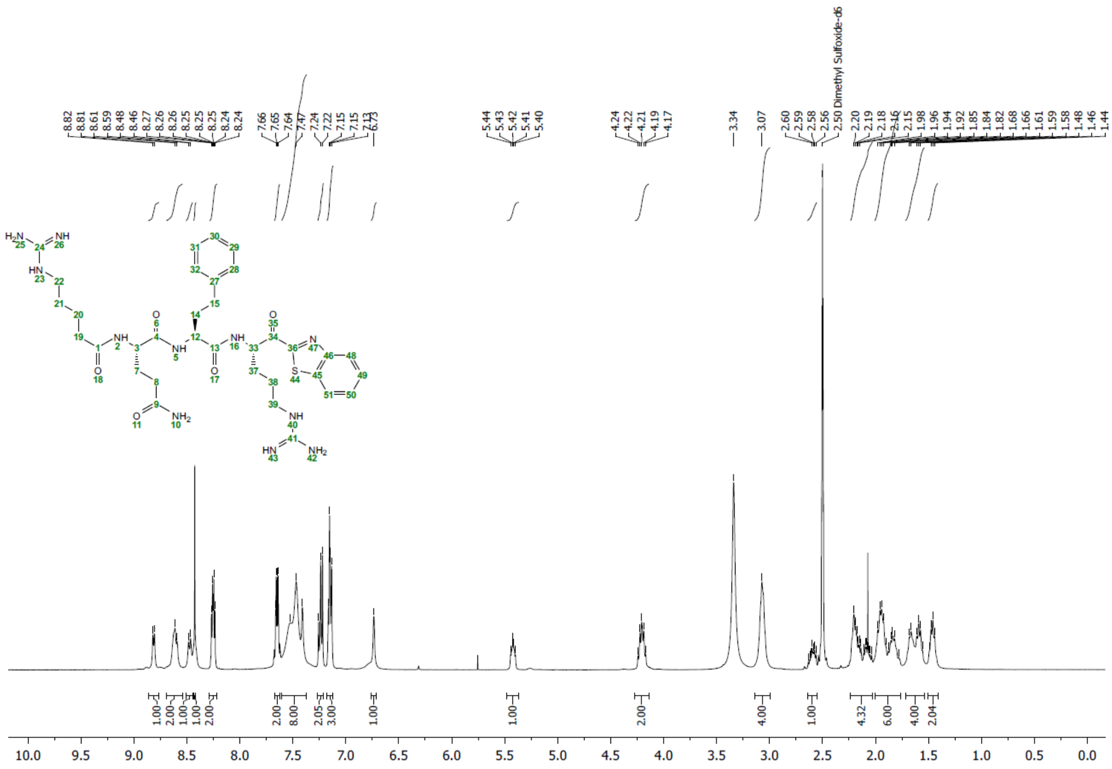

**Figure S20.** $^{1}H$ NMR of compound **N-0430 (#66)**. (H)-RQhFR-Kbt

$^{1}H$ NMR (400 MHz, DMSO) δ 8.81 (d, *J* = 5.8 Hz, 1H), 8.60 (d, *J* = 8.0 Hz, 2H), 8.47 (d, *J* = 7.7 Hz, 1H), 8.43 (s, 1H), 8.28 – 8.22 (m, *J* = 7.7, 4.0, 1.4 Hz, 2H), 7.69 – 7.61 (m, 2H), 7.59 – 7.37 (m, *J* = 23.5 Hz, 8H), 7.27 – 7.20 (m, 2H), 7.18 – 7.12 (m, *J* = 7.2, 5.1 Hz, 3H), 6.73 (s, 1H), 5.47 – 5.37 (m, 1H), 4.26 – 4.15 (m, 2H), 3.14 – 3.00 (m, 4H), 2.66 – 2.55 (m, 1H), 2.25 – 2.02 (m, 4H), 2.01 – 1.75 (m, 6H), 1.73 – 1.53 (m, 4H), 1.47 (dd, *J* = 13.0, 6.3 Hz, 2H).

$^{13}C$

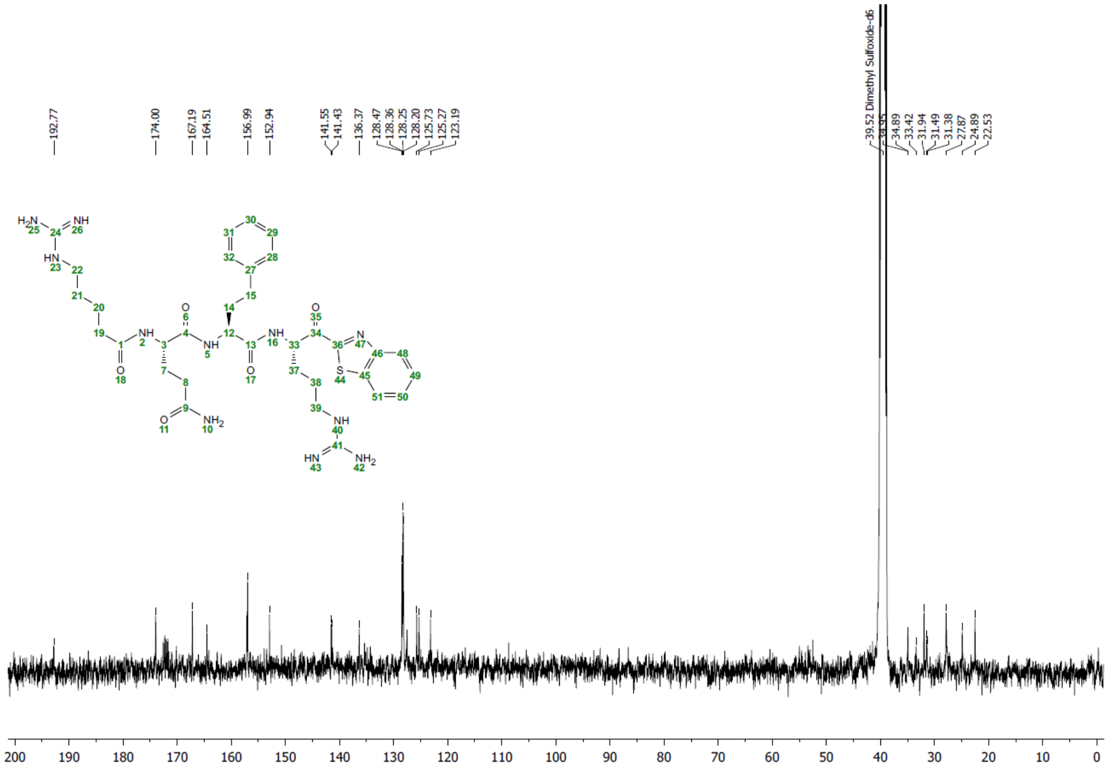

**Figure S21.** $^{13}C$ NMR of compound **N-0430 (#66)**. (H)-RQhFR-Kbt

$^{13}C$ NMR (101 MHz, DMSO) δ 192.77, 174.00, 167.19, 164.51, 156.99, 152.94, 141.55, 141.43, 136.37, 128.47, 128.36, 128.25, 128.20, 125.73, 125.27, 123.19, 39.52, 34.95, 34.89, 33.42, 31.94, 31.49, 31.38, 27.87, 24.89, 22.53.

### Compound #67

[1]H

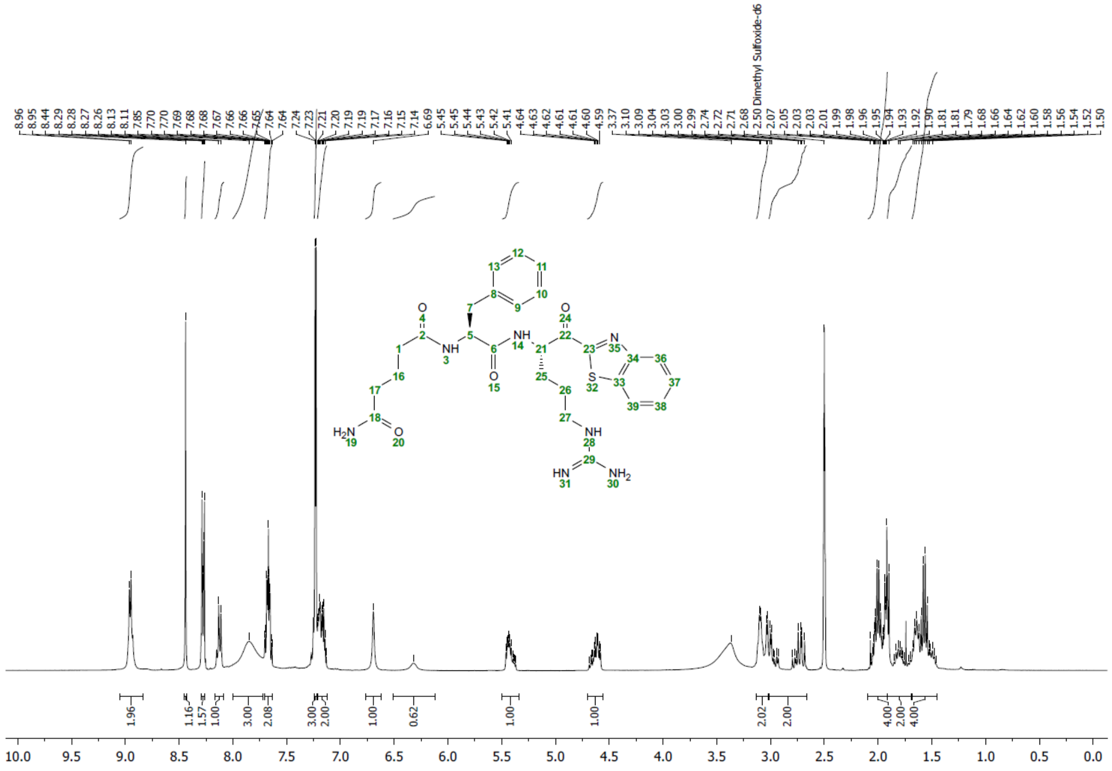

**Figure S22.** [1]H NMR of compound **N-0388 (#67)**. (H)-QFR-Kbt

1H NMR (400 MHz, DMSO) δ 8.95 (d, J = 6.1 Hz, 2H), 8.44 (s, 1H), 8.28 (dd, J = 6.6, 2.8 Hz, 1H), 8.12 (d, J = 8.3 Hz, 1H), 7.85 (s, 3H), 7.71 – 7.62 (m, 2H), 7.23 (d, J = 4.3 Hz, 3H), 7.17 (ddd, J = 13.0, 7.6, 4.7 Hz, 2H), 6.69 (s, 1H), 6.32 (s, 1H), 5.49 – 5.35 (m, 1H), 4.70 – 4.56 (m, J = 10.4, 8.8, 4.9 Hz, 1H), 3.06 (dd, J = 26.1, 4.8 Hz, 2H), 3.01 – 2.66 (m, 2H), 2.10 – 1.91 (m, 4H), 1.91 – 1.69 (m, 2H), 1.68 – 1.44 (m, 4H).

$^{13}C$

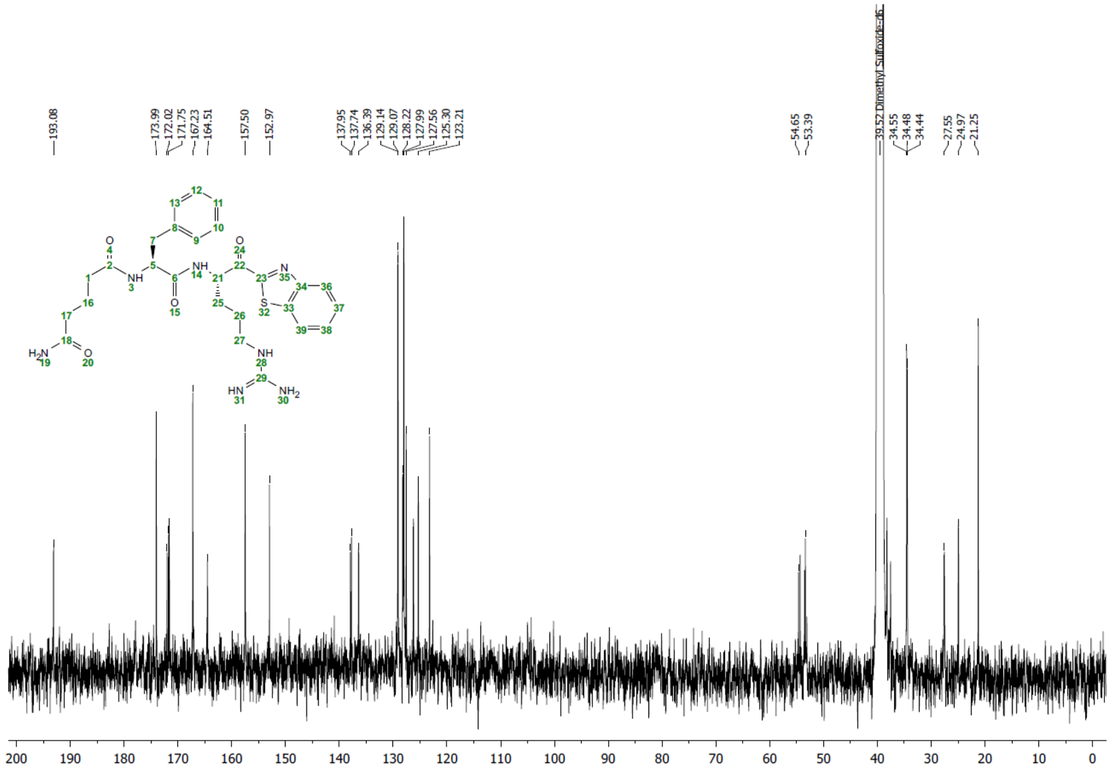

**Figure S23.** $^{13}C$ NMR of compound **N-0388 (#67)**. (H)-QFR-Kbt

13C NMR (101 MHz, DMSO) δ 193.08, 173.99, 172.02, 171.75, 167.23, 164.51, 157.50, 152.97, 137.95, 137.74, 136.39, 129.14, 129.07, 128.22, 127.99, 127.56, 125.30, 123.21, 54.65, 53.39, 39.52, 34.55, 34.48, 34.44, 27.55, 24.97, 21.25.

**Compound #68 (dia 1)**

$^{1}$H

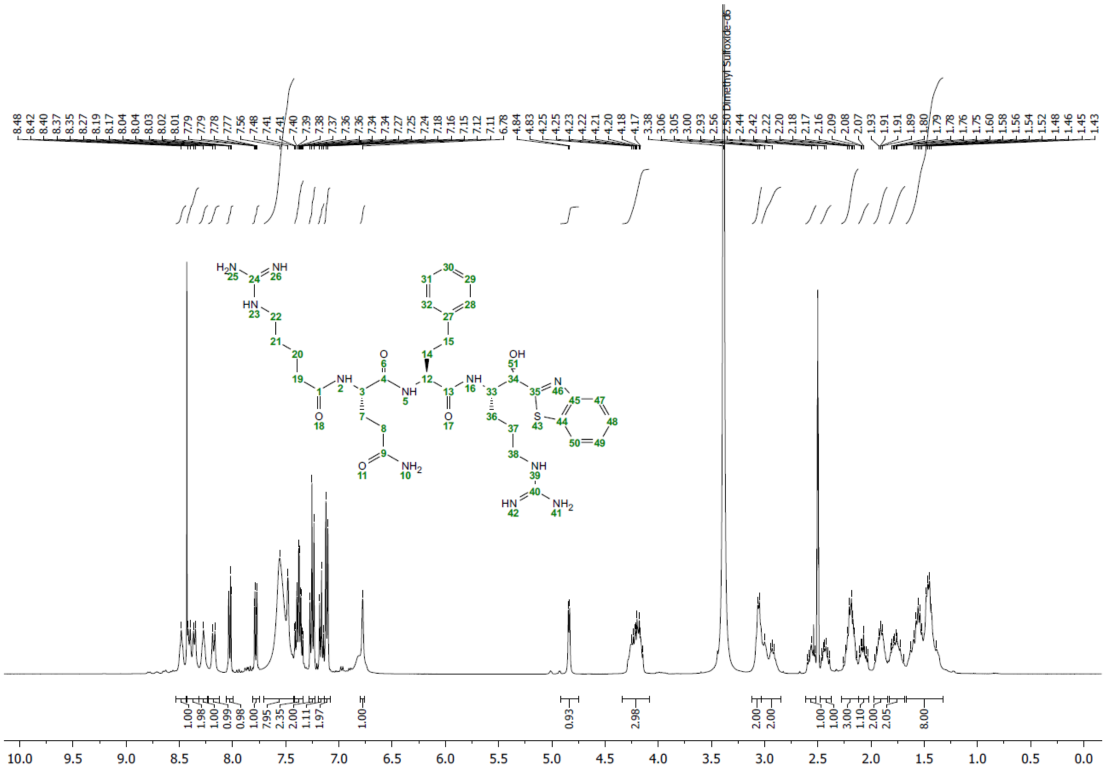

**Figure S24.** $^{1}$H NMR of compound **N-0430-OH-dia 1 (#68)**. (H)-RQhFR-(OH)bt

$^{1}$H NMR (400 MHz, DMSO) δ 8.48 (s, 1H), 8.38 (dd, *J* = 19.8, 7.7 Hz, 2H), 8.27 (s, 1H), 8.18 (d, *J* = 8.8 Hz, 1H), 8.06 – 8.00 (m, 1H), 7.78 (dd, *J* = 7.1, 1.7 Hz, 1H), 7.56 (s, *J* = 31.0 Hz, 8H), 7.38 (tdd, *J* = 8.7, 7.3, 1.5 Hz, 2H), 7.25 (t, *J* = 7.3 Hz, 2H), 7.16 (t, *J* = 7.3 Hz, 1H), 7.11 (d, *J* = 7.0 Hz, 2H), 6.78 (s, 1H), 4.84 (d, *J* = 4.8 Hz, 1H), 4.33 – 4.08 (m, 3H), 3.06 (d, *J* = 5.3 Hz, 2H), 3.02 – 2.84 (m, *J* = 20.1, 15.2 Hz, 2H), 2.57 (dd, *J* = 19.1, 9.6 Hz, 1H), 2.47 – 2.37 (m, 1H), 2.28 – 2.12 (m, 3H), 2.11 – 2.02 (m, 1H), 1.97 – 1.84 (m, 2H), 1.76 (ddd, *J* = 27.9, 17.3, 9.8 Hz, 2H), 1.66 – 1.32 (m, 8H).

$^{13}C$

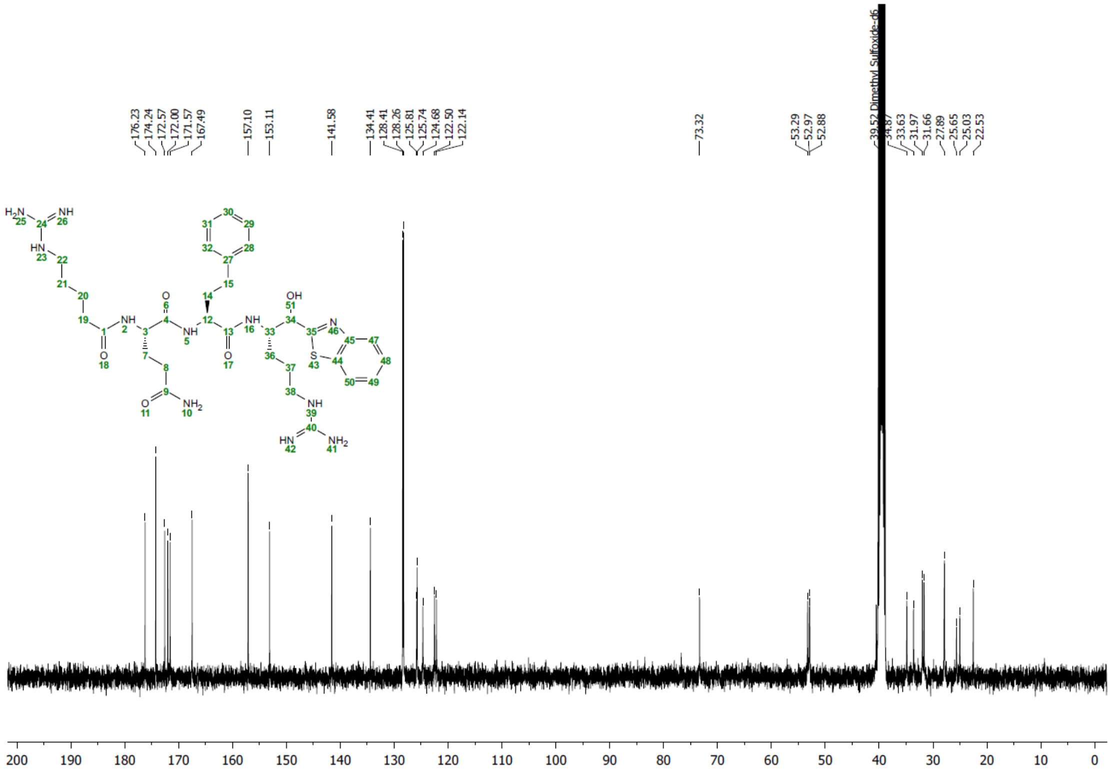

**Figure S25.** $^{13}C$ NMR of compound **N-0430-OH-dia 1 (#68)**. (H)-RQhFR-(OH)bt

13C NMR (101 MHz, DMSO) δ 176.23, 174.24, 172.57, 172.00, 171.57, 167.49, 157.10, 153.11, 141.58, 134.41, 128.41, 128.26, 125.81, 125.74, 124.68, 122.50, 122.14, 73.32, 53.29, 52.97, 52.88, 39.52, 34.87, 33.63, 31.97, 31.66, 27.89, 25.65, 25.03, 22.53.

**Compound #68 (dia 2)**

$^1$H

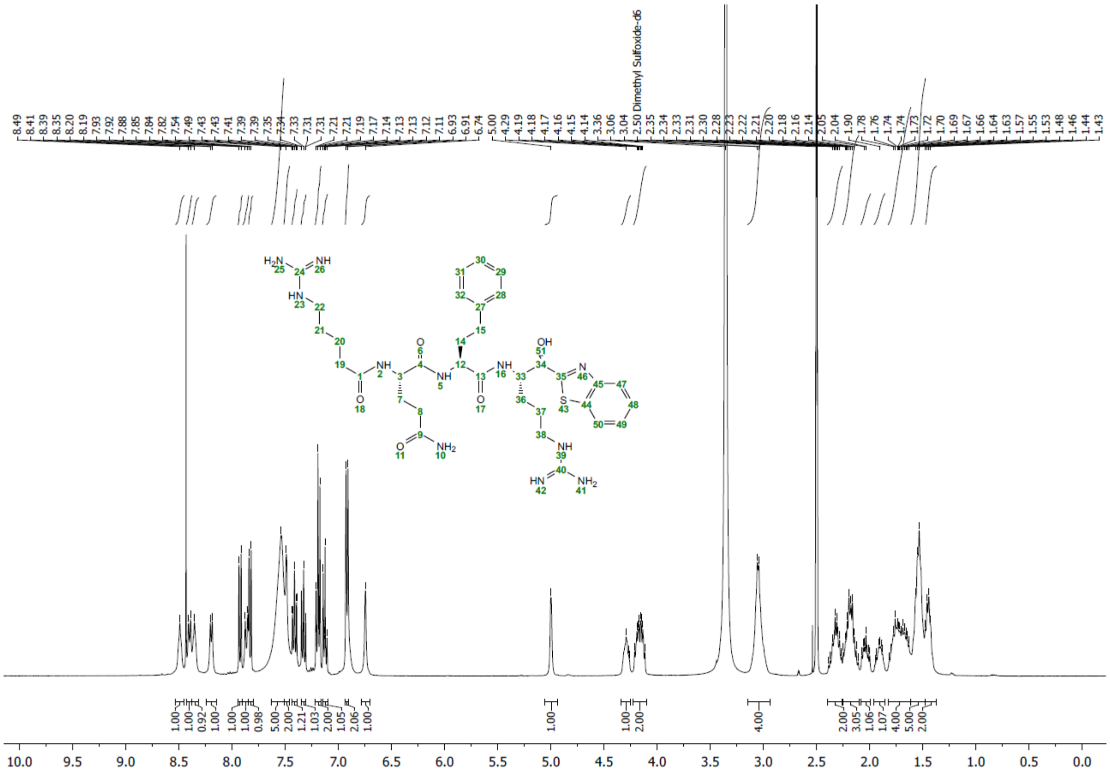

**Figure S26.** $^1$H NMR of compound **N-0430-OH-dia 2 (#68)**. (H)-RQhFR-(OH)bt

$^1$H NMR (400 MHz, DMSO) δ 8.49 (s, 1H), 8.40 (d, *J* = 7.8 Hz, 1H), 8.35 (s, 1H), 8.20 (d, *J* = 7.2 Hz, 1H), 7.92 (d, *J* = 7.7 Hz, 1H), 7.86 (d, *J* = 9.1 Hz, 1H), 7.83 (d, *J* = 8.1 Hz, 1H), 7.54 (s, 5H), 7.49 (s, 2H), 7.44 – 7.39 (m, 1H), 7.35 – 7.30 (m, 1H), 7.19 (t, *J* = 7.2 Hz, 2H), 7.15 – 7.10 (m, 1H), 6.92 (d, *J* = 7.0 Hz, 2H), 6.74 (s, 1H), 5.00 (s, 1H), 4.34 – 4.24 (m, 1H), 4.16 (ddd, *J* = 15.7, 13.7, 8.2 Hz, 2H), 3.05 (d, *J* = 5.0 Hz, 4H), 2.40 – 2.26 (m, 2H), 2.25 – 2.10 (m, 3H), 2.08 – 1.99 (m, 1H), 1.96 – 1.86 (m, 1H), 1.83 – 1.62 (m, 4H), 1.53 (dd, *J* = 20.6, 15.4 Hz, 5H), 1.47 – 1.37 (m, 2H).

$^{13}C$

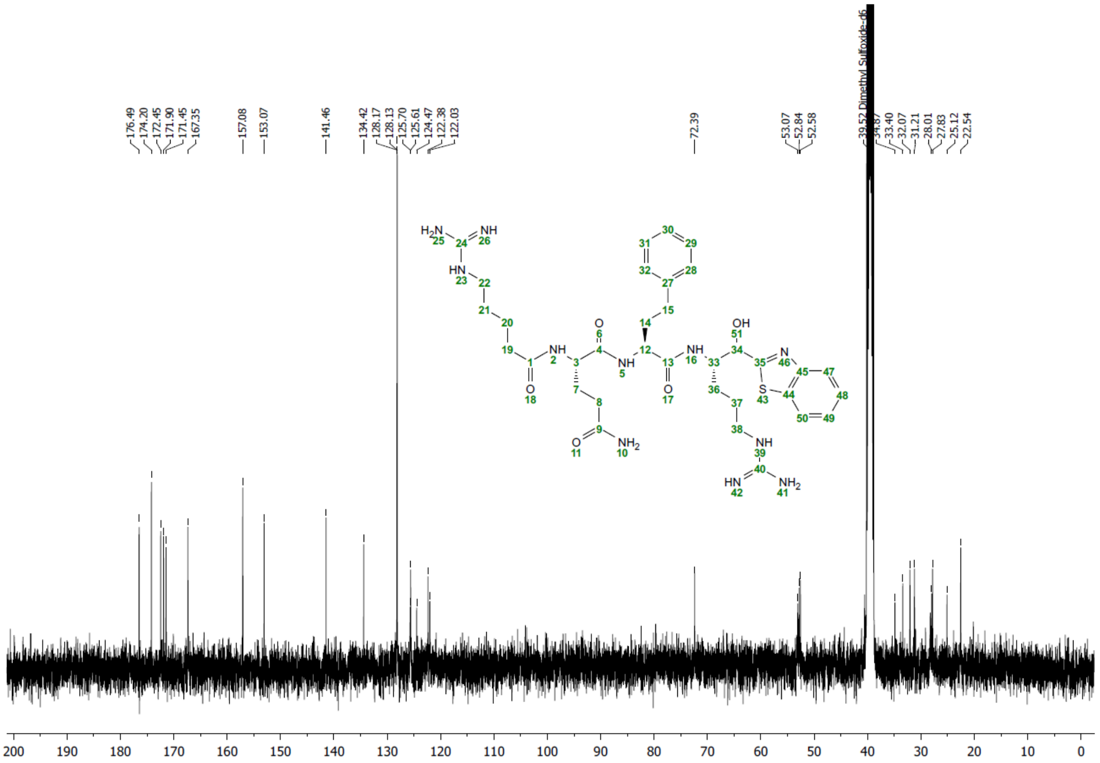

**Figure S27.** $^{13}C$ NMR of compound **N-0430-OH-dia 2 (#68)**. (H)-RQhFR-(OH)bt

13C NMR (101 MHz, DMSO) δ 176.49, 174.20, 172.45, 171.90, 171.45, 167.35, 157.08, 153.07, 141.46, 134.42, 128.17, 128.13, 125.70, 125.61, 124.47, 122.38, 122.03, 72.39, 53.07, 52.84, 52.58, 39.52, 34.87, 33.40, 32.07, 31.21, 28.01, 27.83, 25.12, 22.54.

### Accurate mass measurement (HRMS), isotopic profile of TMPRSS13 inhibitors, and detailed synthesis of compound 65, N-0430, N-0430-OH and N-0388

*Synthesis of compound 65*

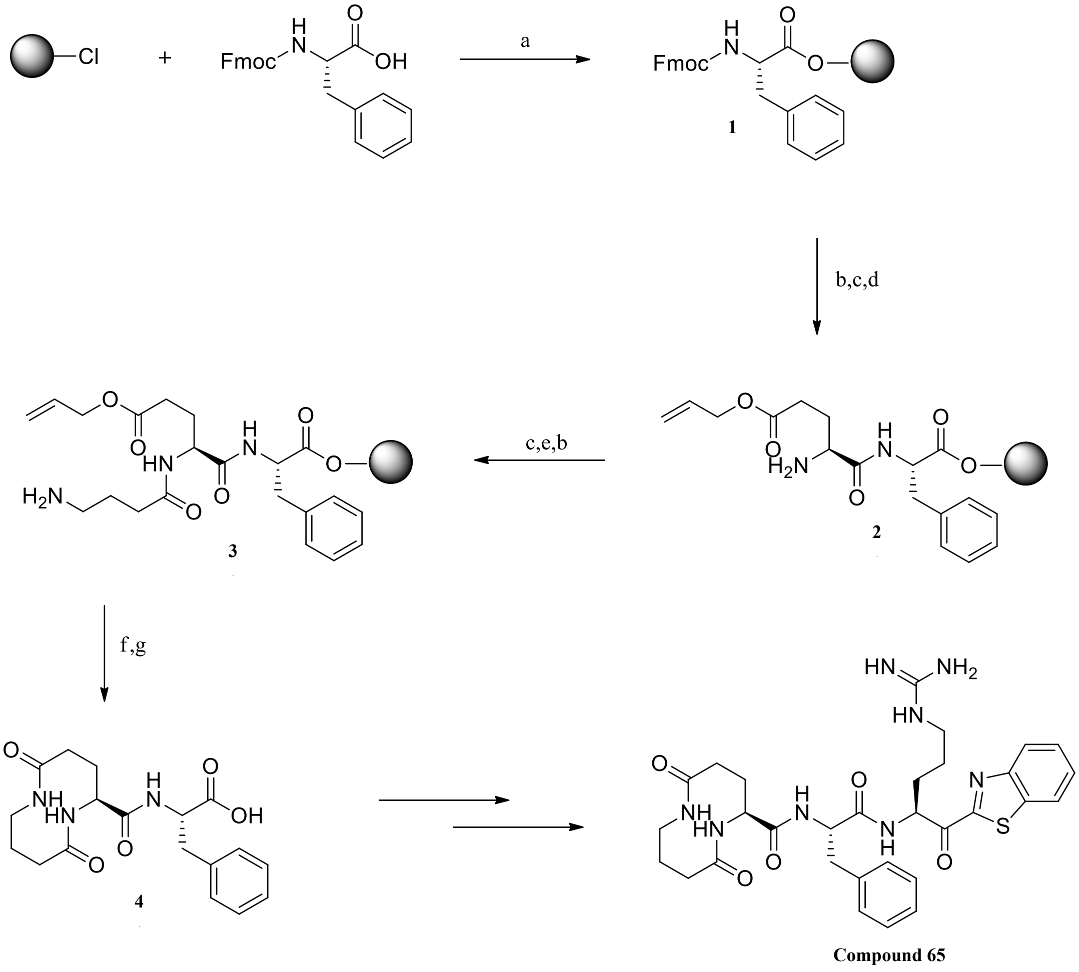

**Figure S28. Solid phase synthesis of [GABA]-Phe (4). Reagents and conditions** : (a) DCM, DIPEA (b) Piperidine/DMF (20:80) (c) Amino acid, HATU, DIPEA, DMF (d) 1-Methylpyrrolidine (25 %), Hexamethylenimine (2 %), HOBt (2 %), NMP/DMSO 1:1, r.t, 1 h (20:80) (e) $PhSiH_3$, $Pd(PPh_3)_4$, $CH_2Cl_2$, r.t, 3 h (f) PyBOP, 6-Cl-HOBt, DIPEA, DMF, r.t, 16 h (g) DCM/HFIP 80:20, r.t, 30 min.

**Compound #66**

*Synthesis of compound #66 (N-0430)*

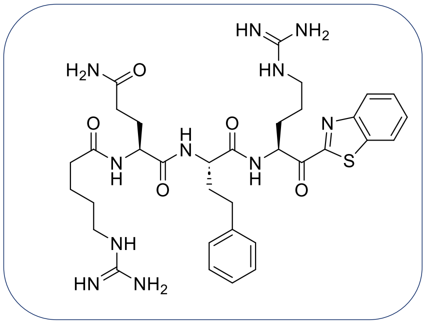

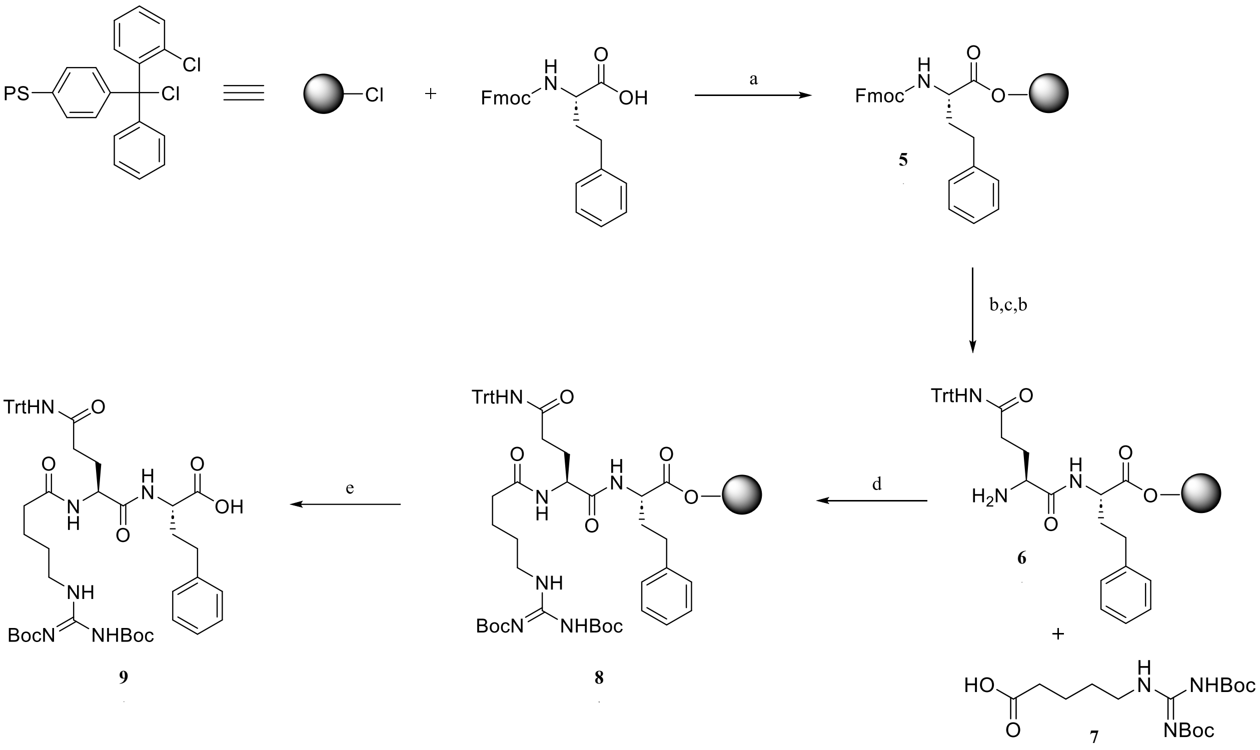

**Figure S29. Solid phase synthesis of (H)Arg(Boc)$_2$-Gln(Trt)-HomoPhe (9). Reagents and conditions**: (a) DCM, DIPEA (b) Piperidine/DMF (20:80) (c) Fmoc-Gln(Trt)-OH, HATU, DIPEA, DMF (d) **7**, HATU, DIPEA, DMF (e) HFIP/DCM (20:80).

**Fmoc-HomoPhe-Resin, Intermediate 5:**

To 500 mg of CTC Resin with a loading 5of 0.5 mmol/g were added Fmoc-HomoPhe-OH (200 mg, 0.5 mmol, 2 eqs.) dissolved in DCM (5 mL per gram of resin), and DIPEA (130 µL, 0.75 mmol, 3 eqs.). The mixture was shaken vigorously for 30-60 min. To endcap any remaining reactive trityl chloride groups, HPLC grade methanol was added (2 mL per gram of resin) and mixed for 15 minutes. The resin was filtered and washed with 3 x DCM, 2 x iPrOH, 2 x DCM, then dried *in vacuo*.

**$NH_2$-Gln(Trt)-HomoPhe-Resin, Intermediate 6:**

A solution of DMF/piperidine (20%) was added to the resin, which was gently shaken for 5 minutes, twice. The resin was filtered and washed with 3 x DMF, iPrOH, 3 x DCM then dried *in vacuo*. A solution of Fmoc-Gln(Trt)-OH (460 mg, 750 µmol, 3 eqs.), HATU (285 mg, 750 µmol, 3 eqs.) and DIPEA (220 µL, 1.25 mmol, 5 eqs.) in DMF (approximatly 10 mL per gram of resin) was added on resin. The mixture was shaken for 2h, filtered, then washed with 3 x DCM, iPrOH, 3 x DCM then dried *in vacuo*.

**(H)Arg$(Boc)_2$-Gln(Trt)-HomoPhe-Resin, Intermediate 8:**

A solution of DMF/piperidine (20%) was added to the resin, which was gently shaken for 5 minutes, twice. The resin was filtered and washed with 3 x DMF, iPrOH, 3 x DCM then dried *in vacuo*. A solution of (H)Arg$(Boc)_2$-OH **7** (285 mg, 625 µmol, 2.5 eqs.), HATU (240 mg, 625 µmol, 2.5 eqs.) and DIPEA (220 µL, 1.25 mmol, 5 eqs.) in DMF (approximatly 10 mL per gram of resin) was added on resin. The resin was shaken for 2h, filtered, then washed with 3 x DCM, iPrOH, 3 x DCM then dried *in vacuo*.

**(H)Arg$(Boc)_2$-Gln(Trt)-HomoPhe-OH, Intermediate 9:**

To 500 mg of derivatized resin was added a solution of 20% HFIP in DCM and shaken twice for 45 minutes. After removal of the solution, the resin was washed with DCM/HFIP (20%) and 3 x DCM. After suspension and co-evaporation in diethylether, the compound was purified by flash chromatography [MeOH/DCM (0.25% AcOH) 0:100 to MeOH/DCM (0.25% AcOH) 10:90] to give tripeptide **9** as a white solid (245 mg, 99%).

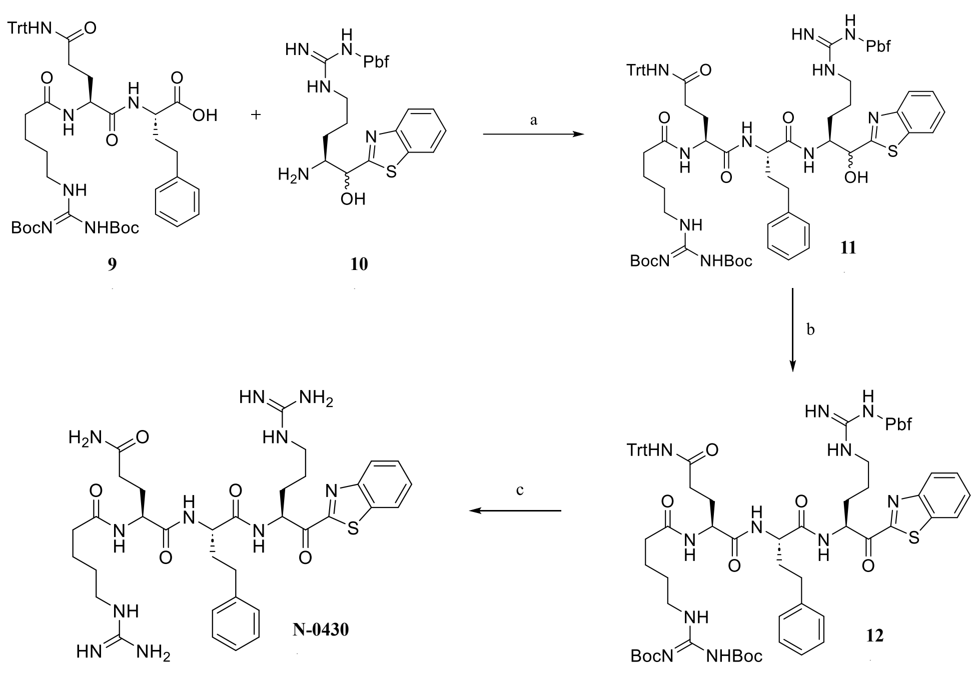

**Figure S30: Solution synthesis of N-0430. Reagents and conditions**: (a) HATU, DIPEA, DMF, 85% (b) DMP, DCM, 96% (c) TFA/$H_2O$/TIPS (95:2.5:2.5).

Warhead synthesis: Compound **10** was prepared as described in *Duchêne et al. 2014*[2] with minor modifications.

To a solution of intermediate **9** (245 mg, 247 µmol, 1 eq.) in anhydrous DMF was added HATU (103 mg, 272 µmol, 1.1 eq.) at 0 °C and the mixture was stirred 5 minutes. $NH_2$-Arg(Pbf)-C(OH)Bt **10** (158 mg, 272 µmol, 1.1 eq.) and DIPEA (216 µL, 1.24 mmol, 5 eq.) were then added and the solution was agitated 15 minutes. The protected tetrapeptide was precipitated on ice, filtrated and washed with cold water twice. The filtrate was dissolved in DCM and washed with brine. The organic phase was dried with sodium sulfate, filtrated and evaporated to give intermediate **11** as a yellow solid. The compound was used in the next step without purification.

DMP (135 mg, 317 µmol, 1.5 eq.) was added to a solution of protected tetrapeptide **11** (600 mg, 211 µmol, 1 eq.) in DCM at 0 °C for 1 hour. The solution was washed with a 10% sodium thiosulfate solution then concentrated. The product was dissolved in ethyl acetate and washed with saturated aqueous sodium bicarbonate and brine. The organic phase was dried with sodium sulfate, filtrated and evaporated. The compound was purified by flash chromatography AcOEt/Hexanes 60:40 to AcOEt/Hexanes 80:20. Intermediate **12** was obtained as a white solid (350 mg, 96%).

350 mg of intermediate **12** was dissolved in a mixture of 1 ml of TFA/$H_2O$/TIPS (95:2.5:2.5) and stirred for 1 hour, until completion of the reaction by UPLC-MS. The TFA/H2O/TIPS solution is added dropwise to 10 mL of cold diethylether (0 °C) in one centrifugation tube and then centrifuged at 4000 rpm for 30 minutes. The supernatant was removed, and the white precipitate was dissolved in water and ACN and lyophilized.

The compound was purified by reverse phase prep-HPLC MS ($C_{18}$) using a ACN/water gradient (0.1% formic acid) from 10-40% of ACN. 20 mg of pure compound was obtained from 100 mg of crude. UPLC-MS retention time : 1.12 min. Purity : 98.2%.

**(H)RQhFR-Kbt (N-0430) :**

$^{1}$H NMR (400 MHz, DMSO) δ 8.81 (d, *J* = 5.8 Hz, 1H), 8.60 (d, *J* = 8.0 Hz, 2H), 8.47 (d, *J* = 7.7 Hz, 1H), 8.43 (s, 1H), 8.28 – 8.22 (m, *J* = 7.7, 4.0, 1.4 Hz, 2H), 7.69 – 7.61 (m, 2H), 7.59 – 7.37 (m, *J* = 23.5 Hz, 8H), 7.27 – 7.20 (m, 2H), 7.18 – 7.12 (m, *J* = 7.2, 5.1 Hz, 3H), 6.73 (s, 1H), 5.47 – 5.37 (m, 1H), 4.26 – 4.15 (m, 2H), 3.14 – 3.00 (m, 4H), 2.66 – 2.55 (m, 1H), 2.25 – 2.02 (m, 4H), 2.01 – 1.75 (m, 6H), 1.73 – 1.53 (m, 4H), 1.47 (dd, *J* = 13.0, 6.3 Hz, 2H).

$^{13}$C NMR (101 MHz, DMSO) δ 192.77, 174.00, 167.19, 164.51, 156.99, 152.94, 141.55, 141.43, 136.37, 128.47, 128.36, 128.25, 128.20, 125.73, 125.27, 123.19, 39.52, 34.95, 34.89, 33.42, 31.94, 31.49, 31.38, 27.87, 24.89, 22.53.
HRMS (*m/z*): $[M+H]^{+}$ calcd for $C_{34}H_{47}N_{11}O_5S$, 361.6814; found, 361.6812.

**Table S6.** Accurate mass measurement for the compound N-0430.

| | |
|---|---|
| Compound | N-0430 Abundant Ion |
| Structure | $C_{34}H_{47}N_{11}O_5S$ |
| Analysis | Qtof |
| Electrospray | $ESI^{+}$ |
| Charge | 2; $[M+2H]^{2+}$ |
| m/z theorical | 361.6814 |
| m/z measured | 361.6812 |
| Δm | 0. 2 |
| Dissolution solvent | Methanol |

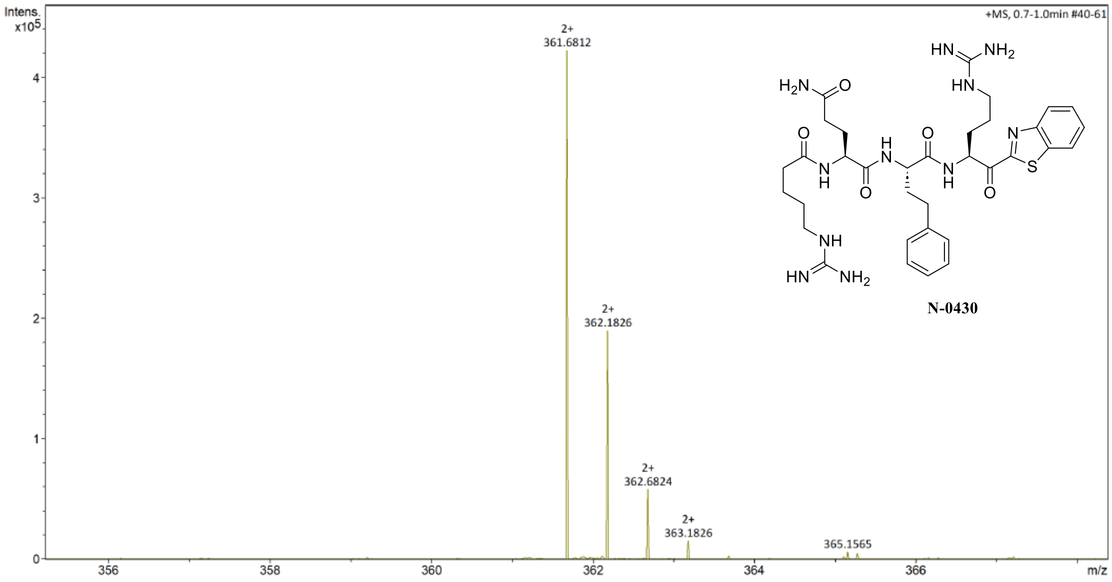

**Figure S31.** Isotopic profile for the most abundant ion (double charged) N-0430, $[M+H]^{2+}$ detected with high-resolution mass spectrometer (Qtof).

**Compound #67**

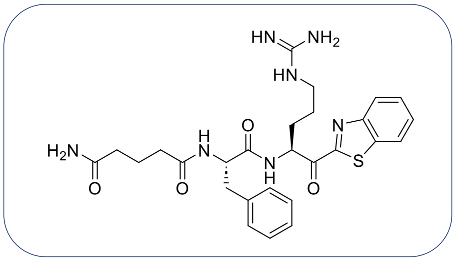

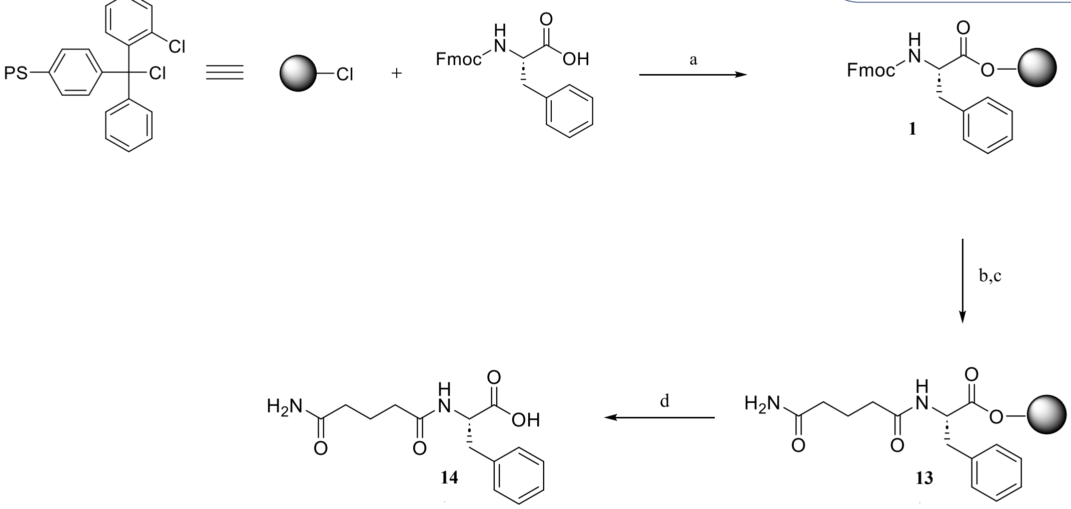

**Figure S32. Solid phase synthesis of (H)Gln-Phe (14). Reagents and conditions**: (a) DCM, DIPEA (b) Piperidine/DMF (20:80) (c) 5-Amino-5-oxopentanoic acid, HATU, DIPEA, DMF (d) HFIP/DCM (20:80).

**Fmoc-Phe-Resin, Intermediate 1:**

To 500 mg of CTC Resin with a loading of 0.3 mmol/g were added Fmoc-Phe-OH (116 mg, 0.3 mmol, 2 eqs.) dissolved in DCM (5 mL per gram of resin), and DIPEA (80 µL, 0.45 mmol, 3 eqs.). The mixture was shaken vigorously for 30-60 min. To endcap any remaining reactive trityl chloride groups, HPLC grade methanol was added (2 mL per gram of resin) and mixed for 15 minutes. The resin was filtered and washed with 3 x DCM, 2 x iPrOH, 2 x DCM, then dried *in vacuo*.

**(H)Gln-Phe-Resin, Intermediate 13:**

A solution of DMF/piperidine (20%) was added to the resin, which was gently shaken for 5 minutes, twice. The resin was filtered and washed with 3 x DMF, iPrOH, 3 x DCM then dried *in vacuo*. A solution of 5-Amino-5-oxopentanoic acid (60 mg, 450 µmol, 3 eqs.), HATU (170 mg, 450 µmol, 3 eqs.) and DIPEA (130 µL, 750 mmol, 5 eqs.) in DMF (approximatly 10 mL per gram of resin) was added on resin. The mixture was shaken for 2h, filtered, then washed with 3 x DCM, iPrOH, 3 x DCM then dried *in vacuo*.

**(H)Gln-Phe-OH, Intermediate 14:**

To 500 mg of derivatized resin was added a solution of 20% HFIP in DCM and shaken twice for 45 minutes. After removal of the solution, the resin was washed with DCM/HFIP (20%) and 3 x DCM. After suspension and co-evaporation in diethylether, the product was used in the next step whitout purification (25 mg, 43%).

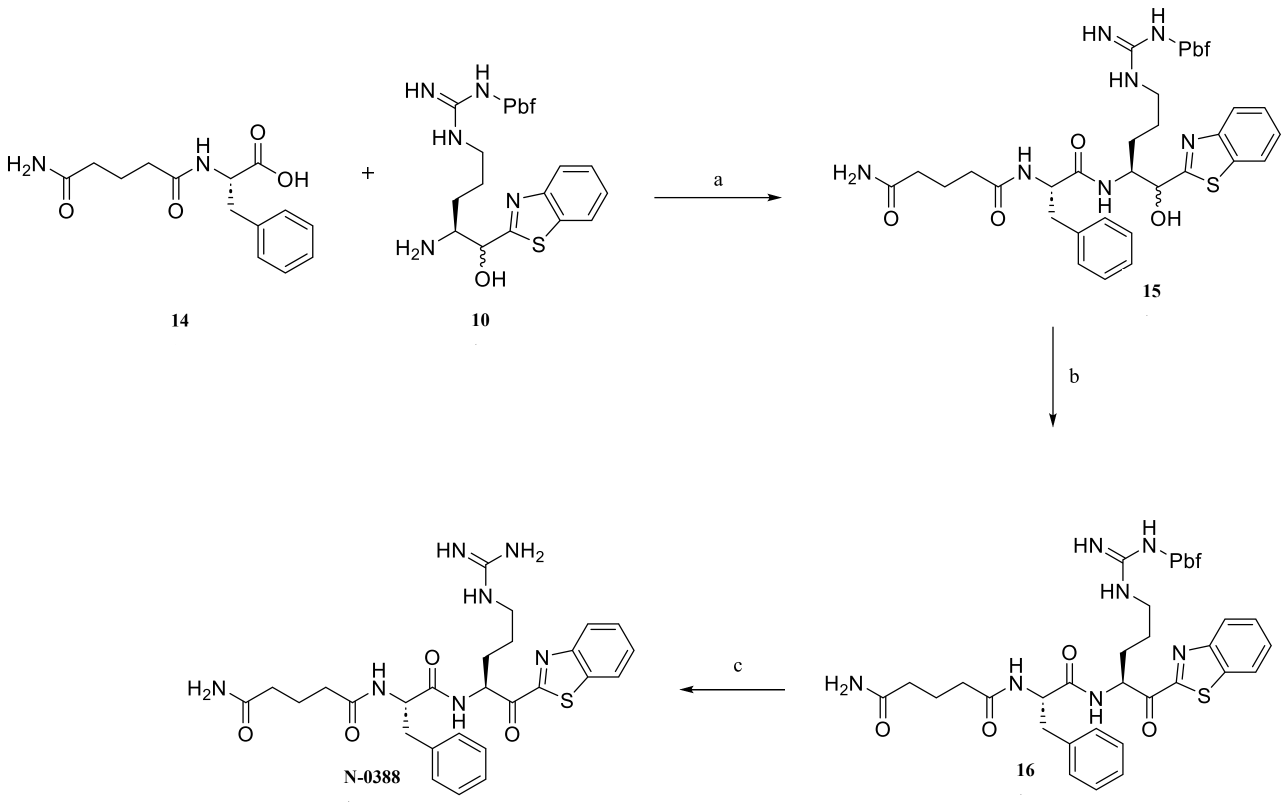

**Figure S33. Solution synthesis of N-0388. Reagents and conditions**: (a) HATU, DIPEA, DMF, 99% (b) DMP, DCM, 99% (c) TFA/$H_2O$/TIPS (95:2.5:2.5).

Warhead synthesis: Compound **10** was prepared as described in *Duchêne et al. 2014*[2] with minor modifications.

To a solution of intermediate **14** (25 mg, 65 µmol, 1 eq.) in anhydrous DMF was added HATU (27 mg, 71 µmol, 1.1 eqs.) at 0 °C and the mixture was stirred 5 minutes. $NH_2$-Arg(Pbf)-C(OH)Bt **10** (42 mg, 71 µmol, 1.1 eqs.) and DIPEA (34 µL, 195 mmol, 3 eqs.) were then added and the solution was agitated 15 minutes. The protected tetrapeptide was precipitated on ice, filtrated and washed with cold water twice. The filtrate was dissolved in DCM and washed with brine. The organic phase was dried with sodium sulfate, filtrated and evaporated. The yellow solid was used in the next step whitout purification.

DMP (41 mg, 96 µmol, 1.5 eqs.) was added to a solution of protected tripeptide **15** (70 mg, 64 µmol, 1 eq.) in DCM at 0 °C for 1 hour. The solution was washed with a 10% sodium thiosulfate solution then concentrated. The product was dissolved in ethyl acetate and washed with saturated aqueous sodium bicarbonate and brine. The organic phase was dried with sodium sulfate, filtrated and evaporated. The compound was purified by flash chromatography MeOH/DCM 0:100 to MeOH/DCM 10:90. Intermediate **16** was obtained as a white solid (60 mg, 99%).

60 mg of intermediate **16** was dissolved in a mixture of 1 ml of TFA/$H_2O$/TIPS (95:2.5:2.5) and stirred for 1 hour, until completion of the reaction by UPLC-MS. The TFA/H2O/TIPS solution is added dropwise to 10 mL of cold diethylether (0 °C) in one centrifugation tube and then

centrifuged at 4000 rpm for 30 minutes. The supernatant was removed and the white precipitate was dissolved in water and ACN and lyophilized.

The compound was purified by reverse phase prep-HPLC MS ($C_{18}$) using a ACN/water gradient (0.1% formic acid) from 10-40% of ACN. 10 mg of pure compound was obtained from 25 mg of crude. UPLC-MS retention time : 1.19 min. Purity : 97.7 %.

**(H)QFR-Kbt (N-0388) :**

1H NMR (400 MHz, DMSO) δ 8.95 (d, J = 6.1 Hz, 2H), 8.44 (s, 1H), 8.28 (dd, J = 6.6, 2.8 Hz, 1H), 8.12 (d, J = 8.3 Hz, 1H), 7.85 (s, 3H), 7.71 – 7.62 (m, 2H), 7.23 (d, J = 4.3 Hz, 3H), 7.17 (ddd, J = 13.0, 7.6, 4.7 Hz, 2H), 6.69 (s, 1H), 6.32 (s, 1H), 5.49 – 5.35 (m, 1H), 4.70 – 4.56 (m, J = 10.4, 8.8, 4.9 Hz, 1H), 3.06 (dd, J = 26.1, 4.8 Hz, 2H), 3.01 – 2.66 (m, 2H), 2.10 – 1.91 (m, 4H), 1.91 – 1.69 (m, 2H), 1.68 – 1.44 (m, 4H).

13C NMR (101 MHz, DMSO) δ 193.08, 173.99, 172.02, 171.75, 167.23, 164.51, 157.50, 152.97, 137.95, 137.74, 136.39, 129.14, 129.07, 128.22, 127.99, 127.56, 125.30, 123.21, 54.65, 53.39, 39.52, 34.55, 34.48, 34.44, 27.55, 24.97, 21.25.

HRMS (m/z): $[M+H]^+$ calcd for $C_{27}H_{33}N_7O_4S$, 552.2388; found, 552.2392.

**Table S7.** Accurate mass measurement for the compound N-0388.

| | |
|---|---|
| Compound | N-0388 Abundant Ion |
| Structure | $C_{27}H_{33}N_7O_4S$ |
| Analysis | Q-TOF (maXis) |
| Electrospray | $ESI^+$ |
| Charge | $[M+H]^+$ |
| m/z theorical | 552.2388 |
| m/z measured | 552.2392 |
| Δm | 0.4 |
| Dissolution solvent | MeOH |

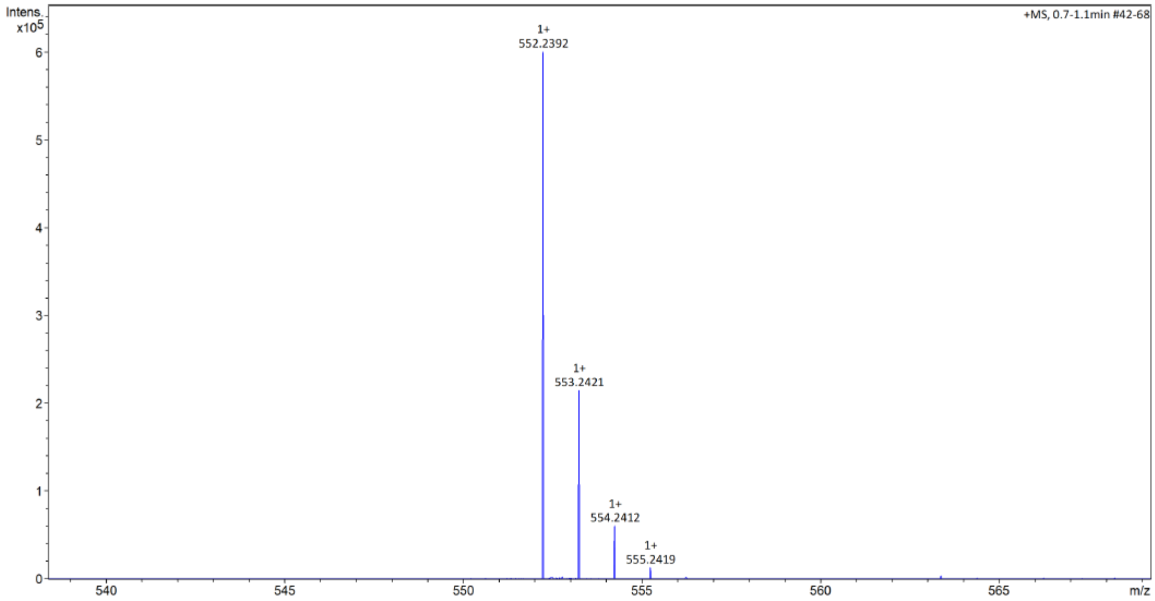

**Figure S34.** Isotopic profile for the most abundant ion N-0388, $[M+H]^+$ detected with high-resolution mass spectrometer (Qtof).

**Compound #68**

*Synthesis of N-0430(OH)*

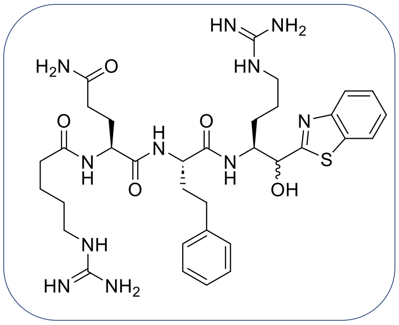

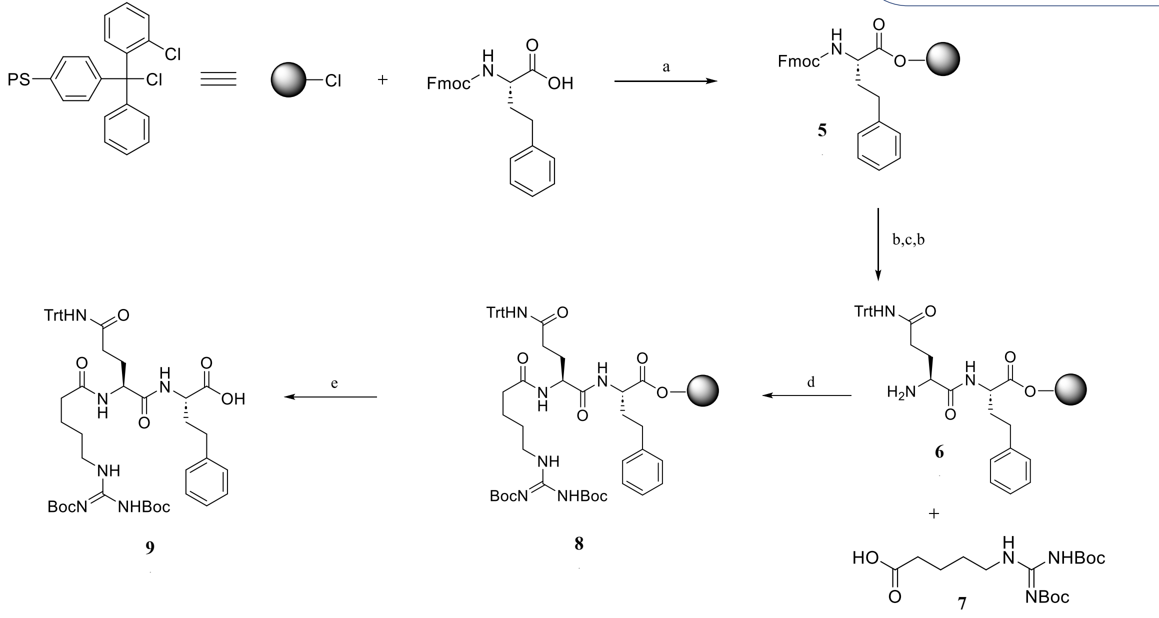

**Figure S35. Solid phase synthesis of (H)Arg(Boc)$_2$-Gln(Trt)-HomoPhe (9)**. Reagents and conditions: (a) DCM, DIPEA (b) Piperidine/DMF (20:80) (c) Fmoc-Gln(Trt)-OH, HATU, DIPEA, DMF (d) **7**, HATU, DIPEA, DMF (e) HFIP/DCM (20:80).

**Fmoc-HomoPhe-Resin, Intermediate 5:**

To 500 mg of CTC Resin with a loading of 0.5 mmol/g were added Fmoc-HomoPhe-OH (200 mg, 0.5 mmol, 2 eqs.) dissolved in DCM (5 mL per gram of resin), and DIPEA (130 µL, 0.75 mmol, 3 eqs.). The mixture was shaken vigorously for 30-60 min. To endcap any remaining reactive trityl chloride groups, HPLC grade methanol was added (2 mL per gram of resin) and mixed for 15 minutes. The resin was filtered and washed with 3 x DCM, 2 x iPrOH, 2 x DCM, then dried *in vacuo*.

**NH$_2$-Gln(Trt)-HomoPhe-Resin, Intermediate 6:**

A solution of DMF/piperidine (20%) was added to the resin, which was gently shaken for 5 minutes, twice. The resin was filtered and washed with 3 x DMF, iPrOH, 3 x DCM then dried *in vacuo*. A solution of Fmoc-Gln(Trt)-OH (460 mg, 750 µmol, 3 eqs.), HATU (285 mg, 750 µmol, 3 eqs.) and DIPEA (220 µL, 1.25 mmol, 5 eqs.) in DMF (approximatly 10 mL per gram of resin) was added on resin. The mixture was shaken for 2h, filtered, then washed with 3 x DCM, iPrOH, 3 x DCM then dried *in vacuo*.

**(H)Arg(Boc)$_2$-Gln(Trt)-HomoPhe-Resin, Intermediate 8:**

A solution of DMF/piperidine (20%) was added to the resin, which was gently shaken for 5 minutes, twice. The resin was filtered and washed with 3 x DMF, iPrOH, 3 x DCM then dried *in vacuo*. A solution of (H)Arg(Boc)$_2$-OH **7** (250 mg, 640 µmol, 2.5 eqs.), HATU (240 mg, 635 µmol, 2.5 eqs.) and DIPEA (220 µL, 1.25 mmol, 5 eqs.) in DMF (approximatly 10 mL per gram of resin) was added on resin. The resin was shaken for 2h, filtered, then washed with 3 x DCM, iPrOH, 3 x DCM then dried *in vacuo*.

**(H)Arg(Boc)$_2$-Gln(Trt)-HomoPhe-OH, Intermediate 9:**

To 500 mg of derivatized resin was added a solution of 20% HFIP in DCM and shaken twice for 45 minutes. After removal of the solution, the resin was washed with DCM/HFIP (20%) and 3 x DCM. After suspension and co-evaporation in diethylether, the compound was purified by flash chromatography [MeOH/DCM (0.25% AcOH) 0:100 to MeOH/DCM (0.25% AcOH) 10:90] to give the desired intermediate **9** as a white solid (200 mg, 85%).

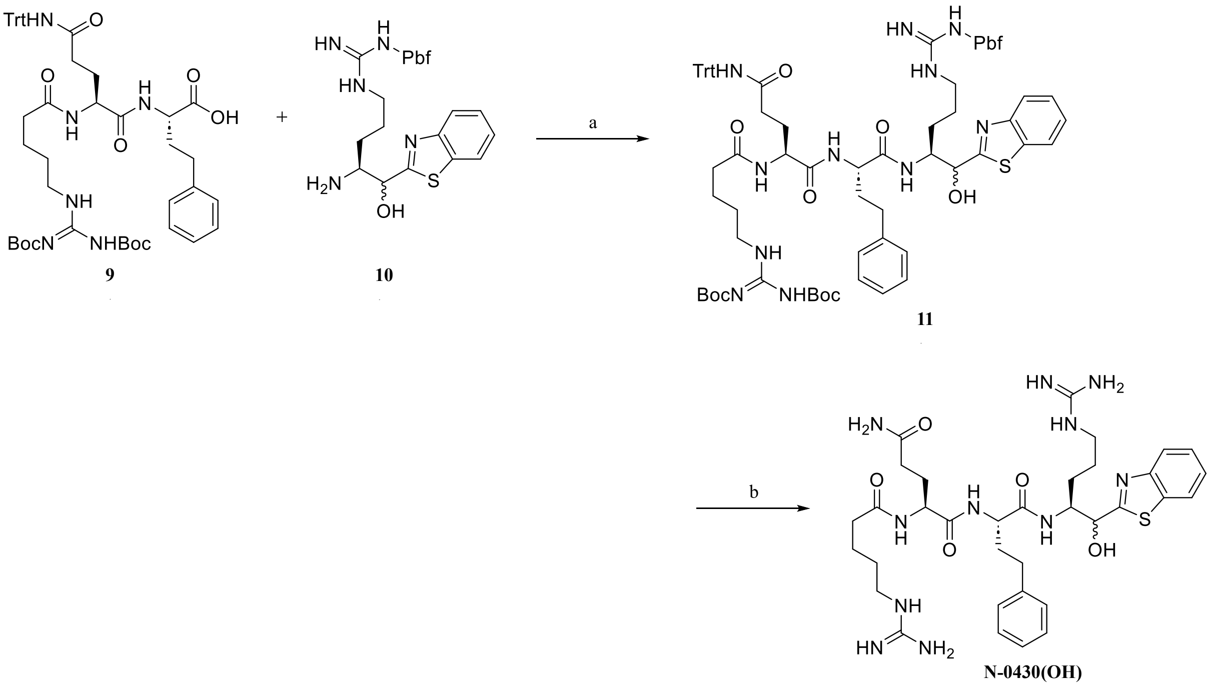

**Figure S36. Solution synthesis of N-0430(OH). Reagents and conditions**: (a) HATU, DIPEA, DMF, 85% (b) TFA/$H_2O$/TIPS (95:2.5:2.5).

Warhead synthesis: Compound **10** was prepared as described in *Duchêne et al. 2014*[2] with minor modifications.

To a solution of intermediate **9** (200 mg, 213 µmol, 1 eq.) in anhydrous DMF was added HATU (90 mg, 235 µmol, 1.1 eq.) at 0 °C and the mixture was stirred 5 minutes. $NH_2$-Arg(Pbf)-C(OH)Bt **10** (137 mg, 235 µmol, 1.1 eq.) and DIPEA (185 µL, 1.07 mmol, 5 eq.) were then added and the solution was agitated 15 minutes. The protected tetrapeptide was precipitated on ice, filtrated and washed with cold water twice. The filtrate was dissolved in DCM and washed with brine. The organic phase was dried with sodium sulfate, filtrated and evaporated. The yellow solid was used in the next step without purification.

400 mg of intermediate **11** was dissolved in a mixture of 1 ml of TFA/$H_2O$/TIPS (95:2.5:2.5) and stirred for 1 hour, until completion of the reaction by UPLC-MS. The TFA/H2O/TIPS solution is added dropwise to 10 mL of cold diethylether (0 °C) in one centrifugation tube and then centrifuged at 4000 rpm for 30 minutes. The supernatant was removed, and the white precipitate was dissolved in water and ACN and lyophilized.

The compound was purified by reverse phase prep-HPLC MS ($C_{18}$) using a ACN/water gradient (0.1% formic acid) from 10-40% of ACN. 45 mg (30 mg of the first dia and 15 mg of th second dia) of pure compound was obtained from 150 mg of crude. UPLC-MS retention time : 1.10 min and 1.12 min. Purity : 99.5 %.

**(H)RQhFR-(OH)Kbt (N-0430-OH) :**

**Dia 1 :**

$^1$H NMR (400 MHz, DMSO) δ 8.48 (s, 1H), 8.38 (dd, *J* = 19.8, 7.7 Hz, 2H), 8.27 (s, 1H), 8.18 (d, *J* = 8.8 Hz, 1H), 8.06 – 8.00 (m, 1H), 7.78 (dd, *J* = 7.1, 1.7 Hz, 1H), 7.56 (s, *J* = 31.0 Hz, 8H), 7.38 (tdd, *J* = 8.7, 7.3, 1.5 Hz, 2H), 7.25 (t, *J* = 7.3 Hz, 2H), 7.16 (t, *J* = 7.3 Hz, 1H), 7.11 (d, *J* = 7.0 Hz, 2H), 6.78 (s, 1H), 4.84 (d, *J* = 4.8 Hz, 1H), 4.33 – 4.08 (m, 3H), 3.06 (d, *J* = 5.3 Hz, 2H), 3.02 – 2.84 (m, *J* = 20.1, 15.2 Hz, 2H), 2.57 (dd, *J* = 19.1, 9.6 Hz, 1H), 2.47 – 2.37 (m, 1H), 2.28 – 2.12 (m, 3H), 2.11 – 2.02 (m, 1H), 1.97 – 1.84 (m, 2H), 1.76 (ddd, *J* = 27.9, 17.3, 9.8 Hz, 2H), 1.66 – 1.32 (m, 8H).

13C NMR (101 MHz, DMSO) δ 176.23, 174.24, 172.57, 172.00, 171.57, 167.49, 157.10, 153.11, 141.58, 134.41, 128.41, 128.26, 125.81, 125.74, 124.68, 122.50, 122.14, 73.32, 53.29, 52.97, 52.88, 39.52, 34.87, 33.63, 31.97, 31.66, 27.89, 25.65, 25.03, 22.53.

**Dia 2 :**

$^1$H NMR (400 MHz, DMSO) δ 8.49 (s, 1H), 8.40 (d, *J* = 7.8 Hz, 1H), 8.35 (s, 1H), 8.20 (d, *J* = 7.2 Hz, 1H), 7.92 (d, *J* = 7.7 Hz, 1H), 7.86 (d, *J* = 9.1 Hz, 1H), 7.83 (d, *J* = 8.1 Hz, 1H), 7.54 (s, 5H), 7.49 (s, 2H), 7.44 – 7.39 (m, 1H), 7.35 – 7.30 (m, 1H), 7.19 (t, *J* = 7.2 Hz, 2H), 7.15 – 7.10 (m, 1H), 6.92 (d, *J* = 7.0 Hz, 2H), 6.74 (s, 1H), 5.00 (s, 1H), 4.34 – 4.24 (m, 1H), 4.16 (ddd, *J* = 15.7, 13.7, 8.2 Hz, 2H), 3.05 (d, *J* = 5.0 Hz, 4H), 2.40 – 2.26 (m, 2H), 2.25 – 2.10 (m, 3H), 2.08 – 1.99 (m, 1H), 1.96 – 1.86 (m, 1H), 1.83 – 1.62 (m, 4H), 1.53 (dd, *J* = 20.6, 15.4 Hz, 5H), 1.47 – 1.37 (m, 2H).

13C NMR (101 MHz, DMSO) δ 176.49, 174.20, 172.45, 171.90, 171.45, 167.35, 157.08, 153.07, 141.46, 134.42, 128.17, 128.13, 125.70, 125.61, 124.47, 122.38, 122.03, 72.39, 53.07, 52.84, 52.58, 39.52, 34.87, 33.40, 32.07, 31.21, 28.01, 27.83, 25.12, 22.54.

HRMS (m/z): $[M+H]^+$ calcd for $C_{34}H_{49}N_{11}O_5S$, 362.6892; found, 362.6891.

**Dia 1**

HRMS (m/z): $[M+H]^+$ calcd for $C_{34}H_{49}N_{11}O_5S$, 362.6892; found, 362.6891.

**Table S8.** Accurate mass measurement for the compound N-0430-OH (1).

| | |
|---|---|
| Compound | N-0430-OH Abundant Ion |
| Structure | $C_{34}H_{49}N_{11}O_5S$ |
| Analysis | Q-TOF (maXis) |
| Electrospray | $ESI^+$ |
| Charge | 2; $[M+2H]^{2+}$ |
| m/z theorical | 362.6892 |
| m/z measured | 362.6891 |
| Δm | 0.1 |
| Dissolution solvent | MeOH |

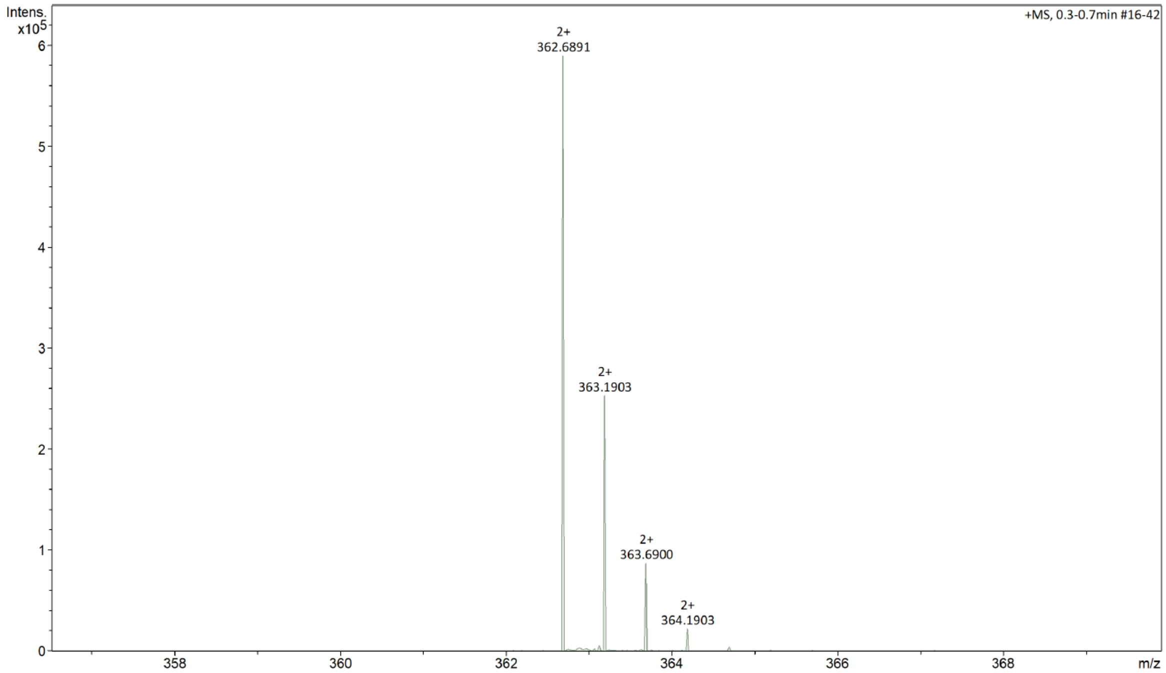

**Figure S37.** Isotopic profile for the most abundant ion (double charged) N-0430-OH (dia 1), $[M+H]^{2+}$ detected with high-resolution mass spectrometer.

**Dia 2**

HRMS (m/z): $[M+H]^+$ calcd for $C_{34}H_{49}N_{11}O_5S$, 362.6892; found, 362.6880.

**Table S9.** Accurate mass measurement for the compound N-0430-OH (2).

| | |
|---|---|
| Compound | N-0430-OH Abundant Ion |
| Structure | $C_{34}H_{49}N_{11}O_5S$ |
| Analysis | Q-TOF (maXis) |
| Electrospray | $ESI^+$ |
| Charge | 2; $[M+2H]^{2+}$ |
| m/z theorical | 362.6892 |
| m/z measured | 362.6880 |
| Δm | 1.2 |
| Dissolution solvent | MeOH |

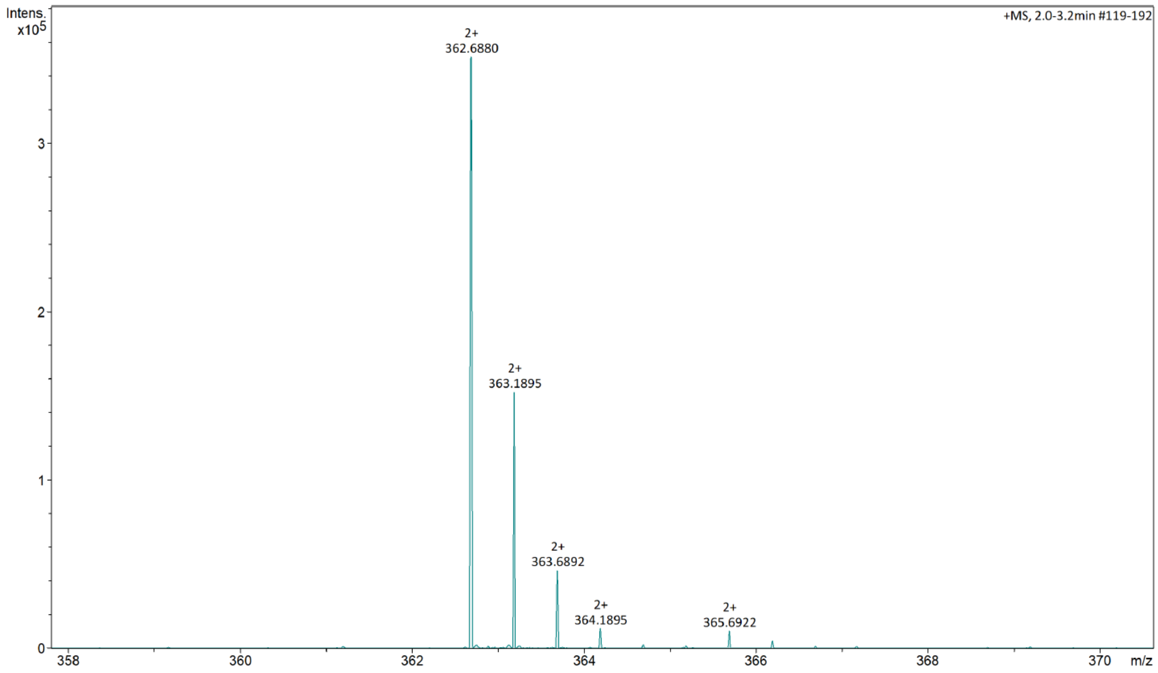

**Figure S38.** Isotopic profile for the most abundant ion N-0430-OH (dia 2), $[M+H]^{2+}$ detected with high-resolution mass spectrometer.
